# Postdoc X-ray in Europe 2017 Work conditions, productivity, institutional support and career outlooks

**DOI:** 10.1101/523621

**Authors:** Maria José Ribeiro, Ana Fonseca, Mariana Moura Ramos, Marta Costa, Konstantina Kilteni, Lau Møller Andersen, Lisa Harber-Aschan, Joana A. Moscoso, Sonchita Bagchi, European Network of Postdoctoral Associations

## Abstract

1.

This survey and data analysis were conducted by the European Network of Postdoctoral Associations (ENPA) with the aim of assessing the current research and work conditions, aspirations and support received by postdoctoral researchers working in Europe.

The Results section is structured into three main parts. The first one describes the study sample of European postdoctoral researchers, including participants’ demographics, funding sources and income, research outputs, and teaching opportunities. The second section focuses on their professional aspirations and institutional support provided. The third part describes the level of engagement of postdoctoral researchers and their institutions in working towards better research conditions and career development, and what initiatives are emerging within this community.

Our Conclusions section pulls together this comprehensive analysis, highlighting some of the most concerning issues currently affecting postdoctoral researchers in Europe. We also make a number of recommendations that would significantly improve the career expectations and aspirations of postdoctoral researchers. These are listed below.

**Conclusions and recommendations:**

1. Longer postdoctoral periods in Southern Europe despite higher publication metrics **Recommendation** Institutions in Southern Europe should develop clear criteria to support postdoctoral researchers’ career progression.
2. Southern and Eastern Europe pay the lowest salaries and have the lowest number of foreign postdoctoral researchers **Recommendation** The salary differences across European countries should be addressed as this could be a barrier to mobility and knowledge exchange from higher to lower pay regions.
3. Lack of access to funding is a significant concern of postdoctoral researchers **Recommendation** Discrepancies in access to funding should be minimized across the different European areas.
4. Postdoctoral researchers in Europe work longer hours than required by contract **Recommendation** The culture of overwork in the research environment should be addressed in order to protect researchers against the risks associated with long hours at work.
5. The majority of full-time postdoctoral work contracts includes an exclusivity clause **Recommendation** Inclusion of exclusivity clauses in contracts for postdoctoral researchers should be optional in order to allow them to enhance their employability outside academia.
6. Postdoctoral researchers’ career development is poorly supported by their institutions **Recommendation** Postdoctoral researchers’ career prospects and career management should be much more supported by institutions in coordination with postdoctoral associations.
7. Lack of postdoctoral representation in governance is linked to unclear institutional duties and rights **Recommendation** Institutional governance bodies should include postdoctoral researcher representatives. This would ensure that the views of this vital staff group are heard, as well as making postdoctoral researchers feel more engaged with their own institutions. A flexible and proactive communication strategy at the institution and research group level should be developed, taking into account the sometimes transient nature of postdoctoral researchers’ posts.
8. Researchers show higher engagement with their local postdoctoral associations than with workers’ unions **Recommendation** Postdoctoral associations are an essential way to advocate for postdoctoral researchers at the governance level. Institutions should engage with, promote and support the work of postdoctoral associations.

## 2. Introduction

### 2.1 Why we did it

In an article published in 2007, Professor Åkerlind identified the basic issues regarding postdoctoral positions <u>(Åkerlind * 2005)</u>. These included the absence of a systematic definition of postdoctoral researchers as well as lack of structure, policy and data regarding postdoctoral appointments. After 11 years, these issues are still the dominating factors, contributing to dissatisfaction and loss of motivation among postdoctoral communities. Based on two reports, one from the UK and the other from the US, Nature published an article in 2014 that described **research as ‘a brutal business’** <u>(“Harsh Reality” 2014)</u>. The article mentioned that, there are too many talented bright young researchers chasing too few secured academic careers. Postdoctoral researchers often describe themselves as ‘lost’ or ‘invisible’ while working under tremendous pressure in an environment with no job security. A recent comparative study of two Dutch universities <u>(van der Weijden et al. 2016)</u> revealed that 85 % of 225 respondents of these studies wanted to stay in academia, but less than 3% was offered a tenure-track position. The authors identified 3 major problems, 1) limited knowledge about the growing population of postdoctoral researchers; 2) postdoctoral researchers are often not recognized as staff at many European universities; 3) the uncertainty of postdoctoral researchers career prospects. This consequently raises the issue of insecurity and uncertainty in their career progression. A similar result was reported recently in 2017 from the Max Planck Society in Germany <u>(PhDnet 2018)</u>.

To increase the knowledge about postdoctoral researchers (who they are, what they do and what their career prospects are) and to further contribute towards the above mentioned discussion in a constructive way, in the second half of 2017 the **European Network of Postdoctoral Associations (ENPA, https://www.uc.pt/en/iii/postdoc/ENPA)** conducted a survey to evaluate working conditions, training opportunities and career perspectives of postdoctoral researchers across Europe. The survey questions can be found in the Supplementary Material and in the ENPA website. The analysis of this survey is presented in this report.

### 2.2 Who we are

The **European Network of Postdoctoral Associations (ENPA)** was formed in 2016 with the ambition to bring together European postdoctoral associations as well as individual postdoctoral scientists currently working in Europe.

ENPA aims to:

1. Represent postdoctoral scientists working in Europe, regardless of their nationality;
2. Advocate for the improvement of current working and training conditions;
3. Advocate for better opportunities for career development and progression;
4. Describe, discuss and compare the organization, objectives, and activities of existing post-doctoral associations/movements;
5. Describe, discuss and compare the integration and career perspectives of postdoctoral researchers and their opinions on science policies at the local and national level. This analysis is based on information we collect through surveys completed by the postdoctoral community;
6. Draft proposal of the best practices regarding the management of postdoctoral researchers to bring to the attention of decision makers.

The postdoctoral researchers that put together the survey and did the analysis are presented at the end of this report.

## 3. Results

In total, **898 postdoctoral researchers** working in European universities and research institutes answered the survey. The respondents belonged to 241 different institutions (see Annex 1) in 27 European countries (see Annex 2). We analyzed the survey data with the aim of comparing responses across different European regions, gender, and research areas. Furthermore, where relevant, we also analyzed the effect of mobility (i.e., whether the researcher was working in his/her home country or not; see Annex 3 for nationality).

To study **differences across Europe**, we divided the responses according to European region (Western Europe, Southern Europe and Eastern Europe - as detailed in Annex 2). About half (54%) of the responses were from postdoctoral researchers working in Western Europe, 42% from Southern Europe and 4% from Eastern Europe. A limitation of this survey is therefore the small number of responses from researchers working in Eastern Europe.

To study **differences across research areas**, we divided the primary research areas in five main research area groups: Social Sciences & Humanities & Economics (SocSci&Hum&Econ); Life Sciences (LifeSci); Environment & Geosciences (Env&Geo); Chemistry & Physics & Maths (Chem&Phys&Maths); and Information Sciences & Engineering (IT&Eng) (see Annex 4 for grouping details). The majority of the survey responses came from researchers working in LifeSci (55%), followed by SocSci&Hum&Econ (16%), Chem&Phys&Maths (13%), Env&Geo (9%) and finally IT&Eng with the lowest number of responses (7%).

Statistical analyses of categorical variables were conducted with chi-square tests. Statistical comparisons of numerical variables were conducted with Kruskal-Wallis test (assuming non-normality of the data).

### 3.1 Postdoctoral researchers: Who they are and what they do?

#### 3.1.1 Demographics

##### Q3-4 Gender and Age

We had a **higher number of responses from women** (61%) than men (Fig. 1A). The gender ratio was similar across the three European regions studied and did not differ significantly (*p* =.48, Fig. 1B). Yet, gender ratio was significantly different when comparing across different research areas (*p* =.001, Fig. 1C). We found the highest percentage of women in the research area group SocSci&Hum&Econ and the lowest in IT&Eng.

**Figure 1.**
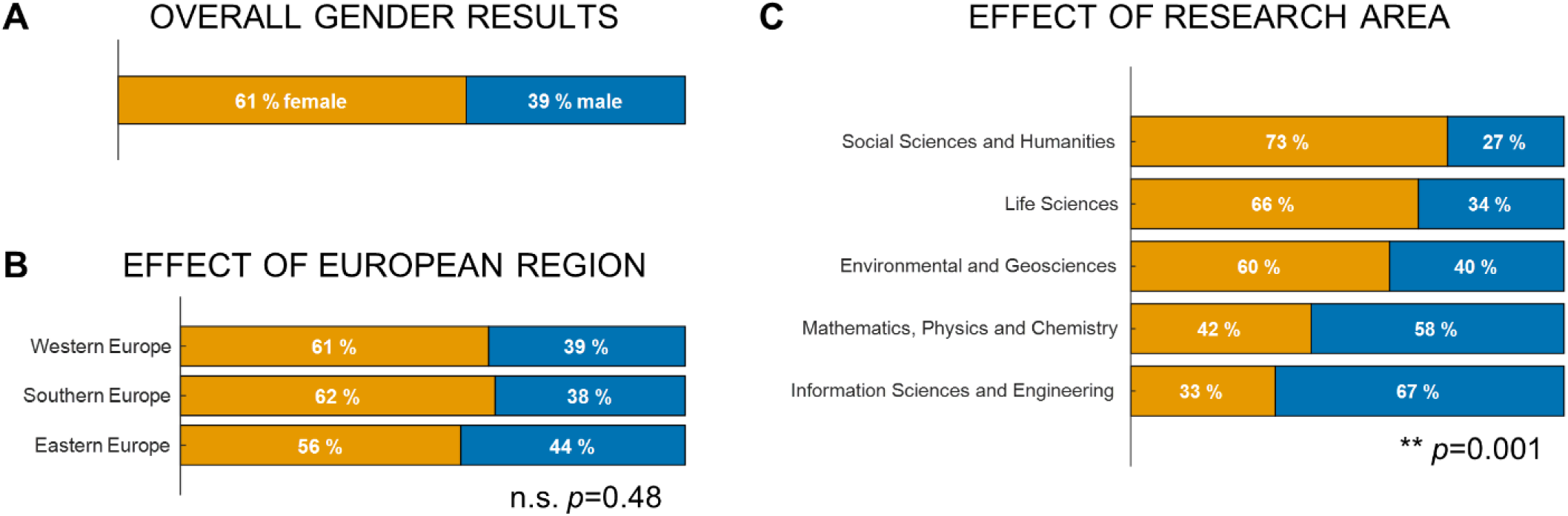
Gender ratio of postdoctoral researchers working in Europe.

The majority of postdoctoral researchers surveyed (78%) were **aged between 30 and 40 years** (median age = 34 years; Fig. 2A). There was no significant age difference across gender (*p* =.1; Fig. 2B). Researchers working in their home country were on average older than researchers working abroad (*p* <.001; median age in years: home country = 35, abroad = 33; Fig. 2C). Moreover, there was also a significant effect of European region, with postdoctoral researchers working in Southern Europe being older than their counterparts in Western and Eastern Europe (*p* <.001; median age in years: Western = 33; Southern = 36; Eastern = 33 - Figure 2D). The age of postdoctoral researchers also varied with research area (*p* <.001; median age in years: SocSci&Hum&Econ = 35; LifeSci = 34; Env&Geo = 36; Chem&Phys&Maths = 33; IT&Eng = 33 – Fig. 2E).

**Figure 2.**
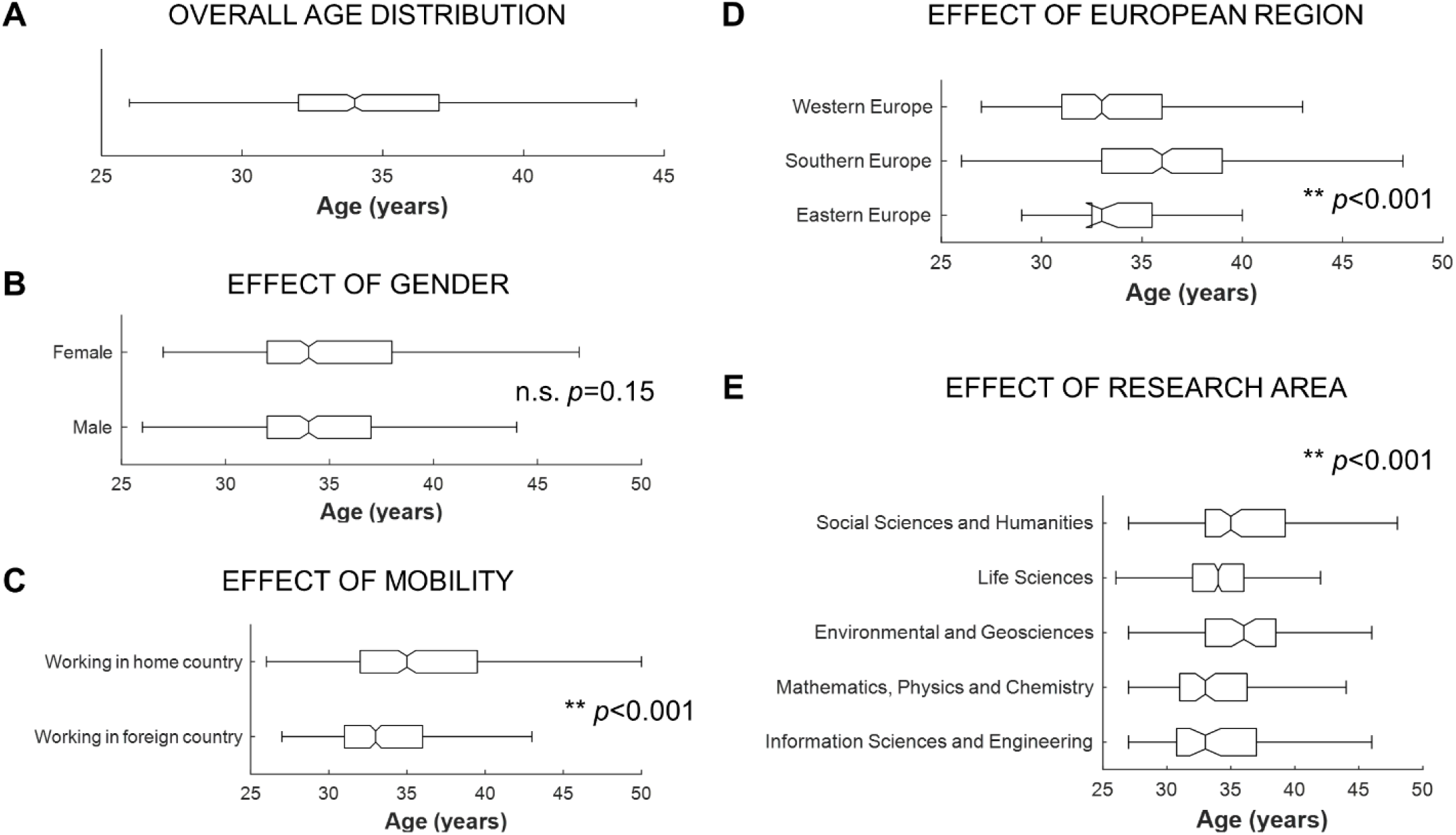
Age of postdoctoral researchers working in Europe.

One explanation for the differences in age observed is that **the age at which researchers finish their PhD is different across European regions and across research areas**. To test this, we compared the age at PhD conclusion across these variables. This was estimated based on the reported researchers’ age and year of PhD conclusion. Results are shown in Figure 3.

**Figure 3.**
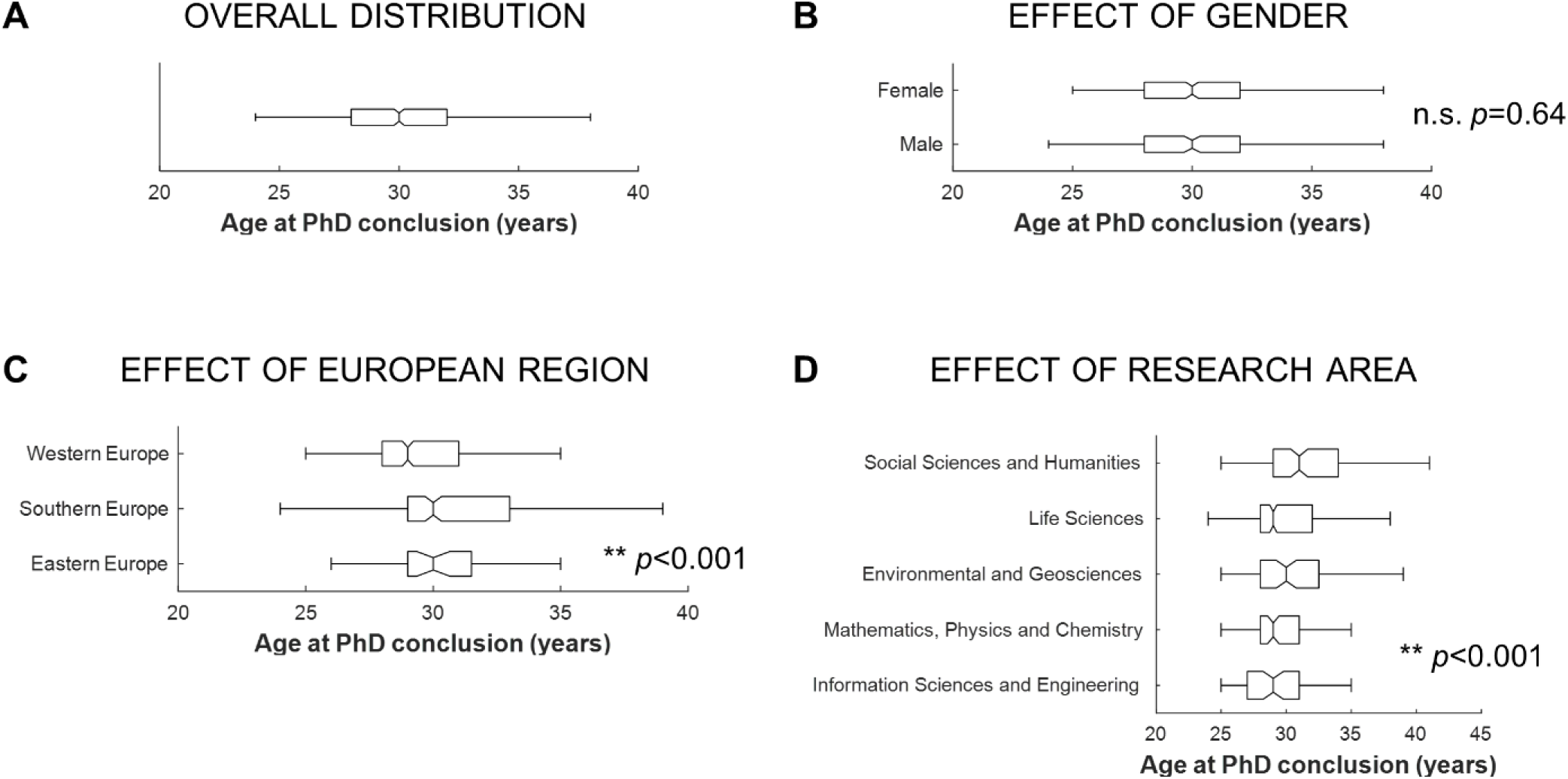
Estimated age at which postdoctoral researchers working in Europe concluded their PhD.

Researchers currently working in Southern and Eastern Europe were awarded their PhD at an older age than researchers working in Western countries (Fig. 3C; median age at PhD conclusion: Western Europe = 29 years; Southern and Eastern Europe = 30 years; *p* <.001). We did not have the information of where these researchers did their PhD, but we can infer that a significant percentage of the researchers obtained their PhD from an institution within the same European region where they were currently working. In fact, as we will describe later on this report, researchers working in Southern Europe were highly likely to have obtained their PhD from their current institution.

No differences were found across gender (Fig. 3B; median age at PhD conclusion = 30 years for both men and women). Yet, age at PhD conclusion varied significantly with research area (Fig. 3D; median age at PhD conclusion: SocSci&Hum&Econ = 31 years; Env&Geo = 30 years; LifeSci, Chem&Phys&Maths and IT&Eng = 29 years; *p* <.001). This finding might explain at least in part why postdoctoral researchers in social and environmental sciences were older than their counterparts.

##### Q5 and Q10 Nationality and Country of work

Table 1 shows the top 10 countries where the people surveyed work and the top 10 nationalities. Most of the participants answering the survey worked in **Portugal** (256), **UK** (178), **Sweden** (162), **Spain** (76) and **Germany** (57) and were born in Portugal (253), Spain (118), Italy (107), UK (55) and Germany (51). People working in 27 different European countries and 54 different nationalities answered the survey (Annex 2 and 3).

**Table 1.** Top 10 countries of work and countries of origin.

| In which country do you currently work? |  | What is your nationality? |  |
| --- | --- | --- | --- |
| Country | No. of people / % | Country | No. of people / % |
| 1- Portugal | 256 / 29% | 1 - Portugal | 253 / 28% |
| 2 - UK | 178 / 20% | 2 - Spain | 118 / 13% |
| 3 - Sweden | 162 / 18% | 3 - Italy | 107 / 12% |
| 4 - Spain | 76 / 8% | 4 - UK | 55 / 6% |
| 5 - Germany | 57 / 6% | 5 - Germany | 51 / 6% |
| 6 - Italy | 38 / 4% | 6 - France | 34 / 4% |
| 7 - France | 23 / 3% | 7 - India | 26 / 3% |
| 8 - Switzerland | 13 / 1% | 8 - Sweden | 24 / 3% |
| 9 - Denmark | 12 / 1% | 9 - Poland | 18 / 2% |
| 10 - Netherlands | 10 / 1% | 10 - Greece | 14 / 2% |

Although within our surveyed sample **the highest percentage of postdoctoral researchers were working in Western Europe (54%), the majority of survey respondents was from Southern Europe** (55%; Figure 4A). Twelve percent of postdoctoral researchers working in Europe were from outside Europe. We found a significant effect of gender (Fig. 4B; *p* <.001). Within the studied male postdoctoral researchers sample there was a higher percentage of researchers from non-European origin than within the female researchers. In turn, within the female researchers there was a higher percentage of researchers from Southern Europe.

**Figure 4.**
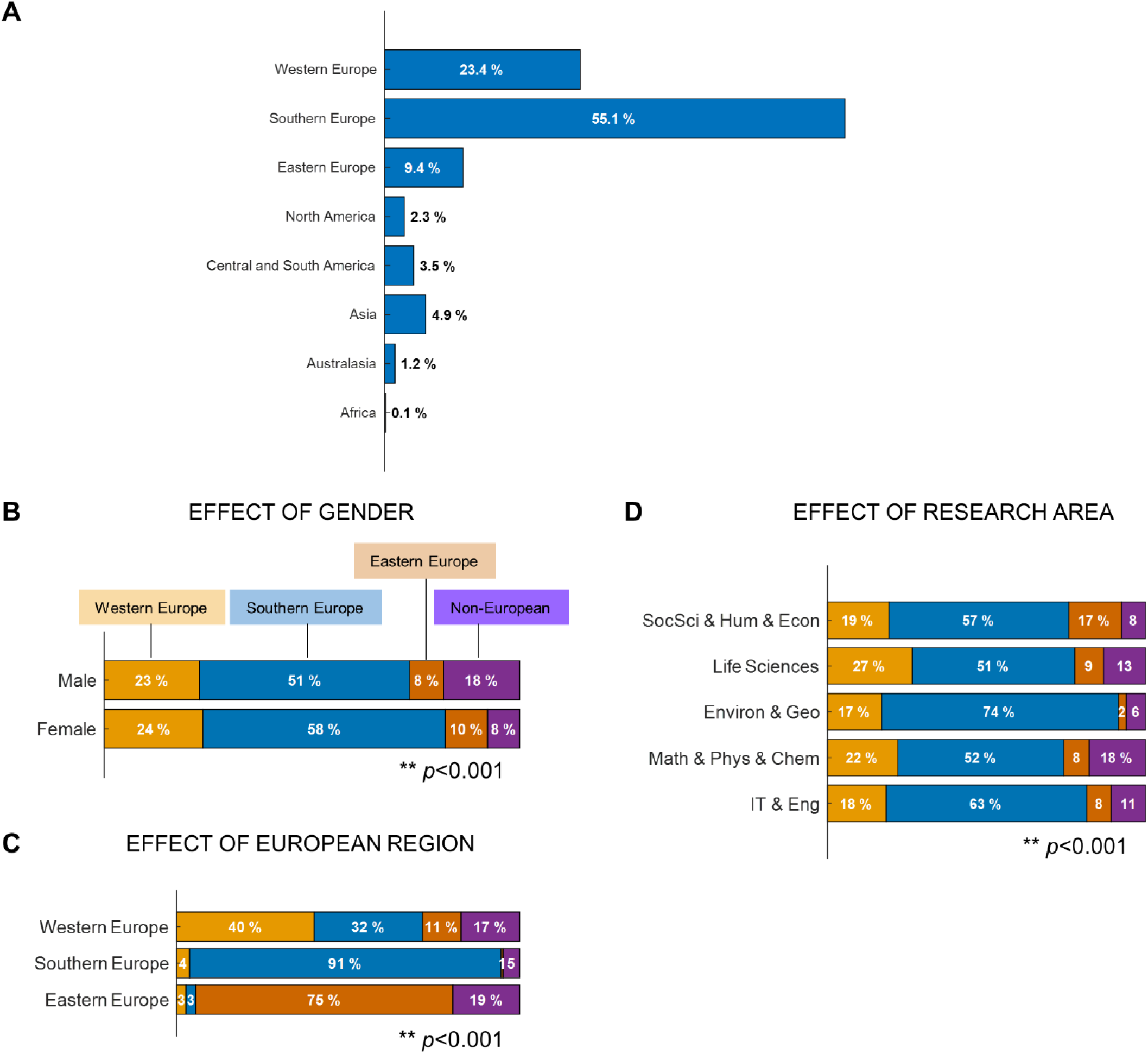
Nationalities of postdoctoral researchers working in Europe.

The majority of postdoctoral researchers working in Southern or Eastern Europe were from Southern (91 %) and Eastern (75 %) Europe, respectively (Figure 4C). In contrast, only 40% of postdoctoral researchers working in Western Europe were originally from this region. This observation highlights the **poor ability of Southern and Eastern European institutions to attract researchers from other European regions**.

The analysis of the mismatch between country of origin and country of work showed that **around half of the researchers surveyed (53%) did not currently work in their country of origin**. There was a significant effect of European region (*p* <.001; Fig. 5A). Western Europe was the European region with larger number of researchers working outside their country of origin. There were no significant differences between the proportion of male and female researchers working abroad (Fig. 5B; *p* =.090). This proportion varied, however, with research area with Chem&Phys&Maths presenting the highest proportion of researchers working outside their country of origin and SocSci&Hum&Econ presenting the smallest proportion (Fig. 5C; *p* <.001).

**Figure 5.**
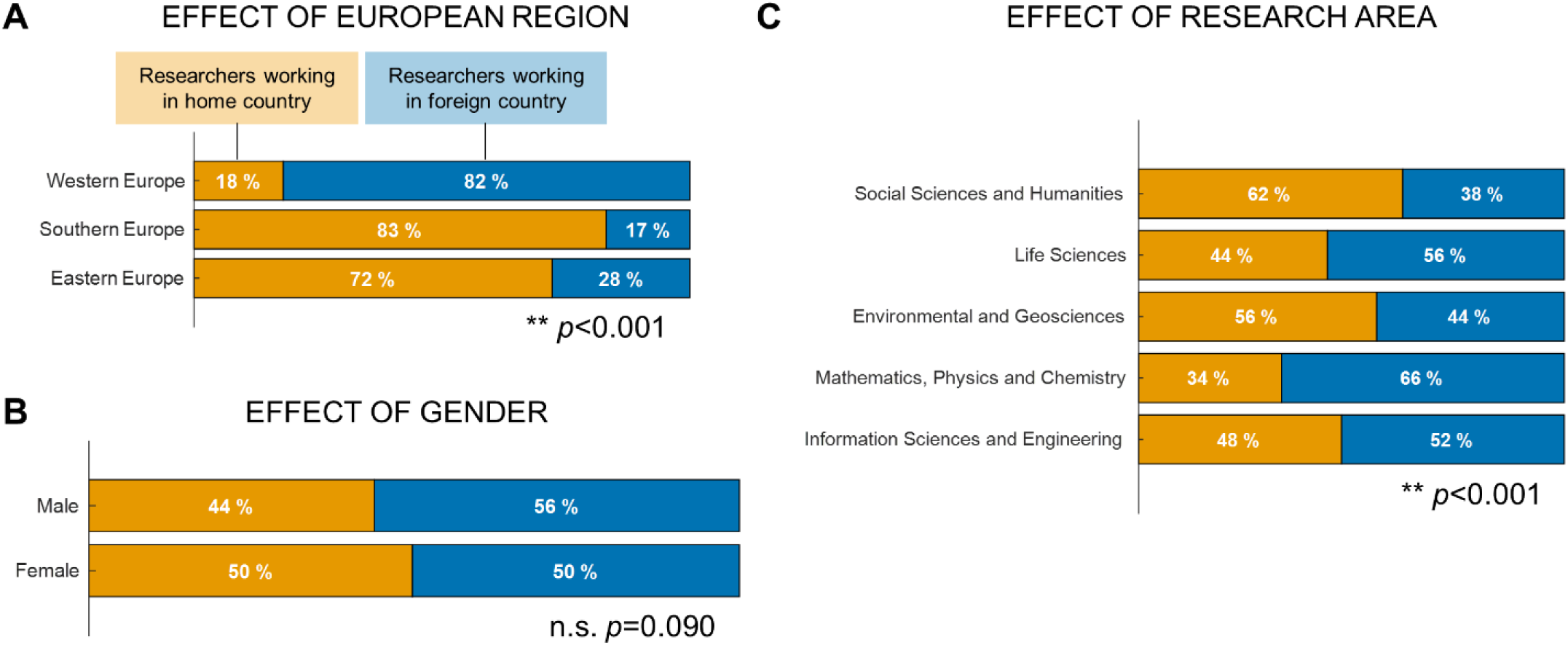
Percentage of postdoctoral researchers working in their home country or abroad.

##### Q6 Fluency on the language of the country of work

Considering only the researchers that were working in a foreign country, **more than half (54%) did not speak or speak with difficulty the language of the country of work** (Fig. 6A). A higher percentage of foreign postdoctoral researchers working in Eastern Europe did not speak the language of the country of work in comparison with the other European regions (Fig. 6C), yet given the small number of respondents from this region this difference was not statistically relevant. There was no significant effect of gender but a significant effect of research area was observed: researchers working in Life Sciences were less likely to speak the language and researchers working in Env&Geo were more likely to speak the language (Fig. 6D; *p* <.001).

**Figure 6.**
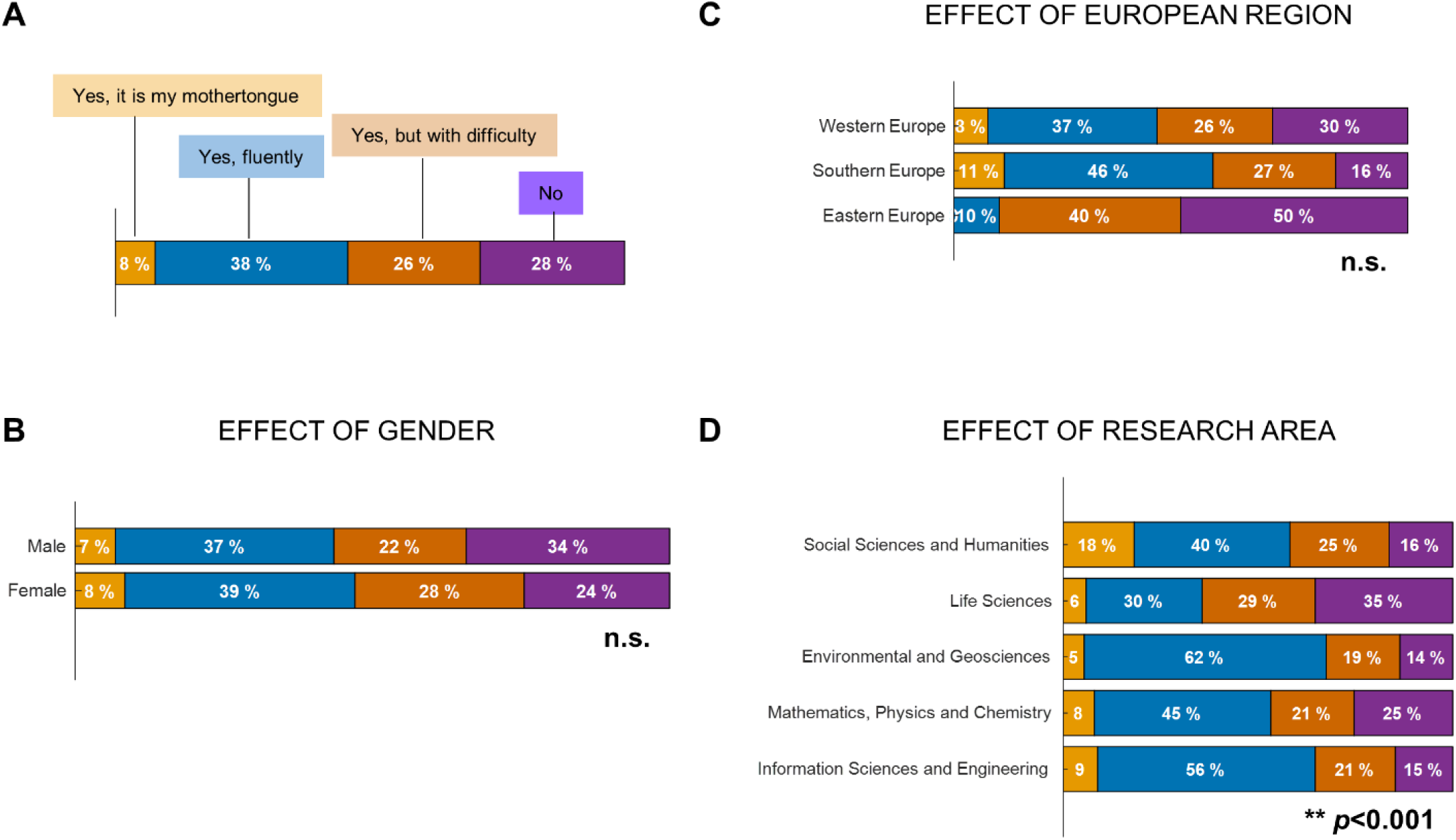
Whether postdoctoral researchers working in a foreign country speak the language of the country of work.

##### Q7-8 Marital status and cohabitation

Among the postdoctoral researchers that answered the survey, 63% were married or cohabiting, 11% were in a relationship but not cohabiting, 2% divorced or widowed and 23% were single. These rates seem to be in agreement with the EU average of 55.3 % and 28.1 % of people aged 20 or over that are married and single, respectively (<u>eurostat 2015</u>). There was a small but significant effect of gender (Fig. 7A; *p* =.03) with male researchers more likely than female researchers to be in a relationship. There was no significant effect of European region or research area (Figure 7 C and D). Also **these ratios did not significantly depend on researcher mobility, i.e. marital status did not depend on whether the researcher was working in their home country or not** (Figure 7B). Finally, around half (51%) of the researchers that were in a relationship but not cohabiting did not live in the same country as their partner, suggesting that moving country due to work requirements might be affecting the relationships of these researchers.

**Figure 7.**
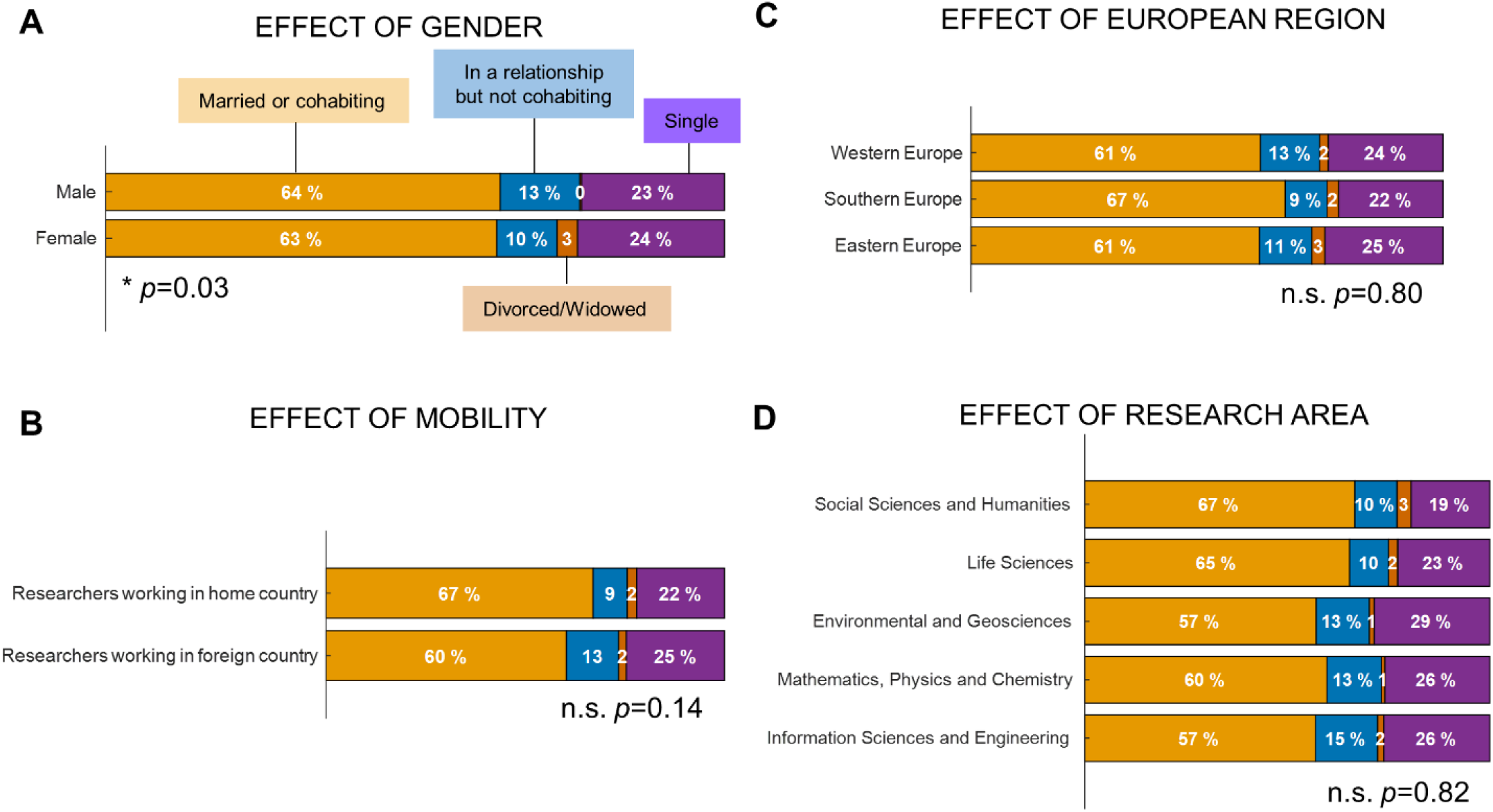
Marital status of postdoctoral researchers working in Europe.

##### Q9 Having children

**Thirty percent of the postdoctoral researchers** that answered the survey **had children**. This figure did not significantly depend on the researcher’s gender or research area (Fig. 8). It did, however, significantly depend whether the researchers were working in their home country or not. **Researchers working in their home country were twice as likely to have children compared with researchers working abroad (Fig. 8B)**. There are several reasons that might explain this observation. On the one hand, researchers that have children have more difficulty moving abroad. On the other hand, researchers working at home are more likely to have children given the extra support provided by extended family. It might also be in part explained by age differences. In fact, researchers working in their home country were significantly older than researchers working abroad (see Fig. 2C).

**Figure 8.**
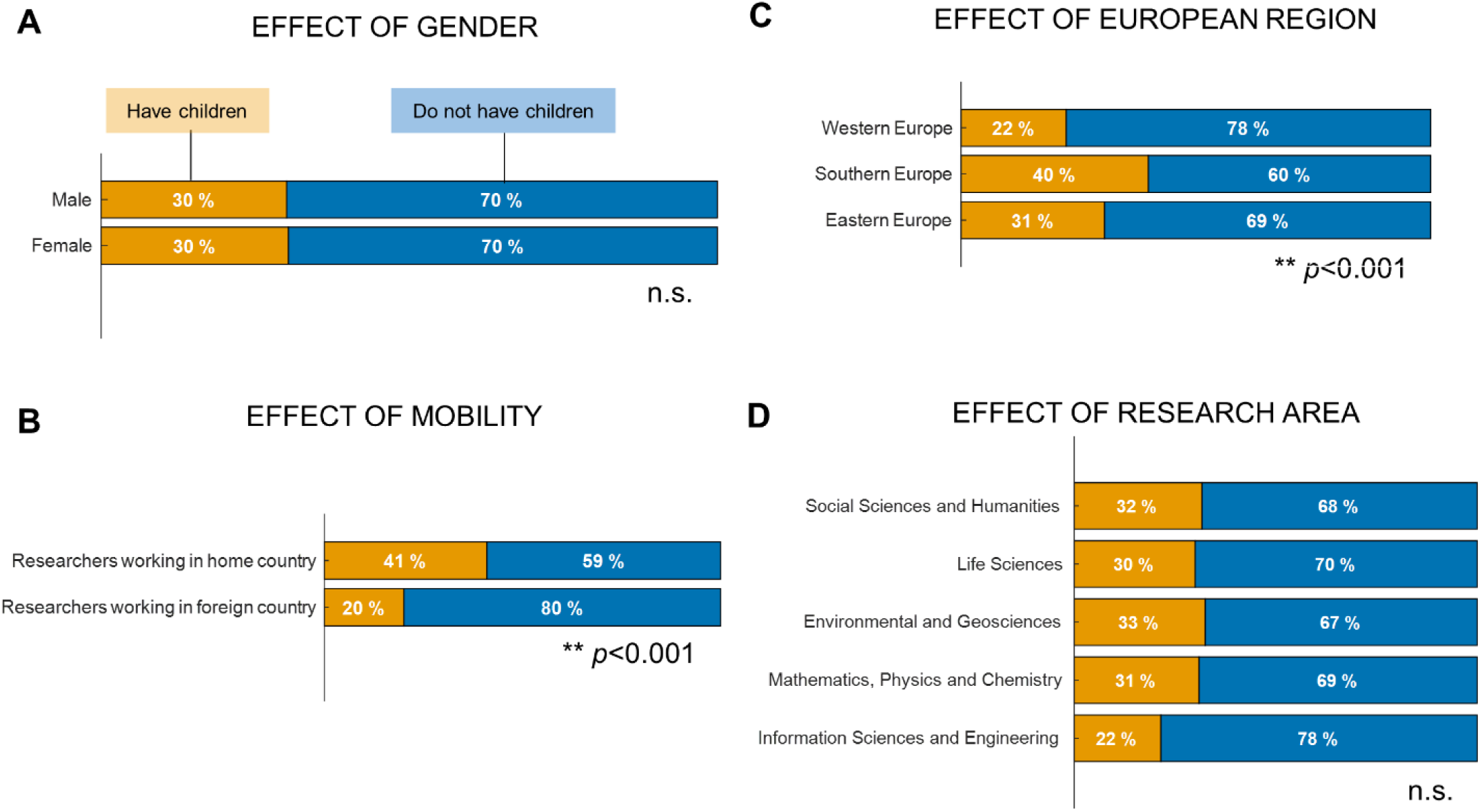
Percentage of the postdoctoral researchers that work in Europe that had children.

There was also a significant effect of European region with researchers working in Southern Europe being more likely to have children than researchers working in other parts of Europe (Fig. 8C). This might be explained by the fact that the great majority of researchers working in Southern Europe were working in their home country (>80%, see Fig. 5).

In order to further explore the factors influencing maternity/paternity choices within the researchers’ population, we studied the influence of mobility within each European region. As expected, within each European region, **researchers working in their home country were always more likely to have children than researchers working abroad** (data not shown). If we consider only researchers working in their home country, then the effect of European region is diminished and no longer significant (Fig. 9). These results suggest that mobility plays a significant role, more than region of work, on the maternity/paternity choices of postdoctoral researchers.

**Figure 9.**
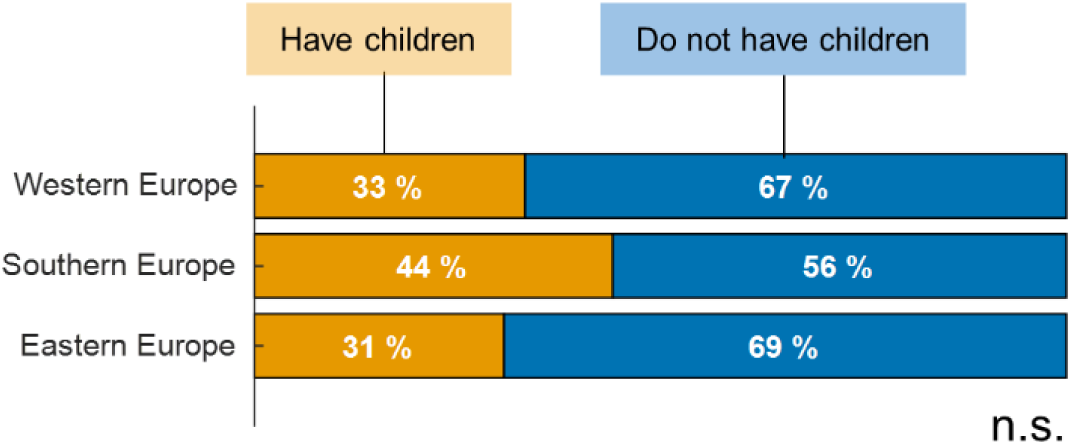
Percentage of postdoctoral researchers that had children within the sample of postdoctoral researchers that were working in their home country, as a function of European region.

#### 3.1.2 Working as a researcher

##### Q12 Primary research area

According to our survey data, the percentage of postdoctoral researchers working in each research area varied across European regions (Fig. 10; *p* <.001). LifeSci was the predominant research area in Western and Southern Europe, while in Eastern Europe the research area group of SocSci&Hum&Econ predominated. Southern Europe presented the largest percentage of postdoctoral researchers working in Env&Geo, while Eastern Europe presented the highest proportion of postdoctoral researchers working in Chem&Phys&Maths and SocSci&Hum&Econ. IT&Eng was the research area with the smallest number of researchers in all European regions.

**Figure 10.**
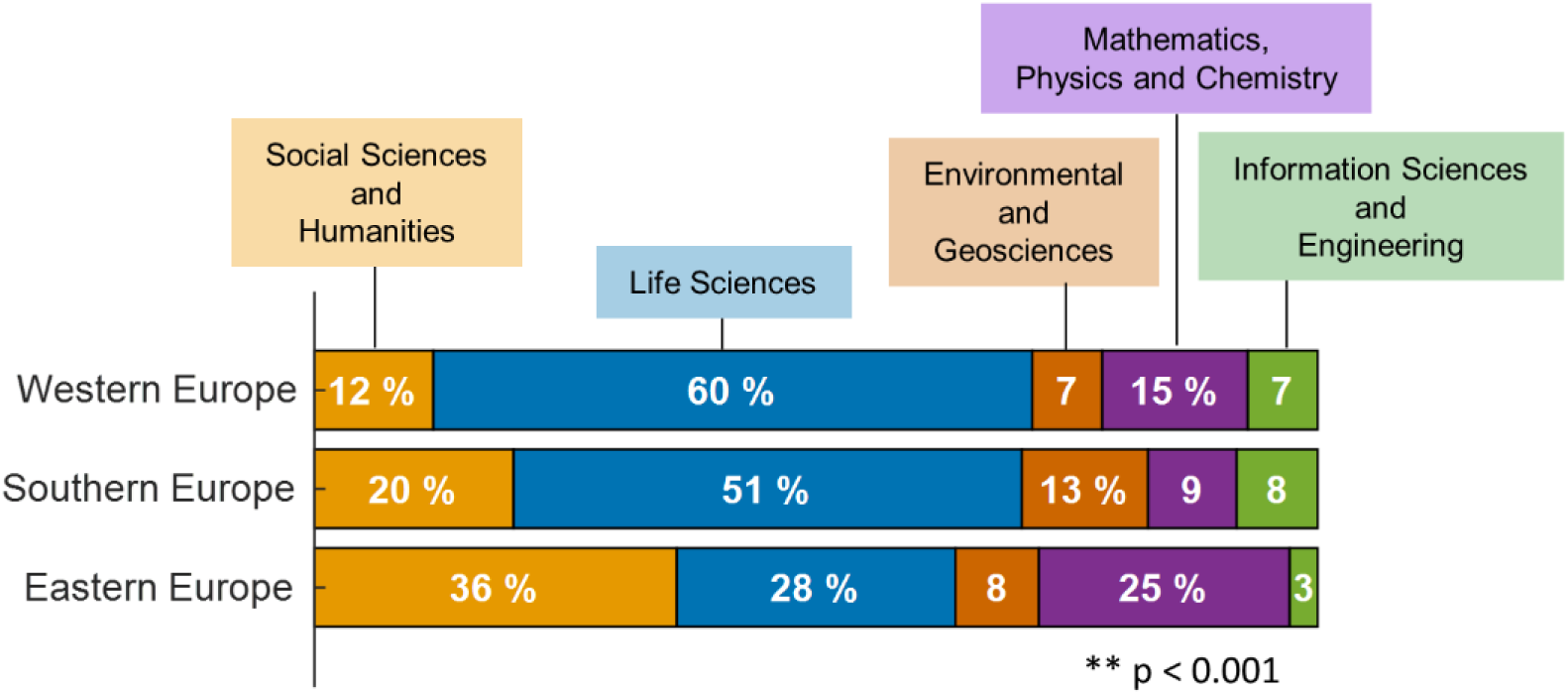
Percentage of postdoctoral researchers working in each of the research area groups within each European region.

##### Q14 Year of PhD conclusion

Researchers holding postdoctoral positions in Europe at the time of survey completion (at the end of 2017) had on average finished their PhD 4 years previously. The median (Interquartile Range - IQR) of the year of PhD conclusion was 2013 (IQR = 2011-2015).

##### Q15 Mobility after PhD conclusion

Thirty one percent of the postdoctoral researchers surveyed obtained their PhD at their current institution, 59% obtained their PhD at another European institution, while 9% obtained their PhD at a non-European institution. The majority of researchers that obtained their PhD outside Europe were from a non-European nationality (74%), meaning that **only a small minority (2%) of European researchers move outside Europe to do their PhD and then come back to Europe for postdoctoral positions**. These results presented a small but significant effect of gender (Fig. 11A; *p* =.05) with female researchers more likely to have obtained their PhD at their current institution and male researchers more likely to have obtained their PhD at a non-European institution. There was a highly significant effect of mobility (Fig. 11B; *p* <.001). Researchers working in their home country were significantly more likely to have obtained their PhD in their current institution. Also there is a significant effect of European region, with **researchers working in Western Europe less likely to have obtained their PhD at their current institution** (*p* <.001, Fig. 11C). There was no significant effect of research area (Fig. 11D).

**Figure 11.**
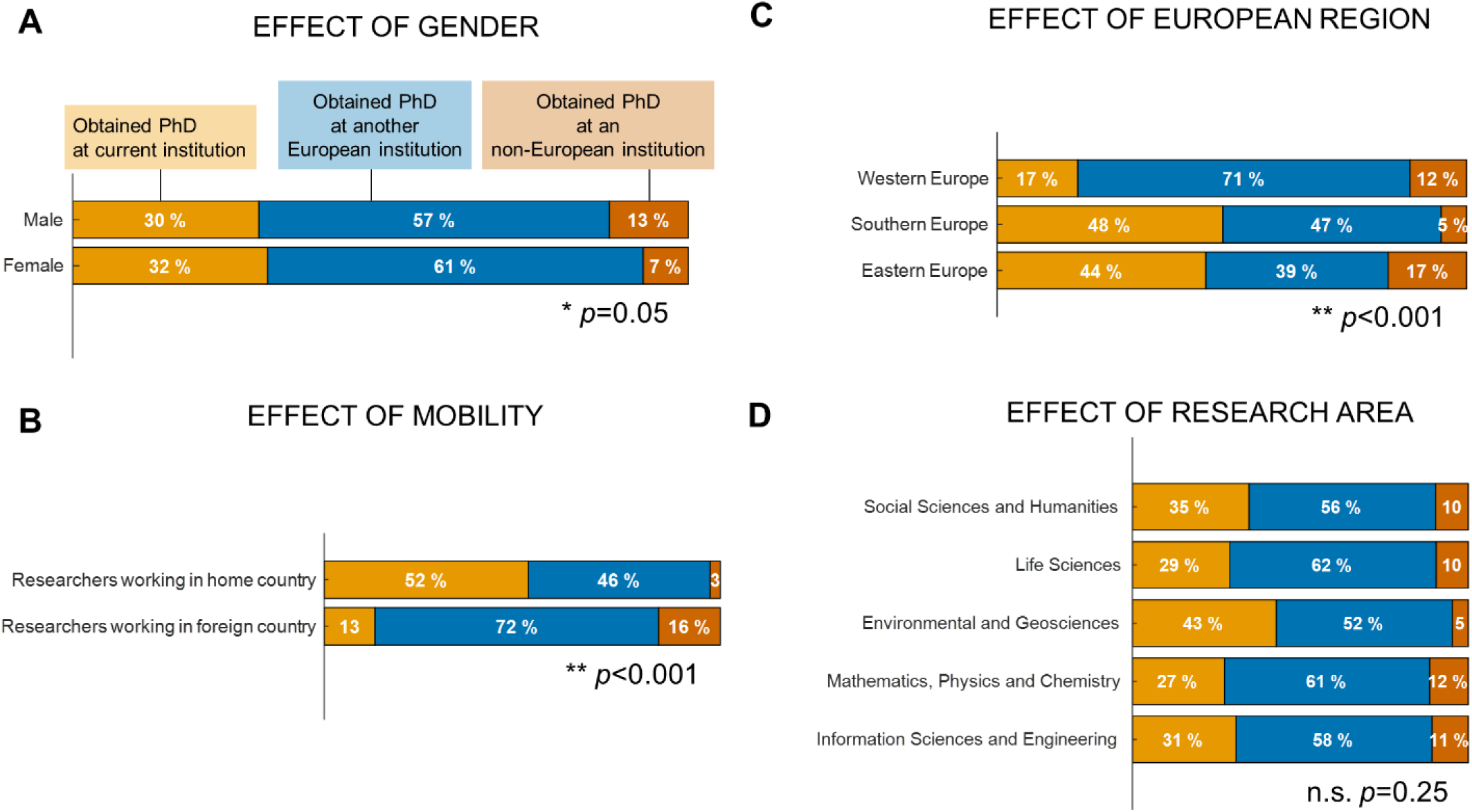
Postdoctoral researcher’s mobility after PhD conclusion.

##### Q16 Time as a postdoctoral researcher

The European postdoctoral researchers surveyed reported that they had been working as graduate researchers for an average of 40 months (3.3 years). Yet, this number varied significantly across European regions with Southern countries harbouring researchers that were working as postdoctoral researchers for longer (medians: Western = 36 months; Southern = 48 months; Eastern = 34 months; Fig. 12A). Moreover, postdoctoral researchers working in their home country reported longer postdoctoral periods (Fig. 12B). Finally, there was no effect of gender or research area (Fig. 12C and D; *p* >.05).

**Figure 12.**
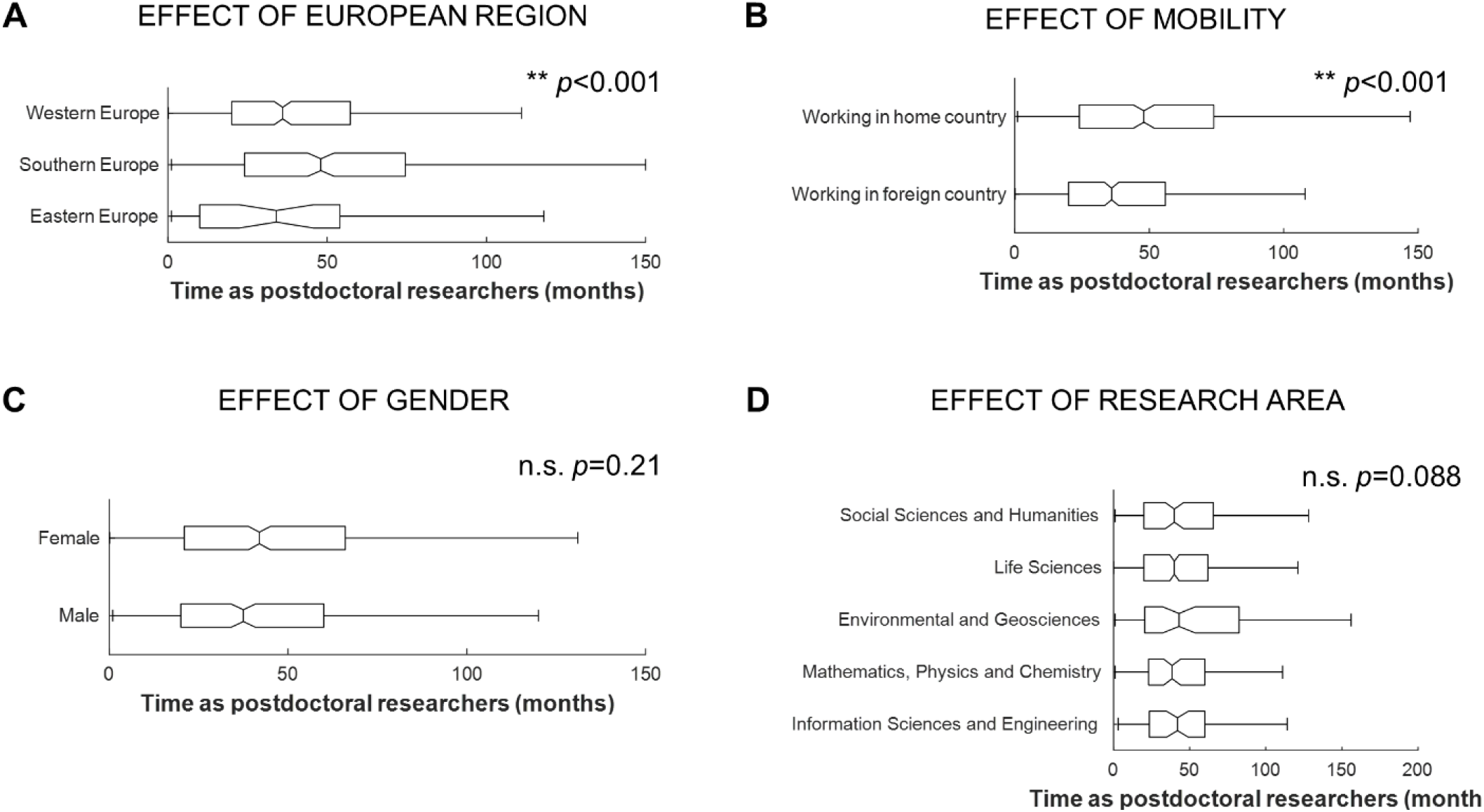
Amount of time, in months, that postdoctoral researchers have been working as graduate researchers.

##### Q17 Previous postdoctoral position

Overall, **43% of postdoctoral researchers were in their first postdoctoral position**, 15% had a previous postdoctoral contract in their current institution, 13% had a previous postdoctoral contract in the same country but different institution, while 29 % had a previous postdoctoral contract in a different country.

These percentages did not depend on the researchers’ gender, but depended on European region, research area and on whether the researcher was working at their home country or not (Fig. 13). Researchers working in their home country were more likely to have had a previous postdoctoral contract in their current institution (*p* <.001). In line with the fact that Southern Europe has the highest number of postdoctoral researchers working in their home country, this European region also showed the highest percentage of postdoctoral researchers that have had a previous postdoctoral contract at their current institution (*p* <.001). Chem&Phys&Maths is the research area group with highest number of postdoctoral researchers that have moved country between postdoctoral contracts (*p* <.001).

**Figure 13.**
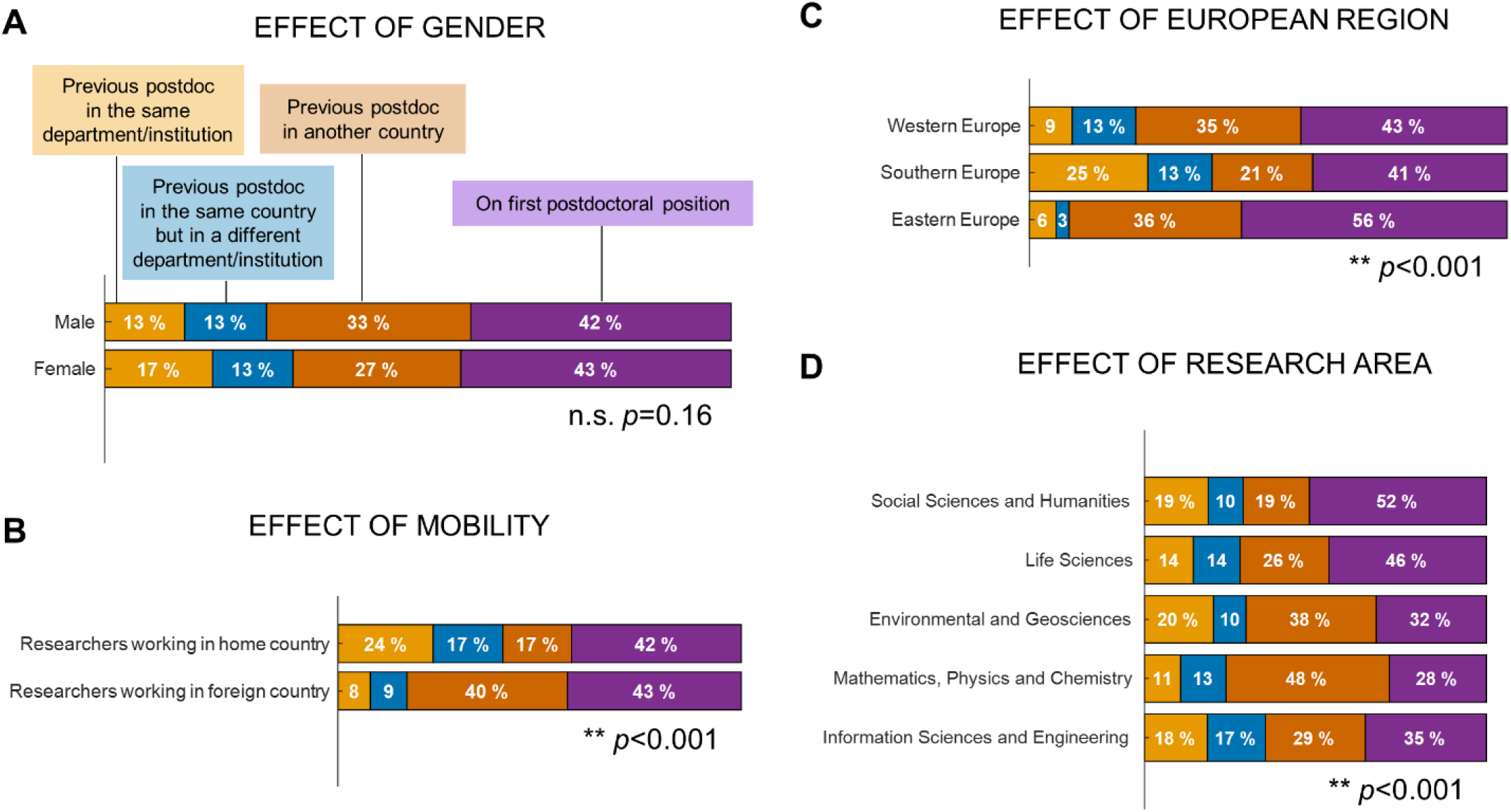
Percentage of postdoctoral researchers on first contract or on subsequent contracts, and whether or not they moved institutions between postdoctoral contracts.

##### Q18 Nature of current fellowship /contract

**Eighty nine percent of the postdoctoral researchers surveyed held a fixed term contract or fellowship**, 4% held a permanent or open-ended contract and 3% a casual, hourly paid arrangement. These percentages did not depend on gender or research area (Fig. 14). They did however depend on mobility and European region. Researchers working in their home country and researchers working in Eastern Europe reported more often to have an open-ended or permanent contract than their counterparts (Fig. 14B and C). However, these percentages were relatively small.

**Figure 14.**
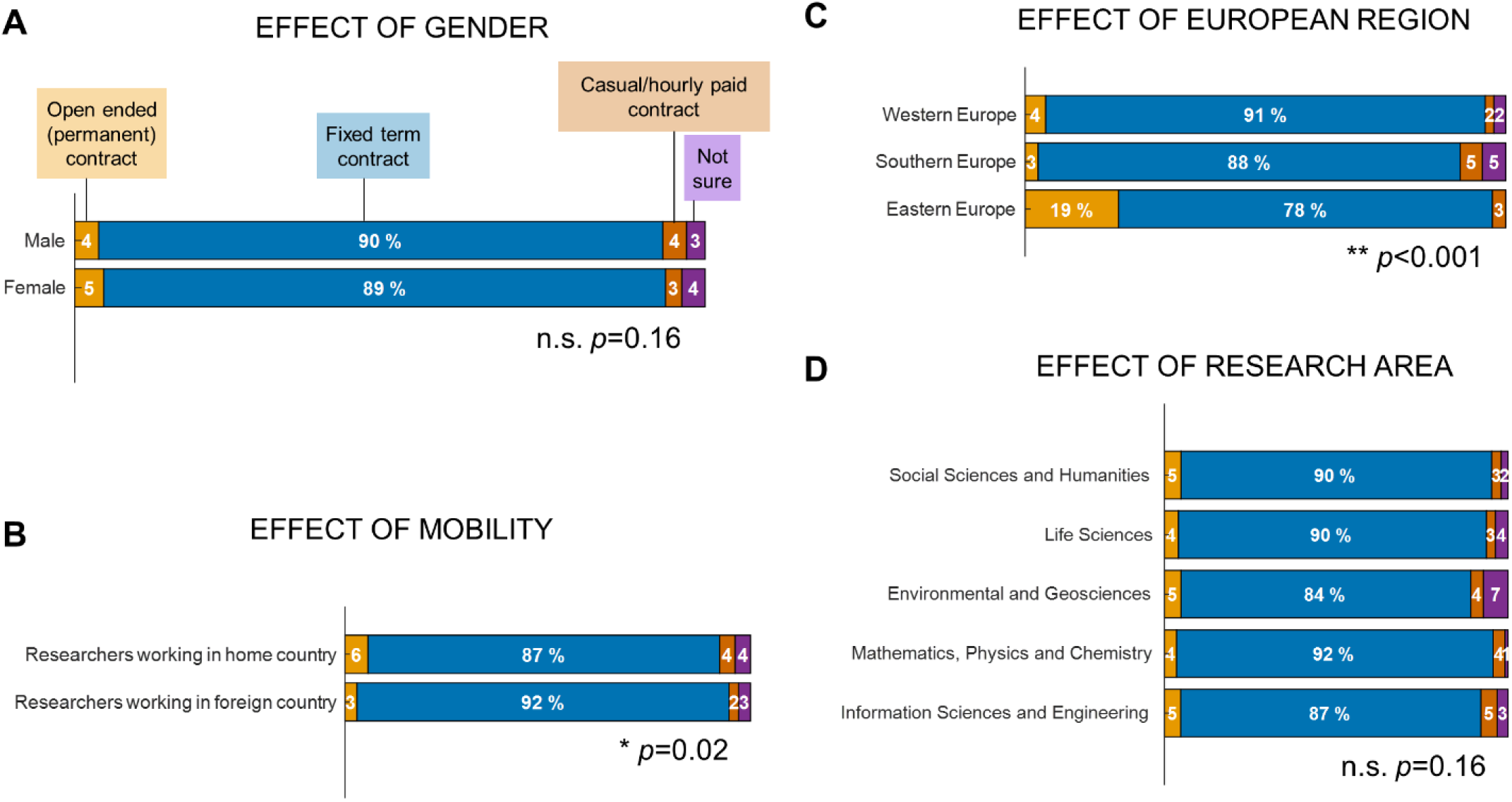
Nature of contracts of the postdoctoral researchers working in Europe.

##### Q19 Funding source

In terms of funding source, over half of the postdoctoral researchers surveyed (55%) were funded by a research grant awarded to a project, 16% were funded by their institution core funding and 27% were funded by their own research fellowship. The effect of gender was marginally significant (Fig. 15A; *p* =.09), with male researchers more likely to be funded by their institution core funding and female researchers more likely to have their own funding. There was a significant effect of European region (Fig. 15B; *p* <.001), with Southern Europe having a higher proportion of researchers with their own funding and Eastern Europe higher proportion of researchers being paid directly by their institution’s core funding. There was a significant effect of research area (Fig. 15D; *p* =.007), with researchers in IT&Eng more likely to be paid by their institution’s core funding and researchers in SocSci&Hum&Econ and Env&Geo more likely to have their own funding. Finally, open-ended contracts were more likely to be funded by the institutions’ core funding than the other types of contract (Fig. 15E).

**Figure 15.**
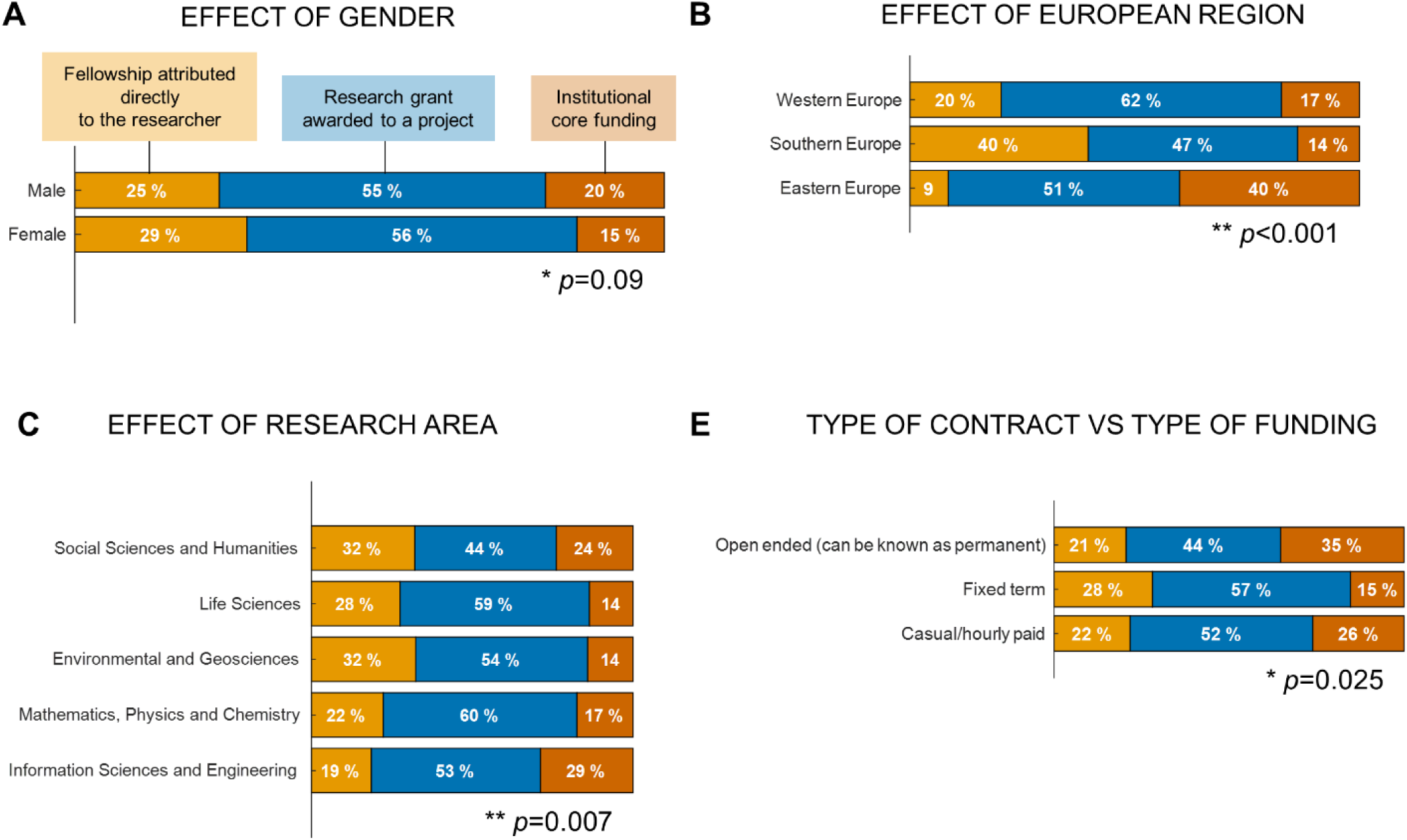
Sources of funding of postdoctoral positions of researchers working in Europe.

##### Q20 Full-time *versus* part-time contract/fellowship and exclusivity clauses

The **majority of postdoctoral researchers were on full time contracts** with only 4% of researchers on part-time contracts. Figure 16 shows the effects of gender, mobility, European region and research area. Women were more likely to be on part-time contracts than men, and SocSci&Hum&Econ was the research area group that had the highest percentage of researchers on part-time contracts. Interestingly, half of the part-time contracts were not by own choice but rather the only available option for these researchers.

**Figure 16.**
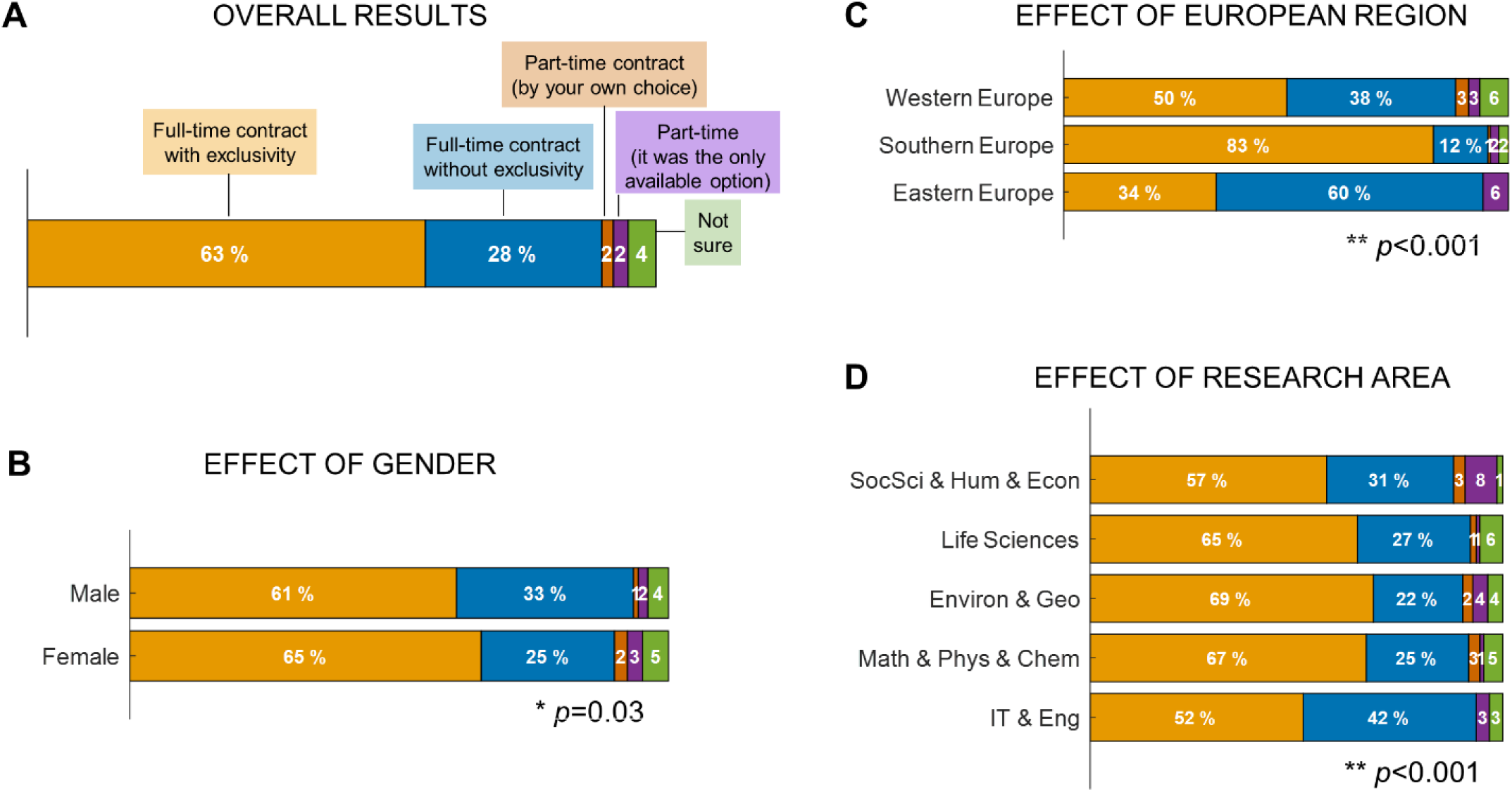
Percentage of postdoctoral researchers on full-time contracts with or without exclusivity and part-time contracts.

The majority of the contracts (63%) had exclusivity clauses meaning that the researchers were not allowed to have a second income (Fig. 16A). Researchers working in **Southern Europe were significantly more likely to have an exclusivity clause** in their contracts than researchers working in other regions (*p*<.001; Fig. 16C). Also, women were slightly more likely to have full time contracts with exclusivity clauses than men (*p*=.03; Fig. 16B).

##### Q21-22 Contracted hours *versus* hours at work

Thirty eight percent of postdoctoral researchers stated that their **work contracts did not include detailed information regarding working hours**. These were mostly from researchers working in Southern Europe (59%) but it also included researchers working in Western and Eastern Europe. When asked about the number of hours each researcher works per week, 79% of the researchers that had number of working hours specified in their contracts **stated that they worked more hours than the number of hours detailed in their work contracts** (on average 8 hours more per week than in their contracts; Fig. 17A). Thirty nine percent of researchers on full-time contracts stated that they work 50 hrs/week or more and 11% work 60 hrs/week or more.

**Figure 17.**
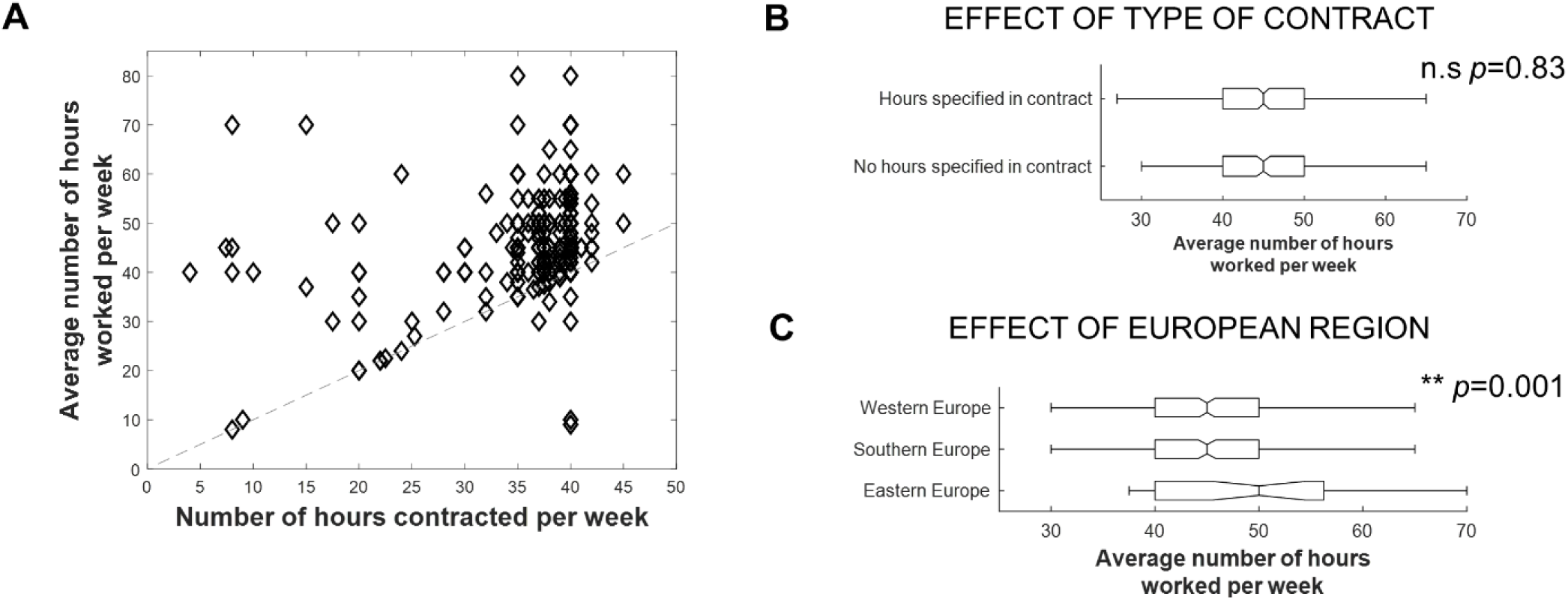
Number of hours reported at work and number of contracted hours. (A) Average number of hours researchers reported that they work per week plotted against number of hours specified in their contract. We included only the researchers that had contracts with number of hours specified. (B) Average number of hours researchers reported that they work per week for researchers that did and did not have working hours detailed in their contract. (C) Effect of European region on the number of hours researchers reported that they work per week.

Considering only researchers on full-time contracts, the number of hours that researchers reported working per week did not depend on whether the contracts included the details of working hours or not (Fig. 17B; *p* =.83). Moreover, it did not depend on gender or research area (data not shown). Researchers working in Eastern Europe stated that they worked on average more hours than researchers working in Western or Southern Europe (Fig. 17C; *p* =.001; median number of hours per week: Western Europe = 45; Southern Europe = 45; Eastern Europe = 50).

Notably, of the researchers that were on part-time contracts, 57% stated that they work more than 35 hours a week, i.e., as if they were on full time contracts.

##### Q23-24 Average annual income

Annual gross salaries of postdoctoral researchers were distributed on a broad range from around 5 000 euros up to over 70 000 euros, with a median of 32 000 euros (Fig. 18). This big range in salaries was mainly an effect of differences across European regions with researchers working in Eastern Europe earning the lowest salaries and researchers working in Western Europe earning the highest salaries (Fig. 18B; median annual gross salaries: Western Europe = 40 479 euros; Southern Europe = 18 000 euros; Eastern Europe = 15 600 euros; *p* <.001). Our data suggested that men earned more than women did (Fig. 18C; median annual gross salaries: men = 33 550 euros; women = 30 472 euros), however, this difference was no longer significant when adjusting for other known features affecting salaries (see Annex 6 for further exploration of this gender difference). We also observed a significant effect of research area, with SocSci&Hum&Econ being the research area group with the lowest average salary and IT&Eng earning the highest average salary (Fig. 18D; median annual gross salaries: SocSci&Hum&EcoN = 22 200 euros; LifeSci = 33 000 euros; Env&Geo = 31 200 euros; Chem&Phys&Maths = 33 975 euros; IT&Eng = 39 600 euros, *p* <.001).

**Figure 18.**
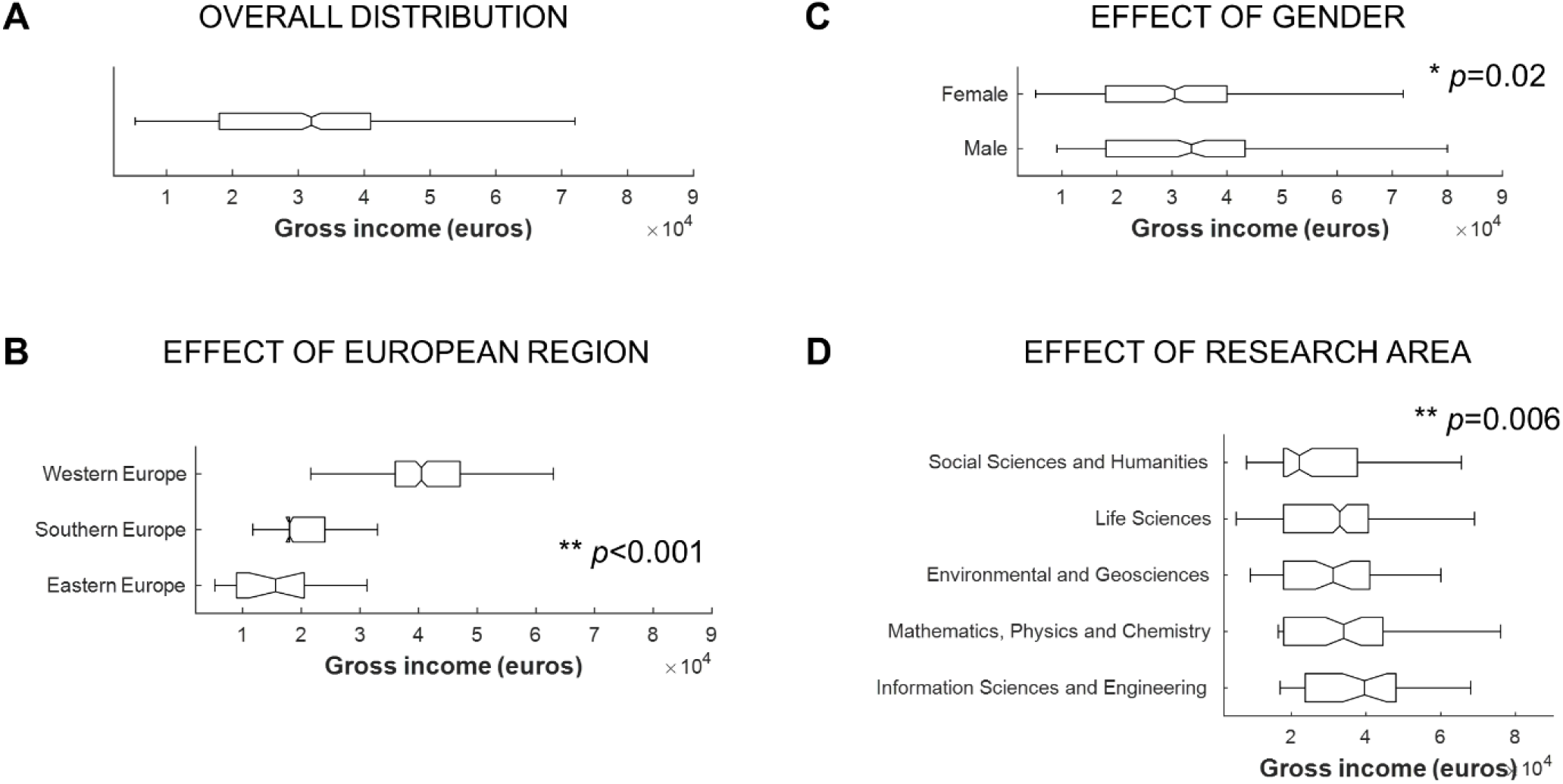
Gross salaries of postdoctoral researchers on full time contracts and working in Europe.

Similar results were observed in the analysis of yearly net salaries, except for the gender effect that was no longer significant (Fig. 19).

**Figure 19.**
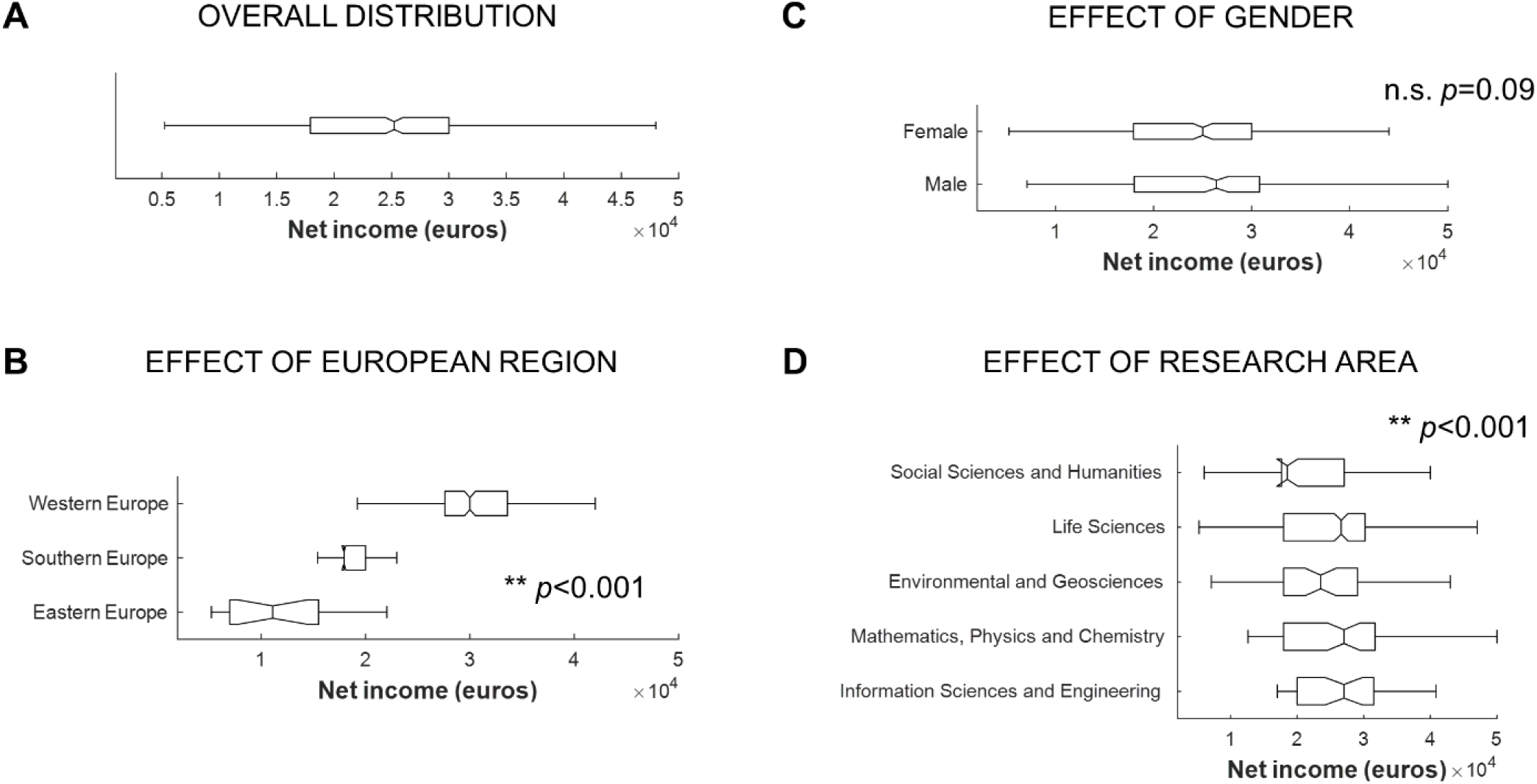
Net salaries of postdoctoral researchers on full time contracts and working in Europe.

##### Q25 Contract/fellowship benefits

When asked if their work contracts included provision for parental leave, access to healthcare, unemployment benefit and/or sick leave, unemployment benefit was the provision that was reported as being less frequently included (Fig. 20A). Moreover, **a significant percentage of researchers were not aware of their rights** and reported they were not sure, **highlighting the lack of knowledge researchers have regarding their rights to social benefits**. The answers depended on European region with researchers working in Southern Europe more likely to report that their contracts did not provide for the social benefits enumerated, while researchers working in Eastern Europe more likely to report that these benefits were provided for (Fig. 20B-E; *p* <.001). It is important to note however that these answers represent the perception of the researchers and not necessarily whether these provisions were included or not in their work contracts.

**Figure 20.**
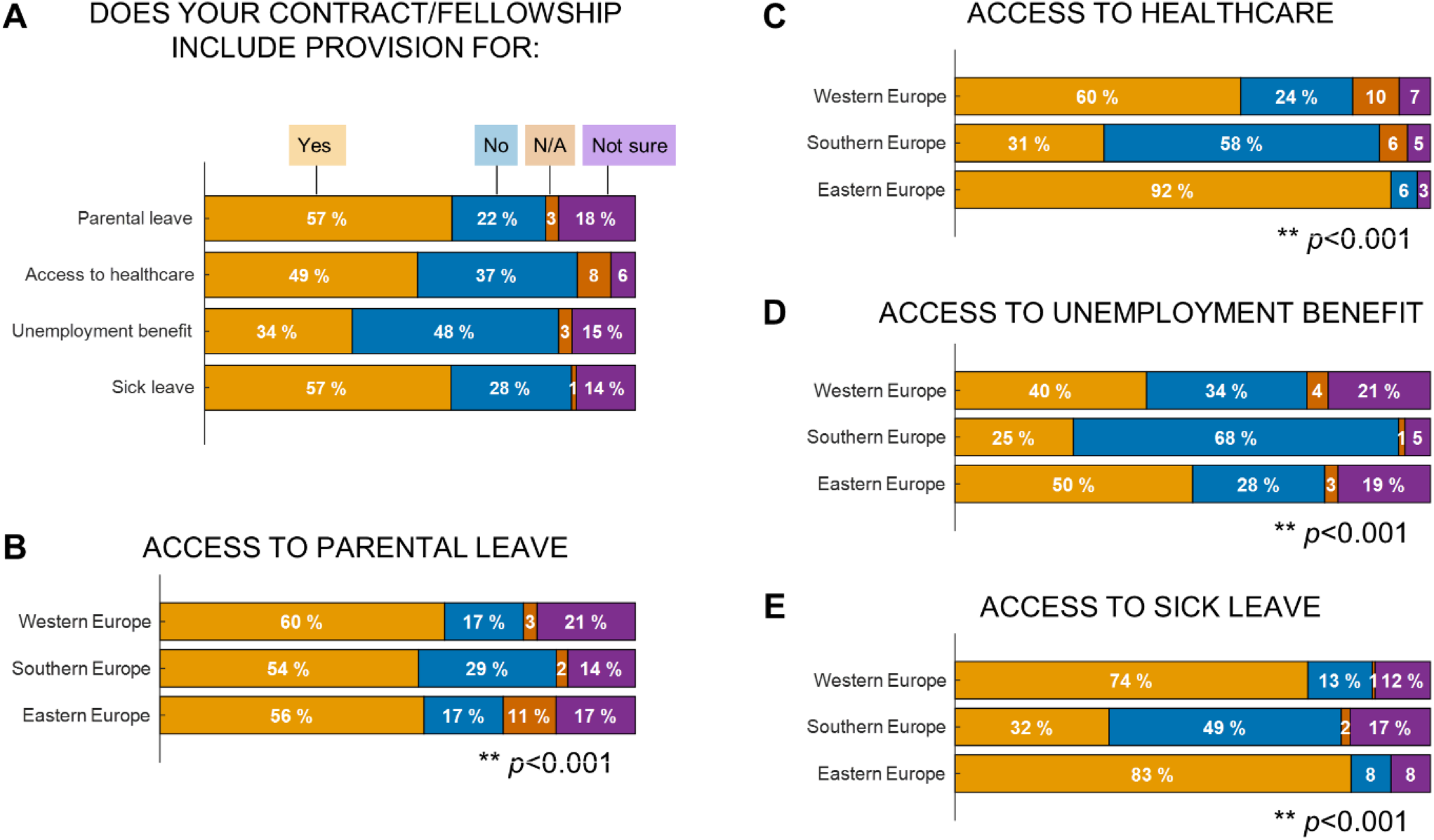
Postdoctoral researchers’ perception on the provision of employer benefits and how European region affects this perception.

#### 3.1.3 Productivity and teaching

Next, we will describe the scientific activities (productivity, teaching and supervision, and funding applications) of European postdoctoral researchers.

##### Q33-36 Publication record

Respondents reported having published a **median of 10 papers** (IQR = 5-16) in peer-reviewed journals. Less than half of all postdoctoral researchers working in Europe had published books (15.4%), book chapters (46.1%) or preprints (17.5%). There were no gender differences regarding number of papers, books, book chapters or preprints (all *p* >.05).

There were significant differences regarding number of papers in the different European regions and research areas (Fig. 21B and 21C). **Researchers working in Southern Europe had a higher number of peer-reviewed scientific papers** than researchers working in other European regions (medians = Western 8; Southern 12; Eastern 8.5; *p* <.001). Moreover, there was an effect of research area with researchers in Env&Geo presenting the highest number of peer-reviewed papers and researchers in SocSci&Hum&Econ the lowest number of publications (Fig. 21C; p<.001).

**Figure 21.**
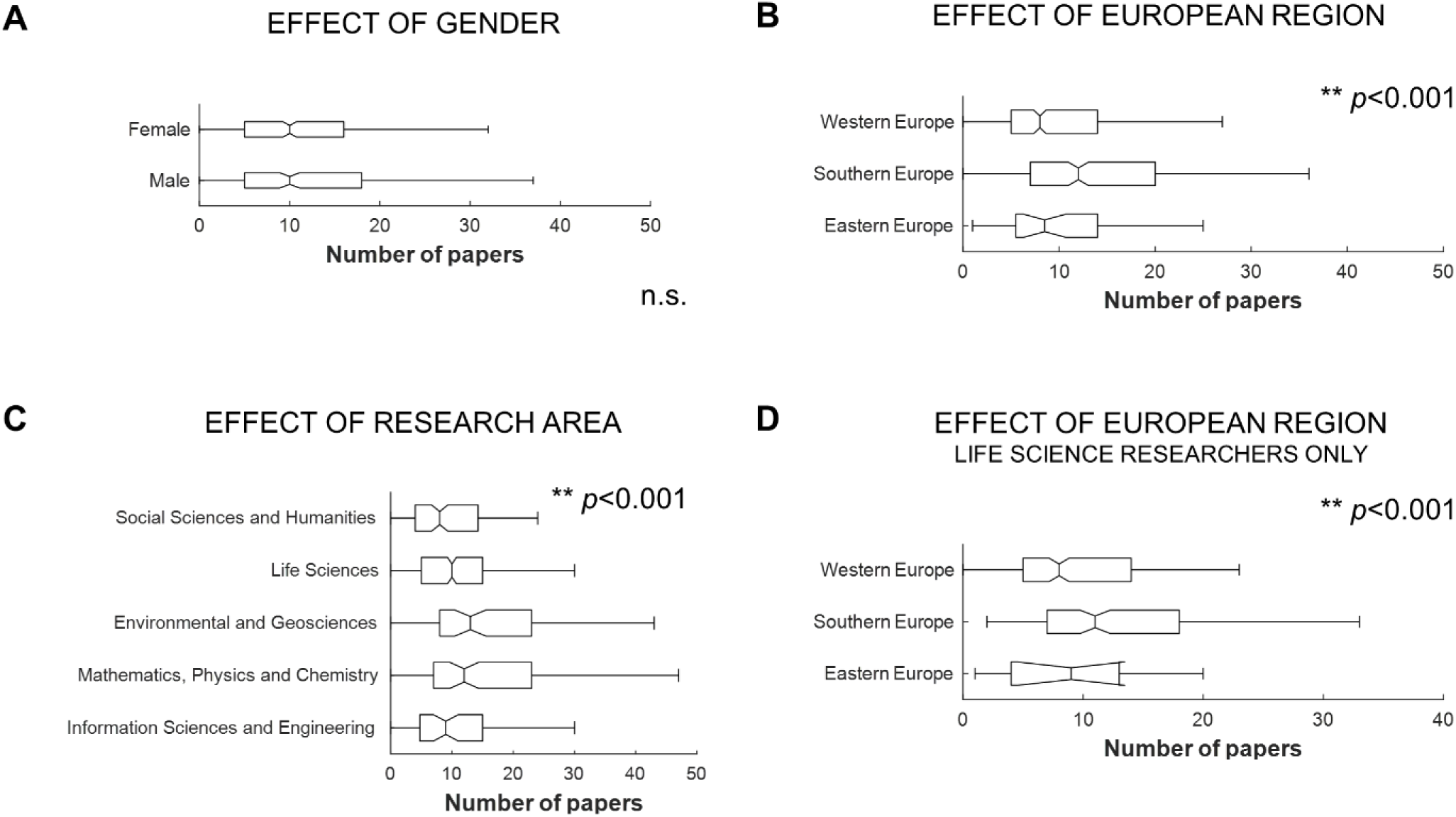
Number of peer-reviewed scientific papers that postdoctoral researchers working in Europe reported having published to date.

In order to determine if the effect across European regions was driven by differences in research areas we compared across European regions researchers working in Life Sciences only (the research area with biggest representation in our sample). A very significant effect of European region was still observed with researchers working in Southern Europe reporting having a higher number of publications (Fig. 21D; *p* <.001).

##### Q37-38 H-index

The h-index of European postdoctoral researchers had a median of 6 and an IQR of 4-9 (Fig. 22A). The h-index of these researchers varied significantly with European region being highest in Southern Europe (Fig. 22B; *p*<.001). We found no effect of gender (Fig. 22C; *p*=.88) and a very significant effect of research area (Fig. 22D; *p*<.001).The h-index increases with years of active publication <u>(Gadd 2018)</u>. Therefore, the difference across European regions could be a result of differences in time as a graduate researcher (see Fig. 12). In order to investigate this possibility, we fit a linear model including as independent variables researchers’ age, time as a researcher since PhD conclusion, method for h-index calculation and European region. The model was highly significant with most of the variance being explained by the relationship between time as a researcher since PhD conclusion and the h-index. However, even after controlling for these variables, the effect of European region was still significant, **with researchers working in Southern Europe having higher h-index than researchers working in the other European regions** (*p* =.016).

**Figure 22.**
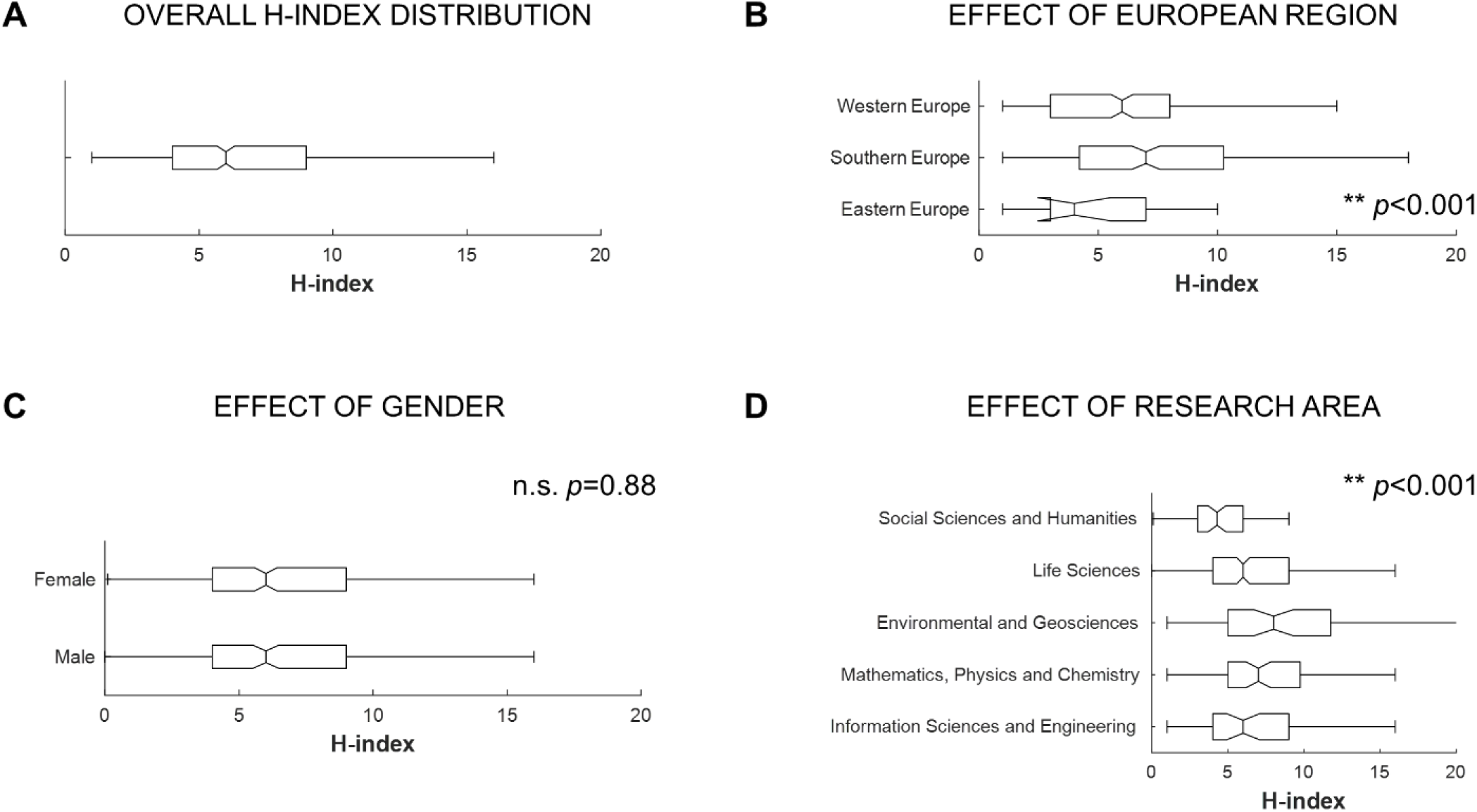
H-index distributions of the postdoctoral researchers working in Europe.

##### Q39 Acknowledgment of research contributions of postdoctoral researchers

The majority (76%) of the respondents reported that they were acknowledged as first authors in the publications when they did most of the work. Three percent of postdoctoral researchers reported that they were not acknowledged as first authors when they should have been, and 21% of respondents stated that they were acknowledged deservingly as first authors only in some publications. This result did not depend on gender *(p* =.32) or European region (*p* =.10), but there was a significant effect of primary research area with researchers in the Math&Phys&Chem and Social&Human&Econ fields being less likely to always being acknowledged as first authors when they should have been than researchers in other fields (Fig. 23; *p* <.001).

**Figure 23.**
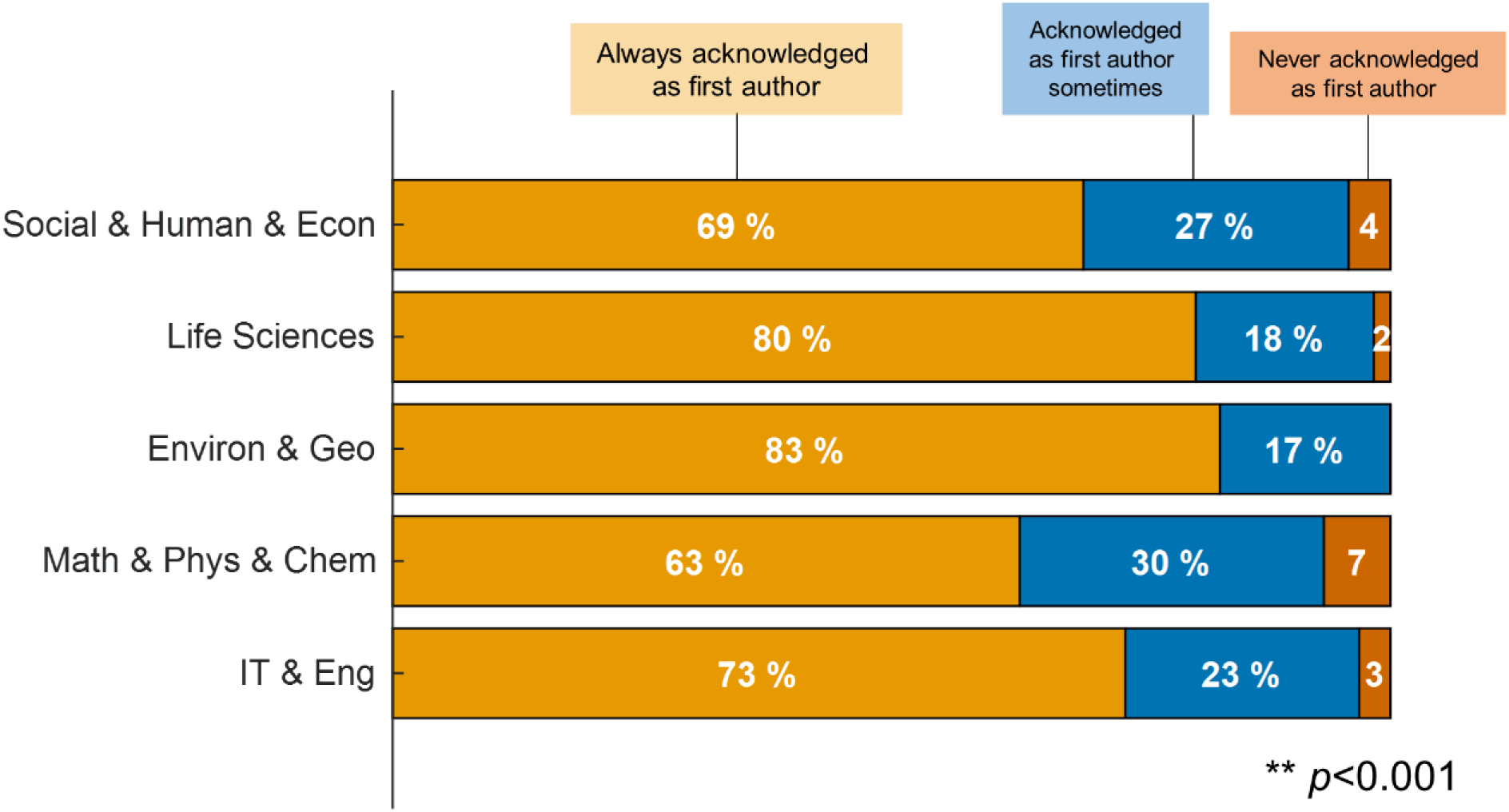
Perception of postdoctoral researchers regarding how they are acknowledged in the publications for which they have done the most significant contributions.

##### Q42 Disagreements on authorship order

About half (54%) of European researchers reported that they **had publications where they had not agreed with the authorship order** (Fig. 24). This disagreement was dependent on gender (*p* <.001; Fig. 24A) with more women than men reporting that there were situations where they disagreed with the authorship order. It was also dependent on European region (*p* = .015; Fig. 24B) with more researchers working in Southern Europe reporting situations where they disagreed. Finally, the differences across research areas were only marginally significant (*p* = .060; Fig. 24C).

**Figure 24.**
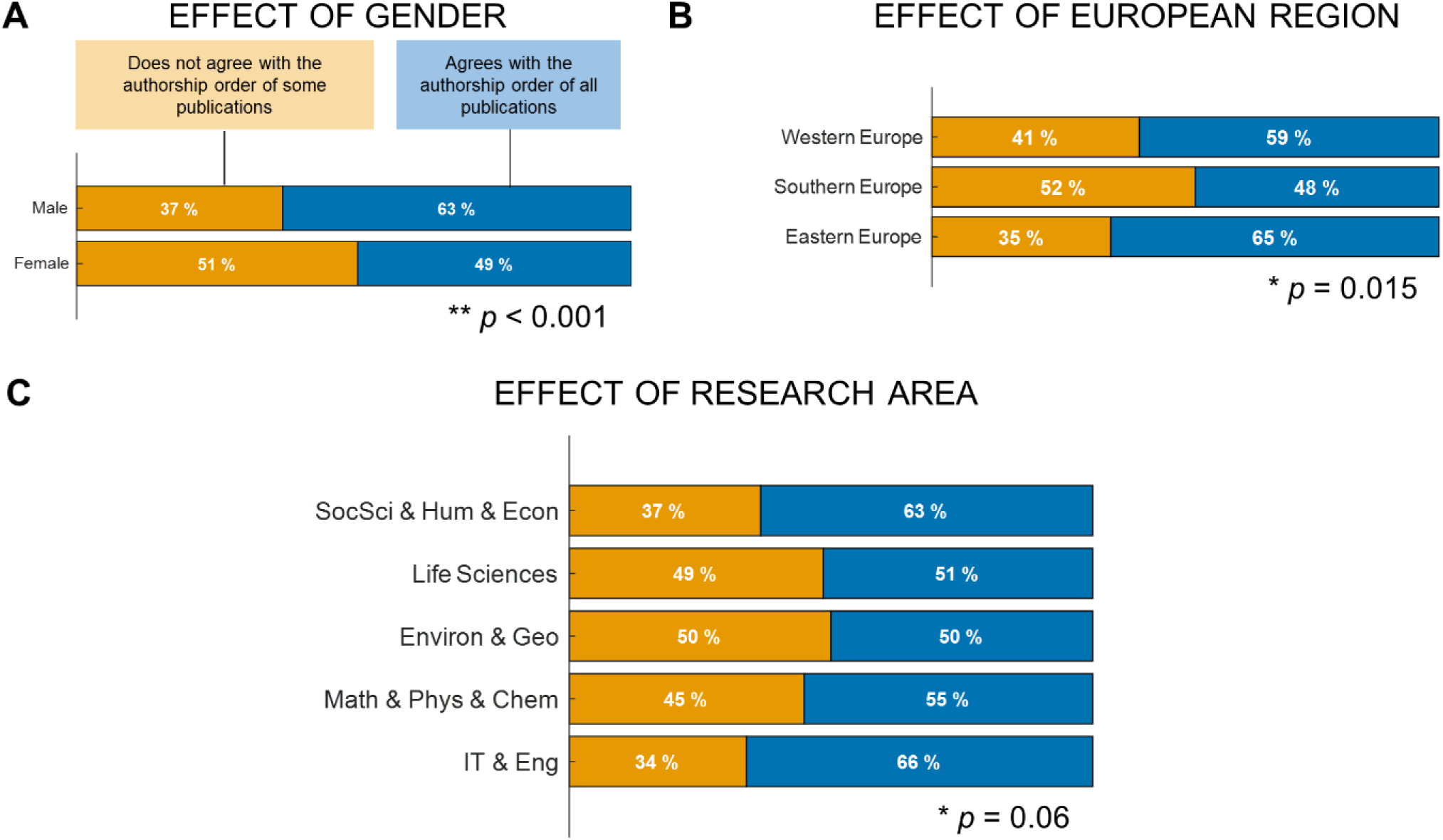
Percentage of postdoctoral researchers that agrees always with publications’ authorship order.

##### Q40-41 Rules on authorship and misconduct

**Less than half** of the surveyed postdoctoral researchers reported that their research group had **clear rules about authorship** or that their group had clear rules on **scientific misconduct** (Fig. 25A and B). There was an effect of European region on the clarity of rules of authorship (*p* <.038) and on rules of misconduct (*p* <.004), with researchers from Western Europe reporting more often that they did not know in both questions. Researchers from Eastern Europe reported less often that their group had clear rules on authorship than researchers from Southern and Western Europe and, in contrast, reported more often that their group had clear rules of misconduct. The answers to this question did not present an effect of gender or research area (*p* >.05; data not shown).

**Figure 25.**
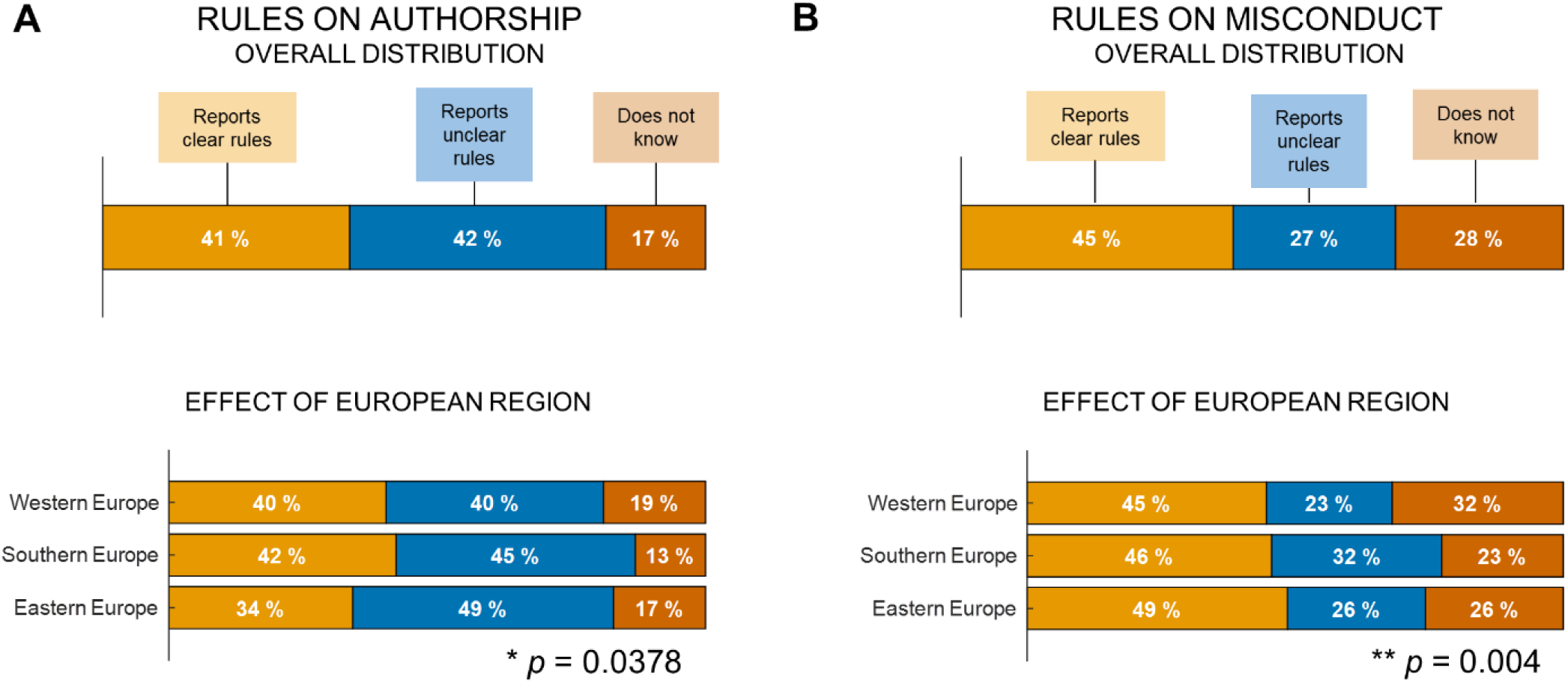
Perception of postdoctoral researchers working in Europe regarding the clarity of rules on authorship (A) and scientific misconduct (B) stipulated by their research group.

##### Q26 Teaching contribution

**Forty seven percent of postdoctoral researchers were involved in graduate or undergraduate teaching** (Fig. 26A). This number did not depend on the researchers’ gender (Fig. 26B). It did, however, depend on researchers’ mobility, with researchers working in their home country more likely to be involved in teaching (Fig. 26C).

**Figure 26.**
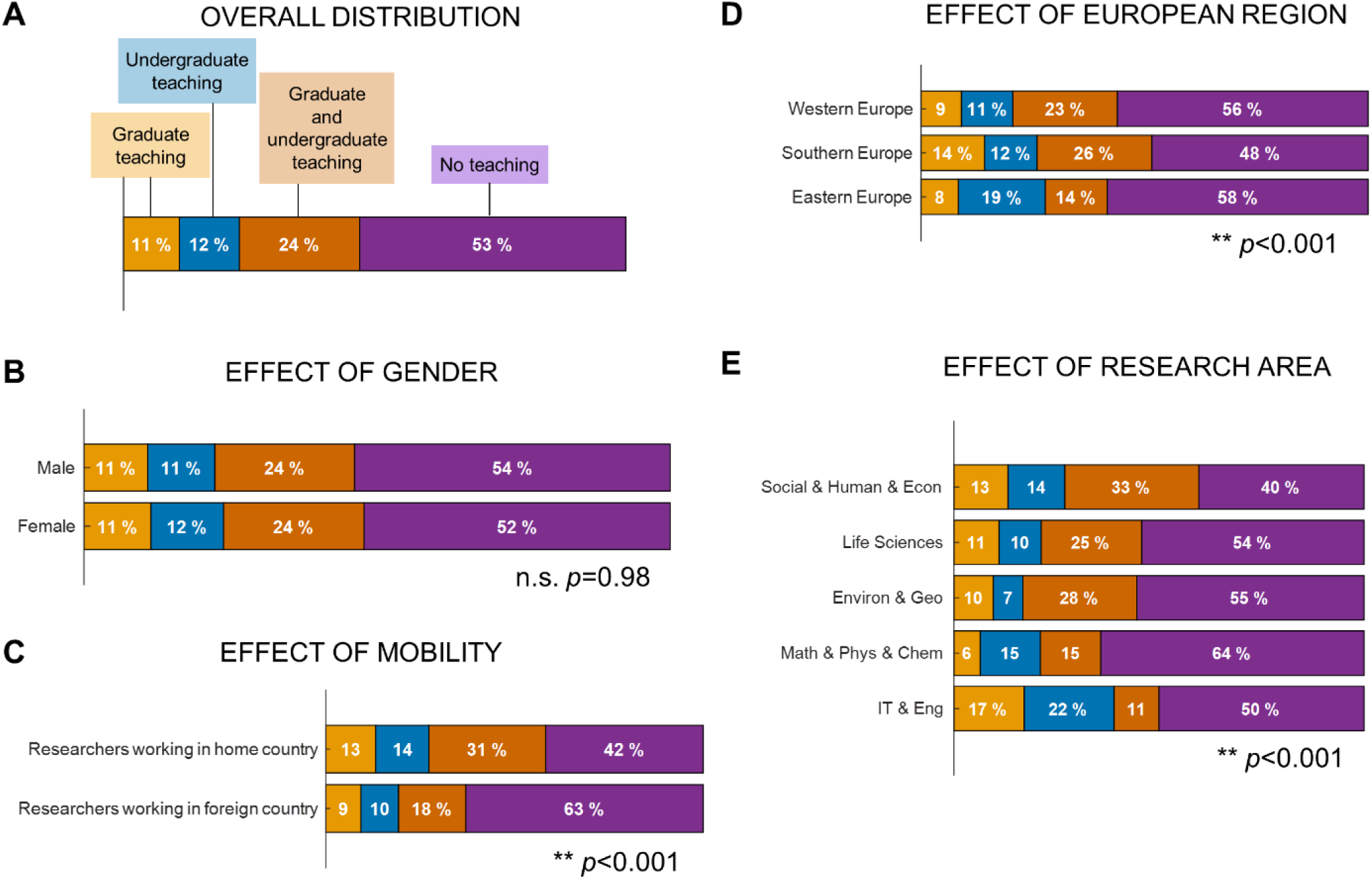
Teaching activities by postdoctoral researchers working in Europe.

It also varied with European region (researchers working in Southern Europe were more likely to be involved in teaching than researchers working in other parts of Europe, *p* <.001; Fig. 26D) and research area (researchers working in SocSci&Hum&Econ were more likely to teach and researchers working in Chem&Phys&Maths were less likely to teach; Fig. 26E; *p* <.001).

##### Q27-28 Teaching as a duty or an opportunity

Of those who teach, 15% say they are obliged to teach and of those who do not teach, 56% would like to teach but are not given the opportunity.

##### Q29-30 Supervising master/doctoral students

On the supervision of master students, 20% said they supervise but their supervision is not formally acknowledged and 6% were not allowed to do any type of supervision (Figure 27A1). Only a small fraction of postdoctoral researchers (1%) reported that they did not want to supervise master students. For doctoral student supervision, 25% said they supervise but their supervision was not formally acknowledged and 13% were not allowed to do any type of supervision (Figure 27B1). There was a significant effect of European region with researchers working in Southern Europe reporting more often that they were allowed to supervise or co-supervise both master and PhD students (Figure 27A2 and B2). Researchers working in Eastern Europe were more likely to report that their supervision was not formally recognized. There was also a significant effect of research area (Figure 27A3 and B3), with postdoctoral researchers within the research area group Math&Phys&Chem reporting more often that their supervision was not recognized and SocSci&Hum&Econ reporting more often that they were not allowed to supervise master or PhD students.

**Figure 27.**
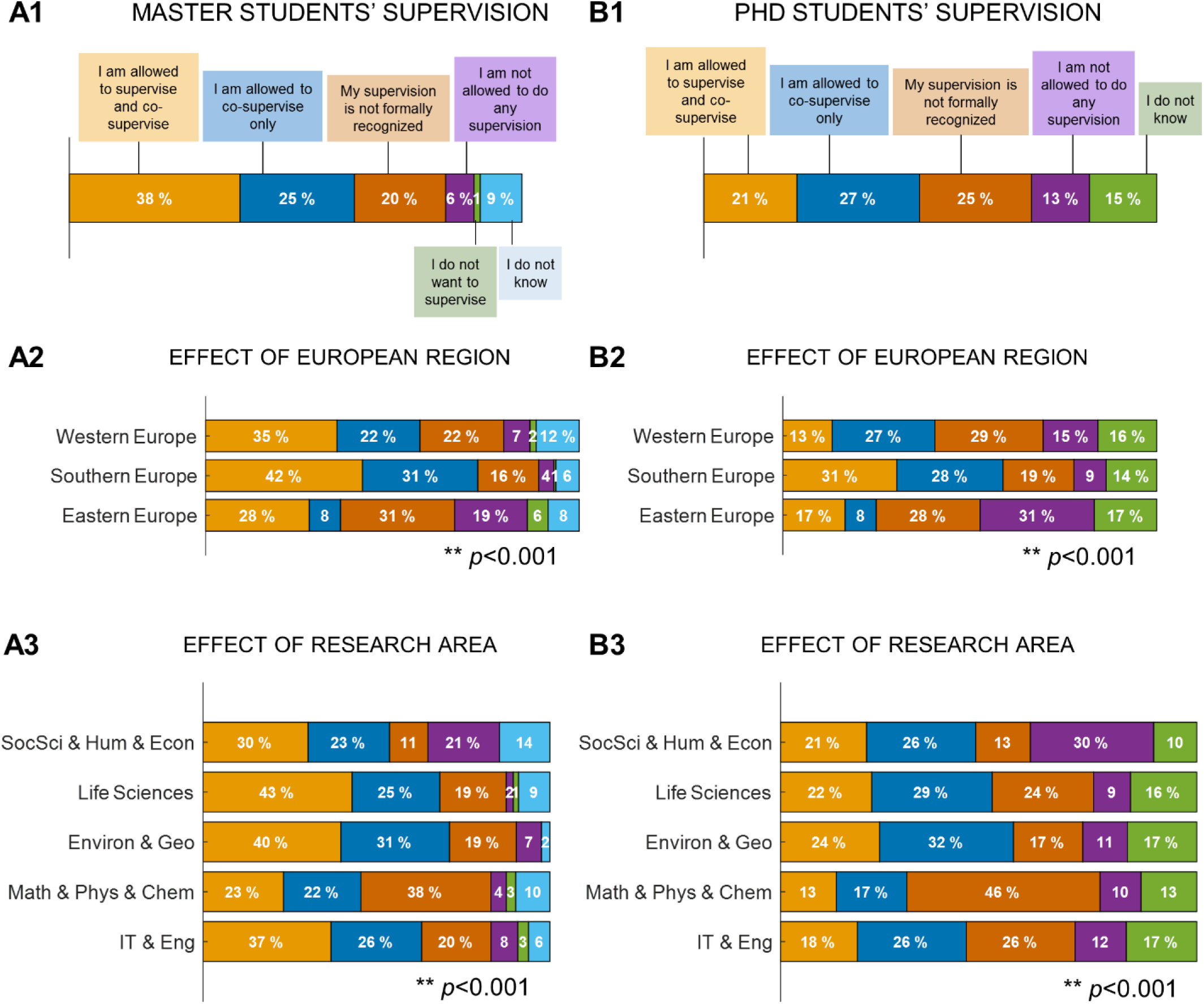
Graduate student supervision by postdoctoral researchers working in Europe.

##### Q31 Lead applicant for funding

Considering grant applications, 62% of the respondents said they collaborate with their supervisor in writing grants and 59% were allowed to apply for funding as a lead investigator of a research project. Collaboration in grant writing and application for funding was not dependent on the researcher gender. However, there was an effect of European region (*p* <.001) and of research area (*p* =.03). A higher percentage of researchers from Southern Europe collaborated in grant writing (Southern = 74%, Western = 53.6%, Eastern = 50%) and were allowed to apply for funding as a lead investigator of a research project (Southern = 71%, Western = 49.3 %, Eastern = 69.4%).

### 3.2 Postdoctoral researchers: Career perspectives and support

Career development is a crucial aspect of the postdoctoral researcher path. Here, we will describe the expectations researchers have regarding their career progress, and the perceived support that their institutions provide.

#### Q48 Long-term aspirations

In terms of future career, Figure 28A shows that **more than half of the respondents would like to work in academia**, either as a professor and researcher (52.1%), as a researcher only (15.3%), as a teacher only (.8%) or in science communication and research management (1.1%). Also, 13.2% of postdoctoral researchers would like to work in a research role outside academia and 6.2% in a non-research role outside academia. As much as 1.7% are still undecided and.6% would like to work in scientific publishing.

**Figure 28.**
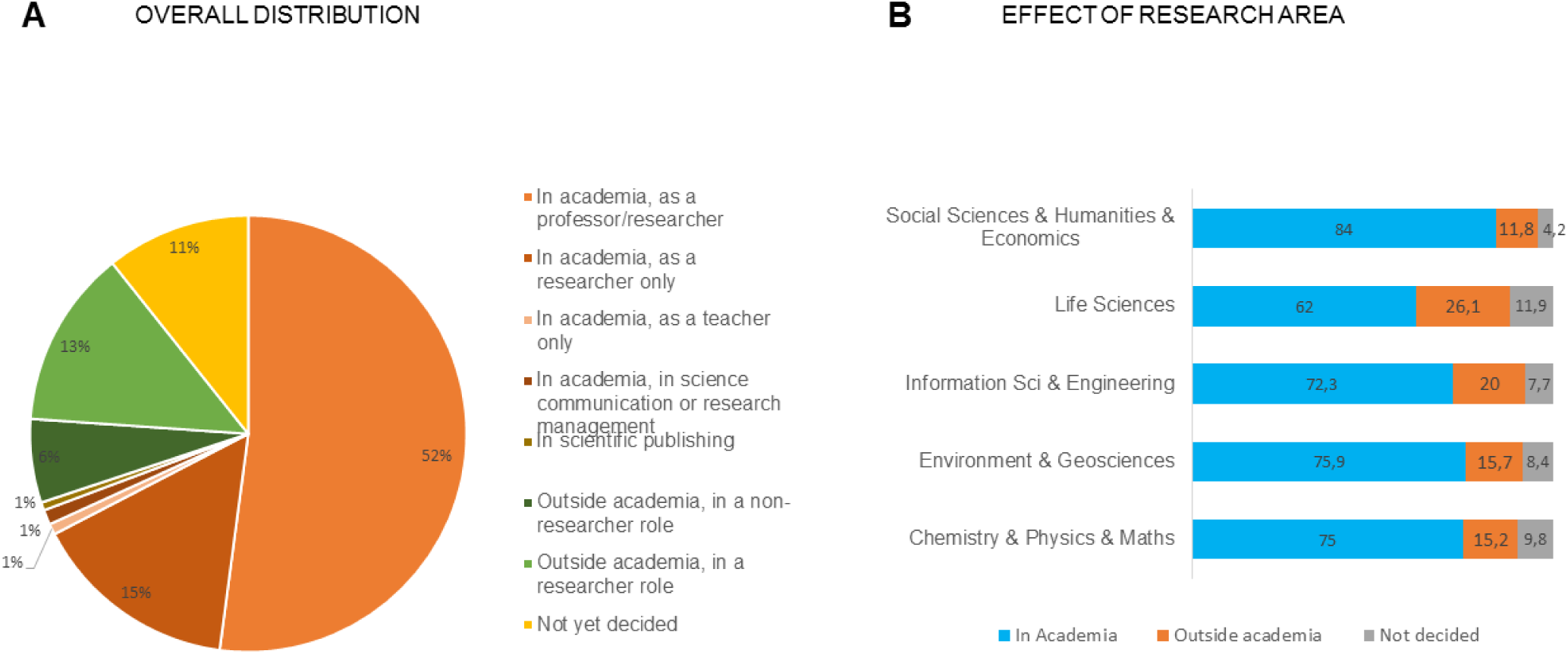
Future career prospects of postdoctoral researchers.

No significant differences in respondents’ future career prospects were found as a function of gender (*p* =.167) or as a function of European region (*p* =.592; data not shown). However, if the main research areas are considered, significant differences were found (*p* <.001): the majority of respondents wishes to continue in academia across all scientific domains (62% - 84%), but in LifeSci and in IT&Eng, a higher proportion of postdoctoral researchers see their future career outside academia (26.1% and 20% respectively, see Fig. 28B).

#### Q49 Clarity of career plan

Participants were asked to rate the clarity of their career plan in a Likert-type response scale from *1 = Not at all clear* to *5 = Very clear*. The majority of respondents positioned in the middle of the scale (2-4, 73.5%), with only 15% of respondents considering that their career plan is “*Not at all clear*” and 11.3% considering their career plan as “*Very clear*” (Fig. 29A). No significant differences were found, as function of gender (*p* =.579), European region (*p* =.180) or research area (*p* =.061).

**Figure 29.**
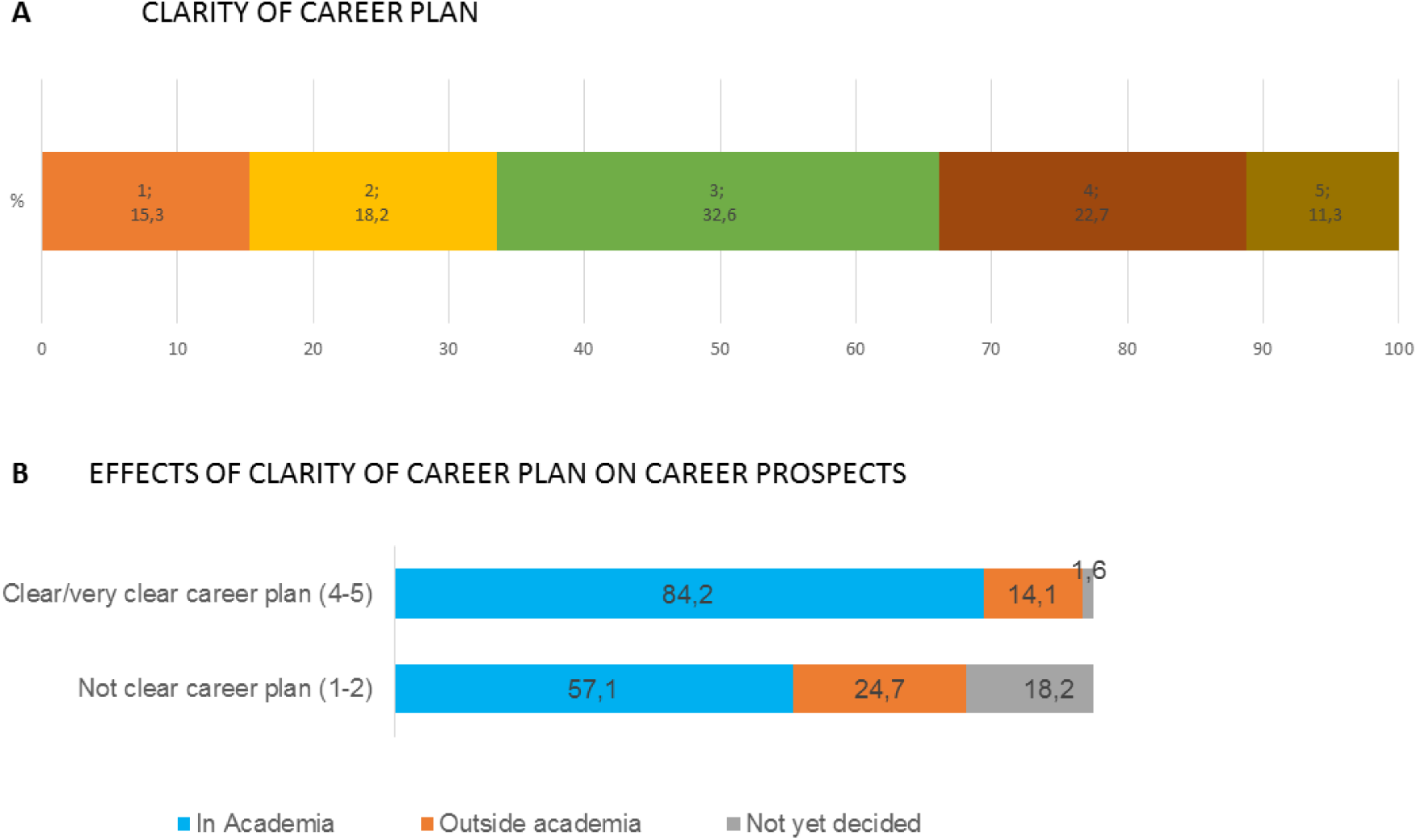
Clarity of career plan and career prospects.

When comparing the respondents who have a clear/very career plan with the ones who do not have a clear career plan (i.e., answered 1-2 to the question) (Fig. 29B), there were significant differences in their future career prospects (*p* <.001): a higher proportion of respondents with a clear/very clear career plan want to continue their career in academia (84.2% *vs.* 57.1%). Conversely, the respondents without a clear/very clear career plan desire with a higher frequency to leave academia (24.7% vs. 14.1%) and are more undecided (18.2%).

#### Q50 Currently job searching

The **majority of respondents was currently looking for jobs (57.3%)**, either in academia (24.3%), outside academia (8.6%) or both in and out of academia (24.4%), as seen in Figure 30A. The pattern was not different as a function of gender (*p* =.386) or European region (*p* =.292) (data not shown).

**Figure 30.**
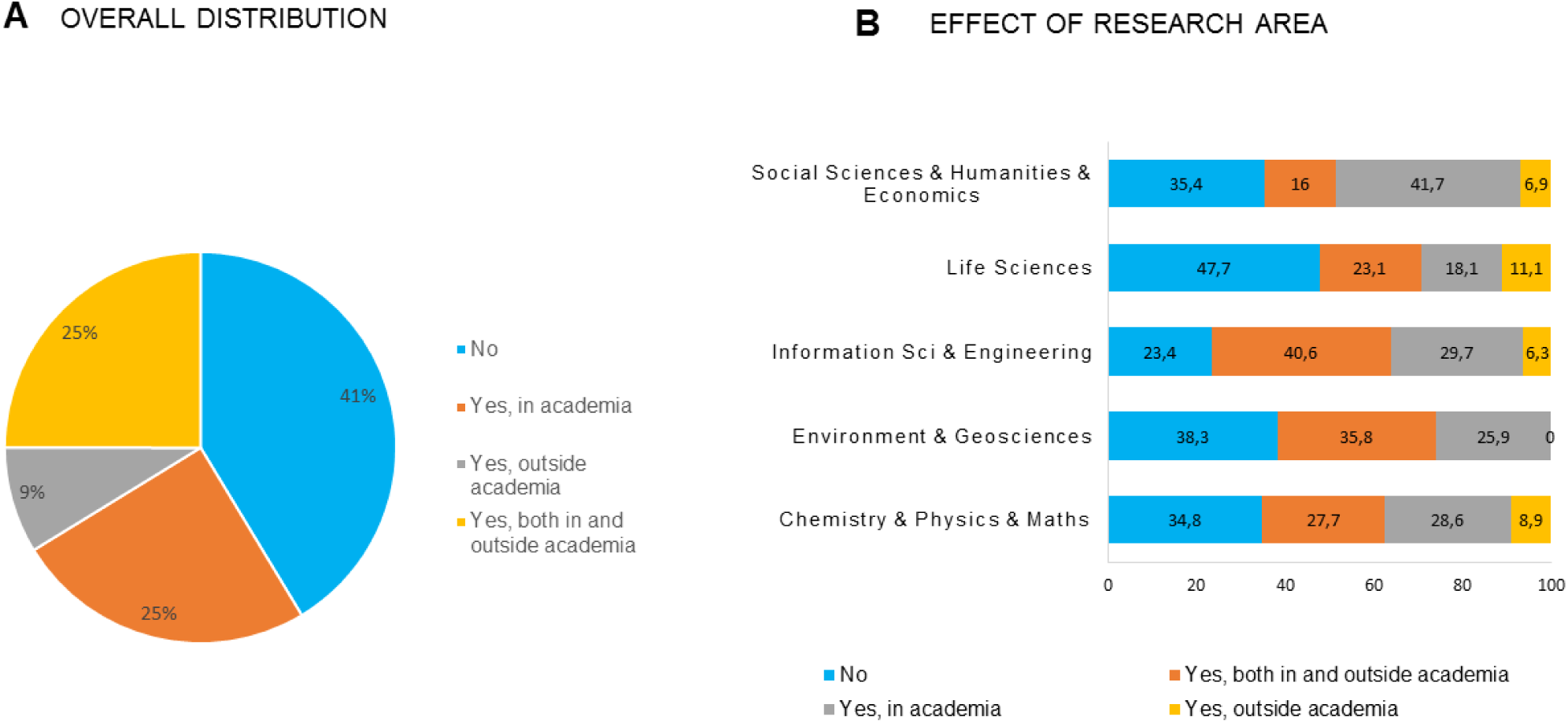
Current job search.

Significant differences emerged as a function of research area (*p* <.001; Fig. 30B): the proportion of job seekers was higher in IT&Eng (76.6%) and lower in LifeSci (52%). Except for SocSci&Hum&Econ where the majority of the job seekers were searching for jobs in academia (41.7% reported seeking a job in academia), in the remaining research domains, postdoctoral researchers were searching for jobs both in and outside of academia. Of note, in the Env&Geo research domain, there was no postdoctoral researchers searching for jobs exclusively outside academia.

Although the majority of respondents would like to work in academia (68.3%, cf. question 48), 21.1% of these were also looking for jobs outside academia. Conversely, of the ones who would like to work outside academia, 31.9% were searching for jobs both in and outside academia (Fig. 31).

**Figure 31.**
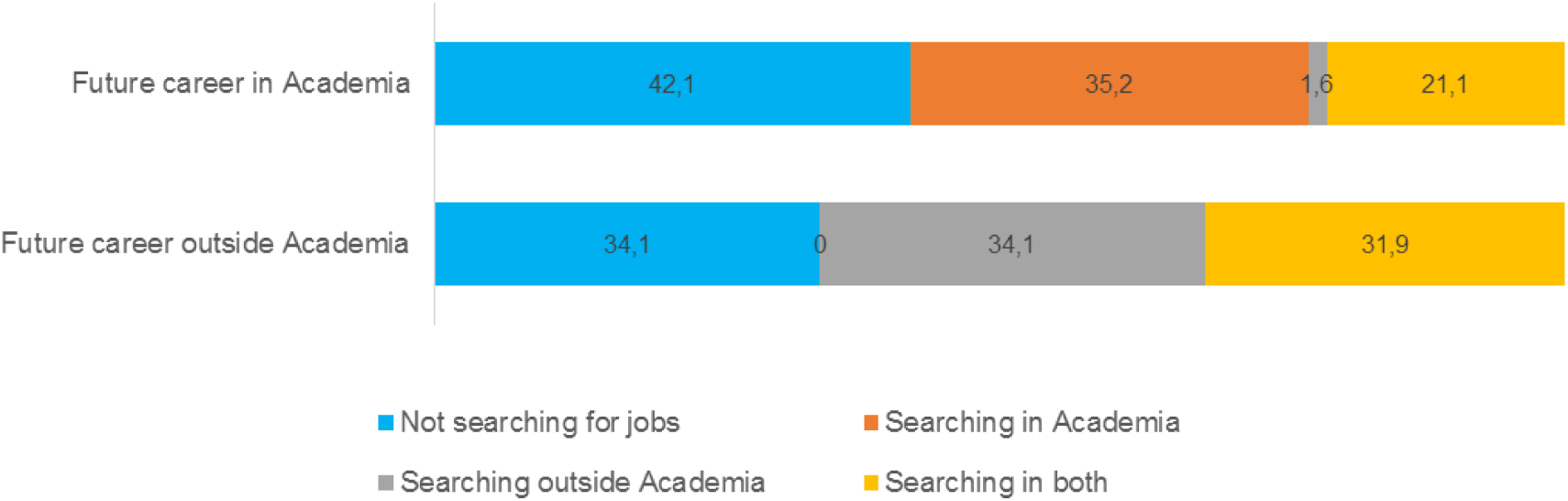
Type of job search carried out by researchers that would like to work, in the future, in or outside academia.

#### Q51-52 Office for postdoctoral researchers and satisfaction with support provided

In terms of institutional support, **only 38% of respondents identified that their institution had an office to support postdoctoral researchers**, while 62% reported that they did not have a support office or that they were not aware of its existence (Fig. 32A).

**Figure 32.**
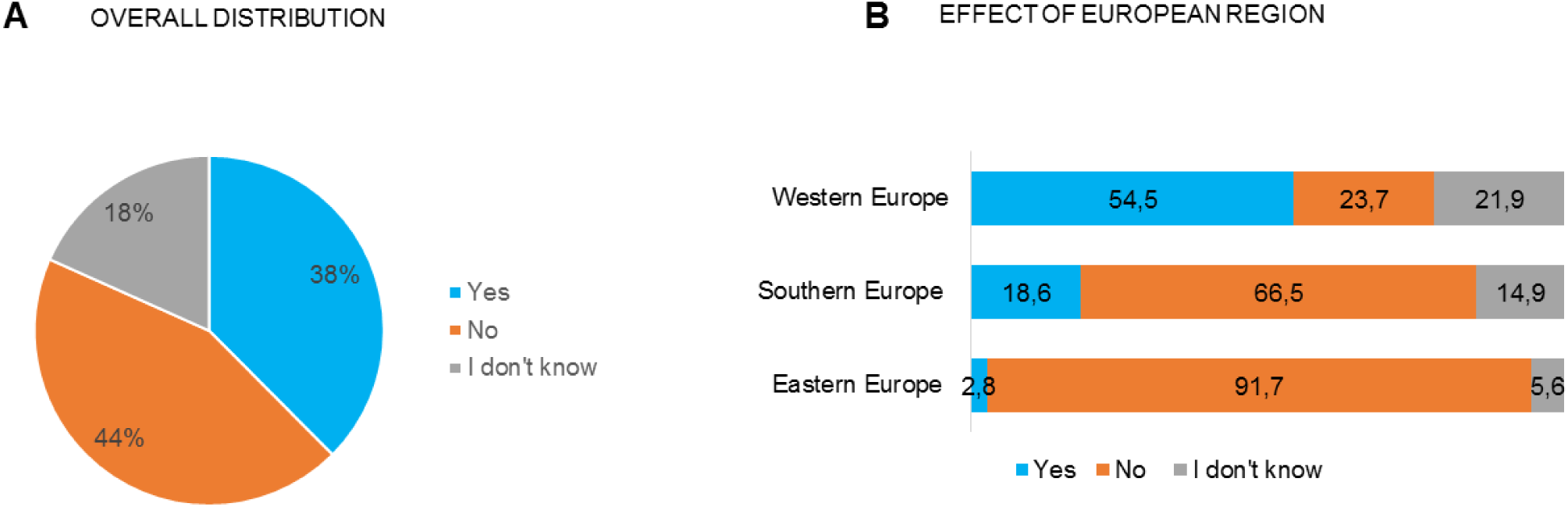
Existence of a Postdoctoral Office for support: Overall distribution and effect of European region.

Despite no significant differences being found as a function of gender (*p* =.06) or research area (*p* =.075), the pattern seems to differ across European region (*p* <.001): a significantly **higher proportion of respondents working in institutions from the Western Europe reported the existence of a postdoctoral office** (54.5%) when compared with Eastern and Southern European institutions (Fig. 32B).

Participants were asked to rate the satisfaction with the support they receive from their institutional office in a Likert-type response scale from *1 = Not satisfied at all* to *5 = Very much satisfied* (Fig. 33).

**Figure 33.**
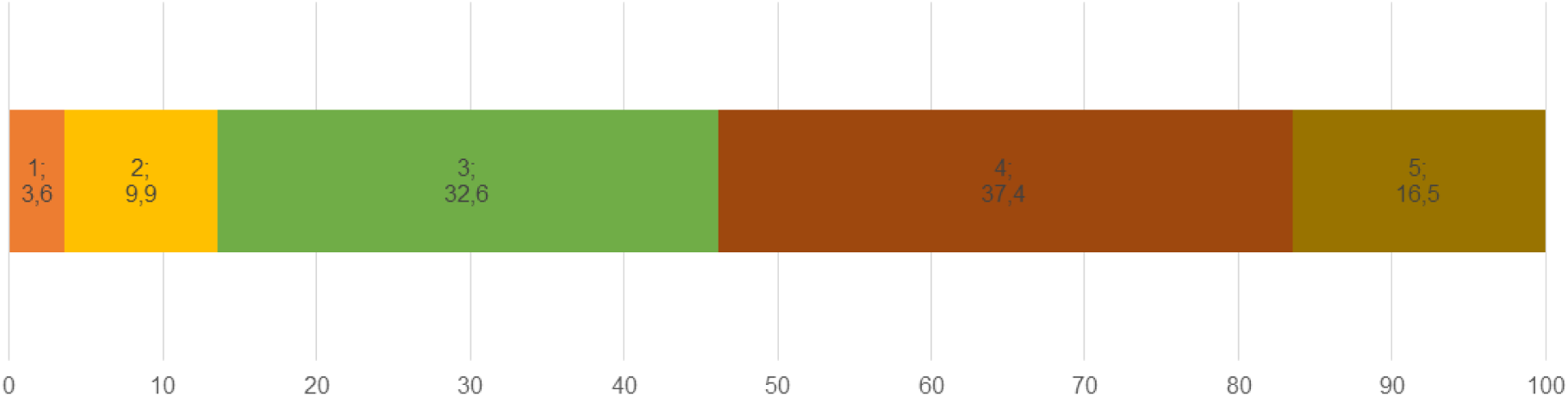
Perceived satisfaction with the postdoctoral office.

Of the participants who identified a support office in their institution, **more than 50% were satisfied (4) or very much satisfied (5) with the support received**. No significant differences were found as a function of gender (*p* =.806) and European region (*p* =.103) (data not shown). However, significant differences were found as a function of research domain (Fig. 34; *p* =.037): respondents in the field of Chem&Phys&Maths (28.9%) and IT&Eng (25.0%) are more frequently very much satisfied with the support they receive from the postdoctoral office.

**Figure 34.**
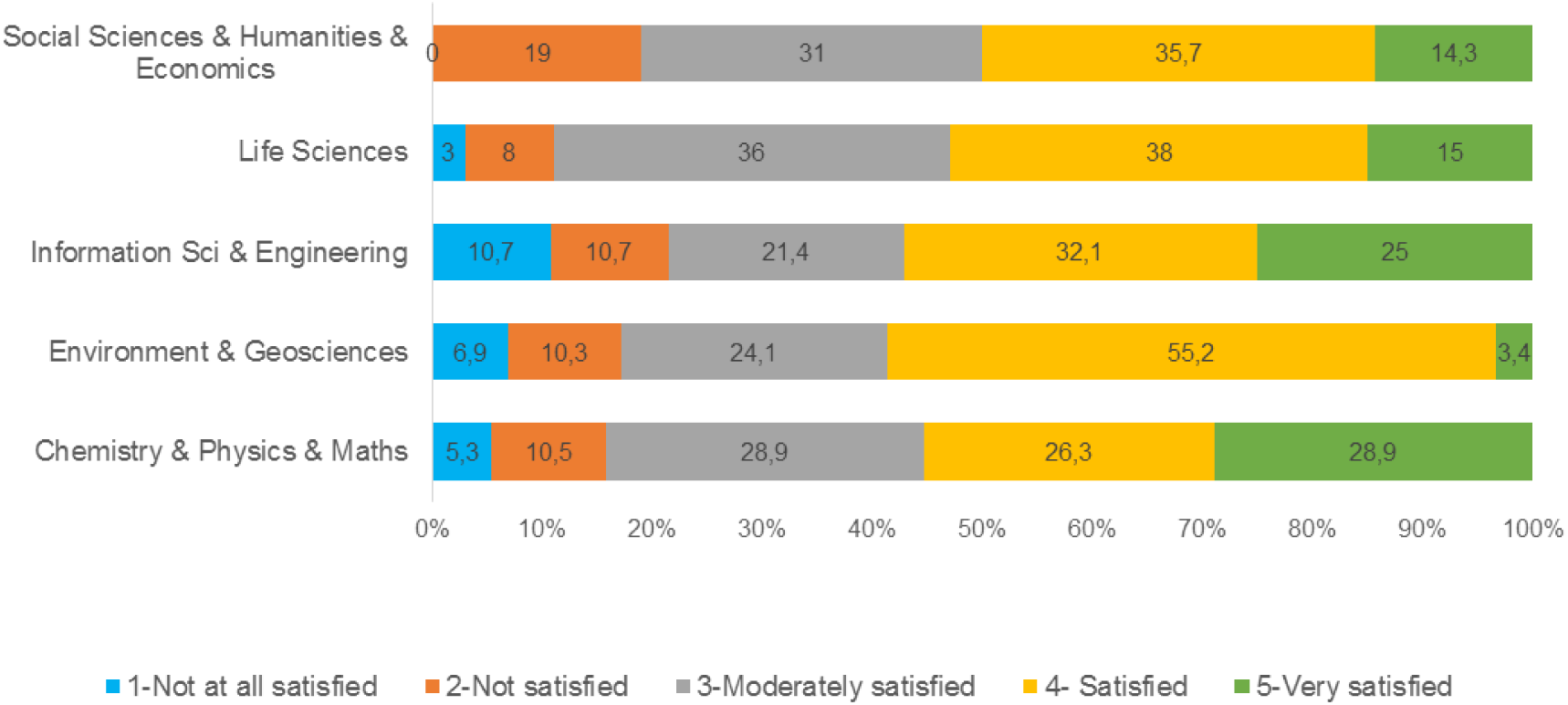
Perceived satisfaction with the postdoctoral office as a function of research area.

#### Q53 Perceptions on institutional services and support

Participants were asked about their perceptions regarding the personal and researcher development services offered by their institution (Fig. 35). In terms of transferable skills training, advice about career development opportunities and encouragement of personal and career development, between **45-48% of postdoctoral researchers agreed that their institutions provide or support those type of services**. Regarding mentorship programmes and assistance with conflict resolution that percentage drops to 27% and 26.6%, respectively.

**Figure 35.**
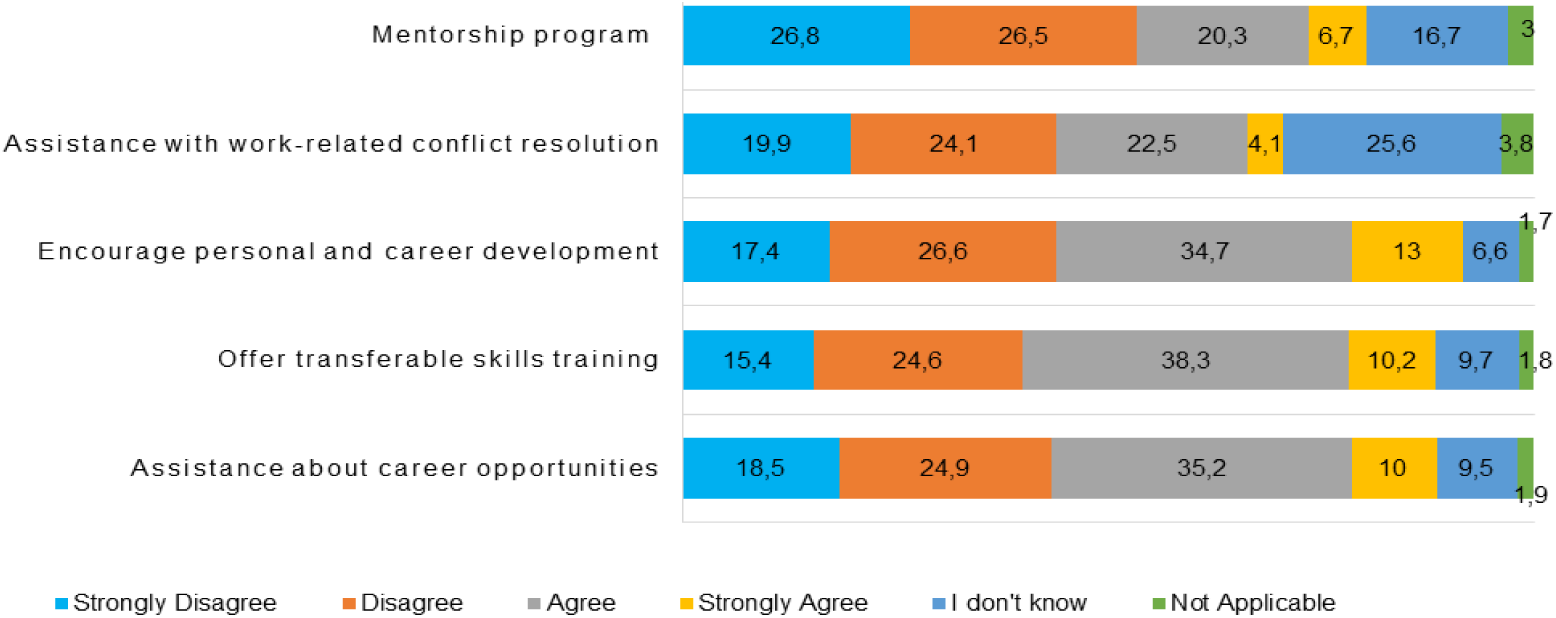
The extent to which researchers agree that the institution where they work provides each service.

We did not find significant gender differences regarding the extent to which researchers agreed that the institution where they work provides career development support services (data not shown). Differences were found concerning European region (*p* <.05) and research area (*p* <.05). As can be seen in Figure 36A, the proportion of postdoctoral researchers that report that their institutions provide support in career development is significantly higher in the Western Europe (35 to 65%) when compared to Eastern and Southern Europe.

**Figure 36.**
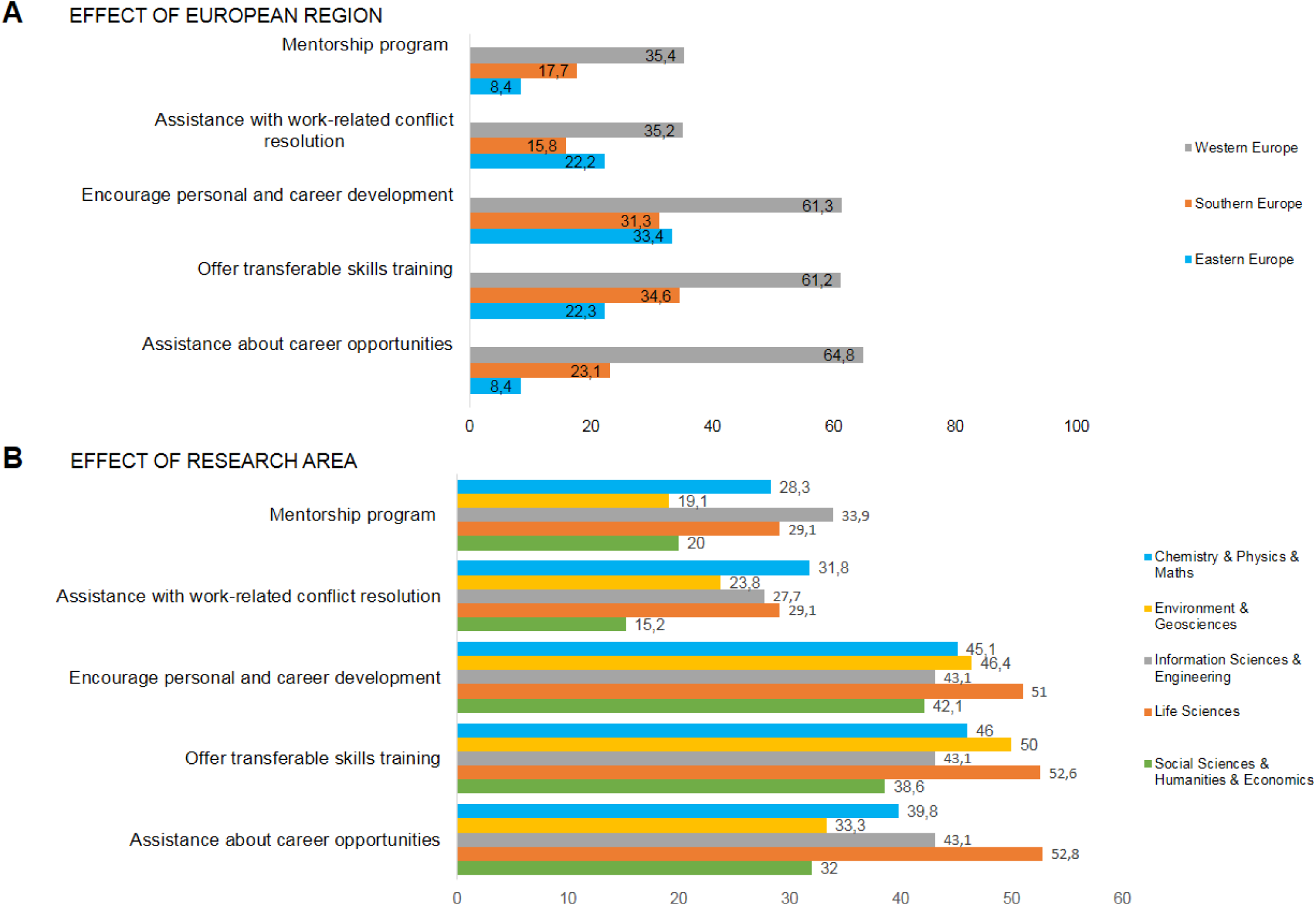
Proportion of postdoctoral researchers that agrees (agree or strongly agree) that their institution offers support in different services as a function of European region and Research Area

Also, considering the main research areas (Fig. 36B), researchers in SocSci&Hum&Econ reported not being provided with as much transferable skills training (38.6%) and assistance with work conflict resolution programmes (15.2%) as the other research areas (transferable skills training - more than half of the respondents in LifeSci and Env&Geo reported the institution offered such training; work conflict resolution: Chem&Phy&Maths, LifeSci and IT&Eng reported a proportion of about 30%). In terms of provision of advice about career opportunities, postdoctoral researchers in SocSci&Hum&Econ (32.0%) and Env&Geo (33.3%) reported lower support than the other main research areas.

#### Q54. Training undertaken

In terms of training, the training areas that postdoctoral researchers (1) have undertaken, (2) would like to take and (3) are not currently interested, are listed in Table 2. Noteworthy, there is a **high proportion of respondents (67%) who would like to undertake training in career management**, and this is congruent with the low proportion of respondents who have a very clear career plan (see Fig. 29A).

**Table 2.**
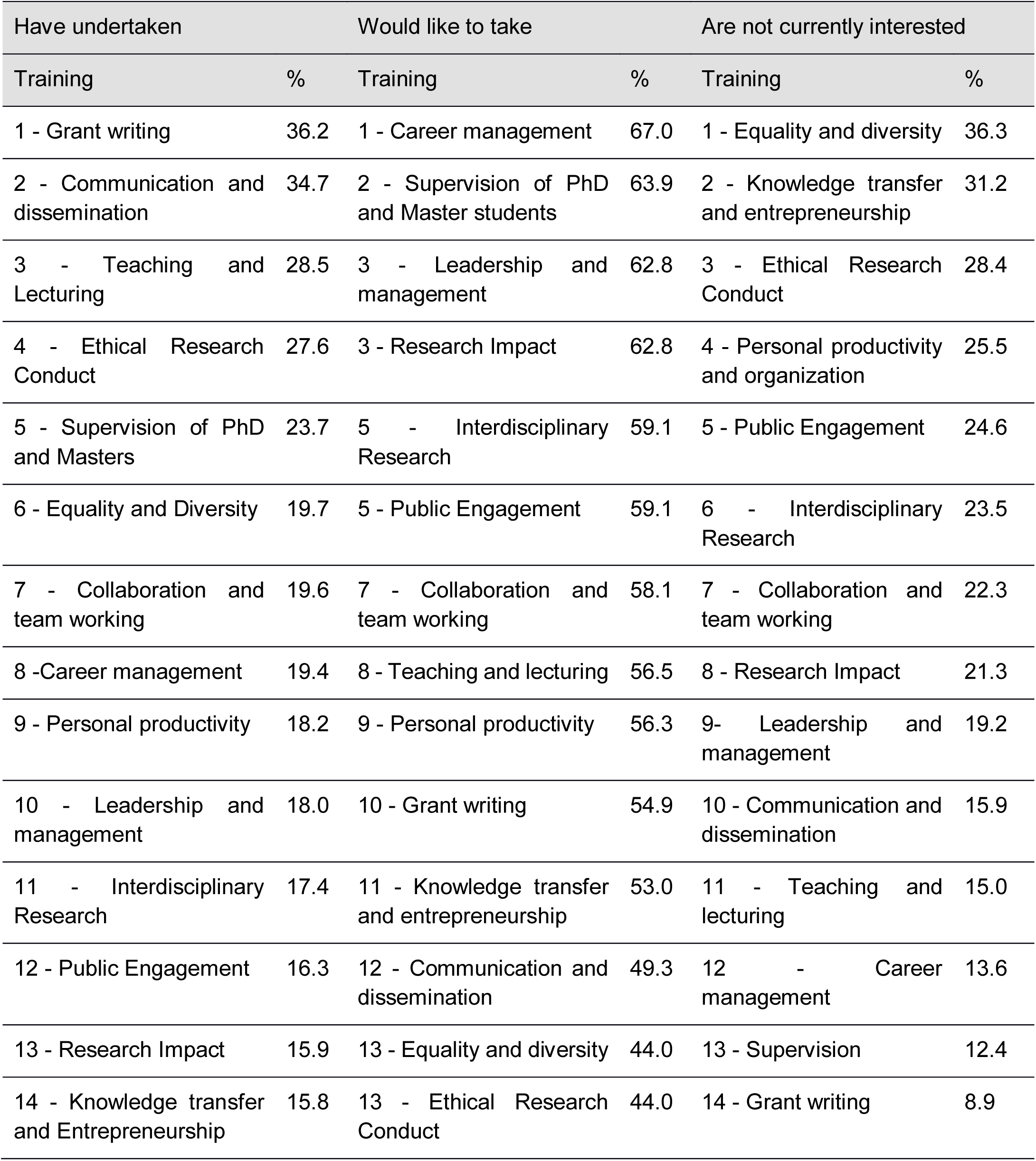
Training areas that postdoctoral researchers…

| Have undertaken |  | Would like to take |  | Are not currently interested |  |
| --- | --- | --- | --- | --- | --- |
| Training | % | Training | % | Training | % |
| 1 - Grant writing | 36.2 | 1 - Career management | 67.0 | 1 - Equality and diversity | 36.3 |
| 2 - Communication and dissemination | 34.7 | 2 - Supervision of PhD and Master students | 63.9 | 2 - Knowledge transfer and entrepreneurship | 31.2 |
| 3 - Teaching and Lecturing | 28.5 | 3 - Leadership and management | 62.8 | 3 - Ethical Research Conduct | 28.4 |
| 4 - Ethical Research Conduct | 27.6 | 3 - Research Impact | 62.8 | 4 - Personal productivity and organization | 25.5 |
| 5 - Supervision of PhD and Masters | 23.7 | 5 - Interdisciplinary Research | 59.1 | 5 - Public Engagement | 24.6 |
| 6 - Equality and Diversity | 19.7 | 5 - Public Engagement | 59.1 | 6 - Interdisciplinary Research | 23.5 |
| 7 - Collaboration and team working | 19.6 | 7 - Collaboration and team working | 58.1 | 7 - Collaboration and team working | 22.3 |
| 8 - Career management | 19.4 | 8 - Teaching and lecturing | 56.5 | 8 - Research Impact | 21.3 |
| 9 - Personal productivity | 18.2 | 9 - Personal productivity | 56.3 | 9- Leadership and management | 19.2 |
| 10 - Leadership and management | 18.0 | 10 - Grant writing | 54.9 | 10 - Communication and dissemination | 15.9 |
| 11 - Interdisciplinary Research | 17.4 | 11 - Knowledge transfer and entrepreneurship | 53.0 | 11 - Teaching and lecturing | 15.0 |
| 12 - Public Engagement | 16.3 | 12 - Communication and dissemination | 49.3 | 12 - Career management | 13.6 |
| 13 - Research Impact | 15.9 | 13 - Equality and diversity | 44.0 | 13 - Supervision | 12.4 |
| 14 - Knowledge transfer and Entrepreneurship | 15.8 | 13 - Ethical Research Conduct | 44.0 | 14 - Grant writing | 8.9 |

Regarding the training undertaken by postdoctoral researchers, the pattern was mostly similar across gender (Fig. 37). Nevertheless, significant gender differences were noted in the fields of equality and diversity (*p* =.008), ethical and research conduct (*p* =.026), and leadership and management (*p* =.001), with a higher percentage of women undertaking training in these areas.

**Figure 37.**
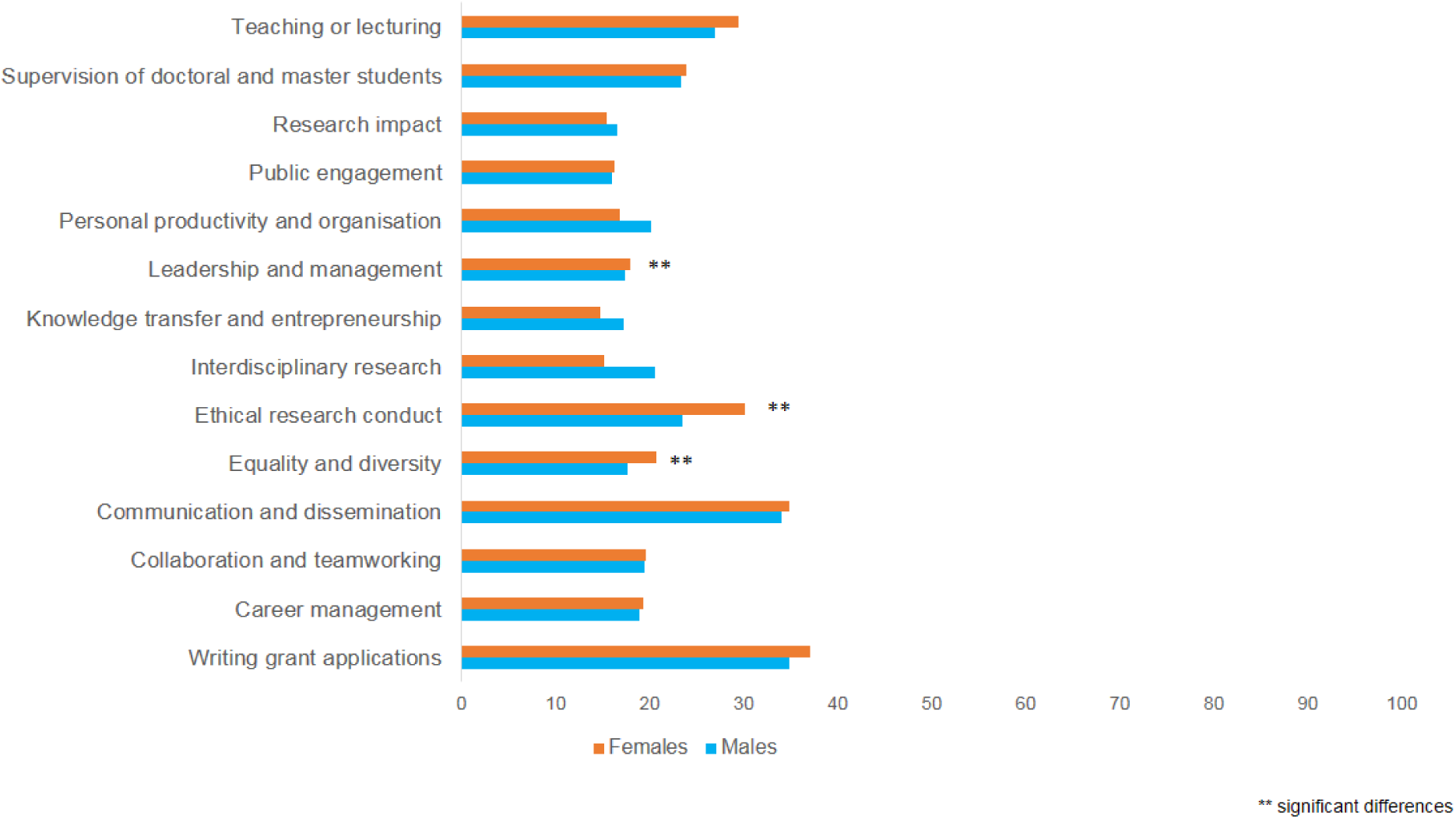
Proportion of postdoctoral researchers that reported having attended training in the different topics listed, as a function of gender.

Some significant differences were also found as a function of research area (Fig. 38). Specifically: a higher proportion of postdoctoral researchers in IT&Eng and LifeSci undertaken training in career management (*p* =.02); a higher proportion of postdoctoral researchers from Env&Geo have undertaken training in communication and dissemination (*p* =.027), public engagement (*p* <.001) and research impact (*p* =.001); a higher proportion of postdoctoral researchers from LifeSci undertaken training in Ethical research conduct (p <.001); and a higher proportion of postdoctoral researchers from Chem&Phys&Maths undertaken training in Supervision of PhD and Master students (*p* =.012) and Leadership and Management (*p* =.033).

**Figure 38.**
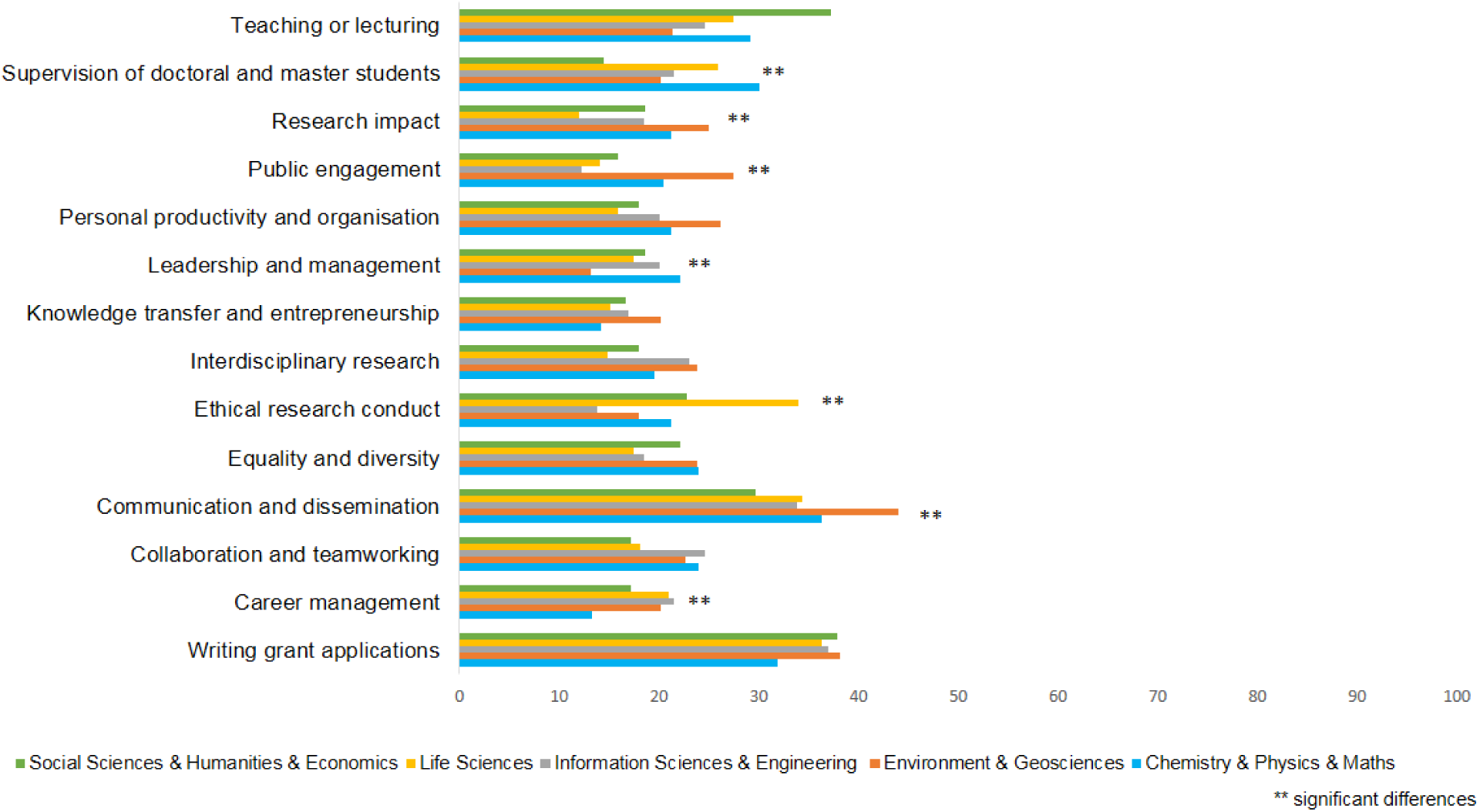
Proportion of postdoctoral researchers that reported having attended training in the different topics listed, as a function of research area.

Finally, significant differences were also found as a function of European region (Fig. 39). Postdoctoral researchers in Western Europe have undertaken with a higher frequency training in writing grant applications (*p* <.001; which may confer advantage in attracting funding), equality and diversity (p <.001), public engagement (*p* <.001), and supervision of PhD and master theses (*p* =.017), and with less frequency in knowledge transfer and entrepreneurship (*p* <.001). Respondents working in Eastern Europe more frequently had training in collaboration and team working (*p* <.001), and in teaching or lecturing (*p* =.028). Finally, respondents working in Southern Europe had more training in communication and dissemination (*p* =.001), and less training in career Management (*p* <.001).

**Figure 39.**
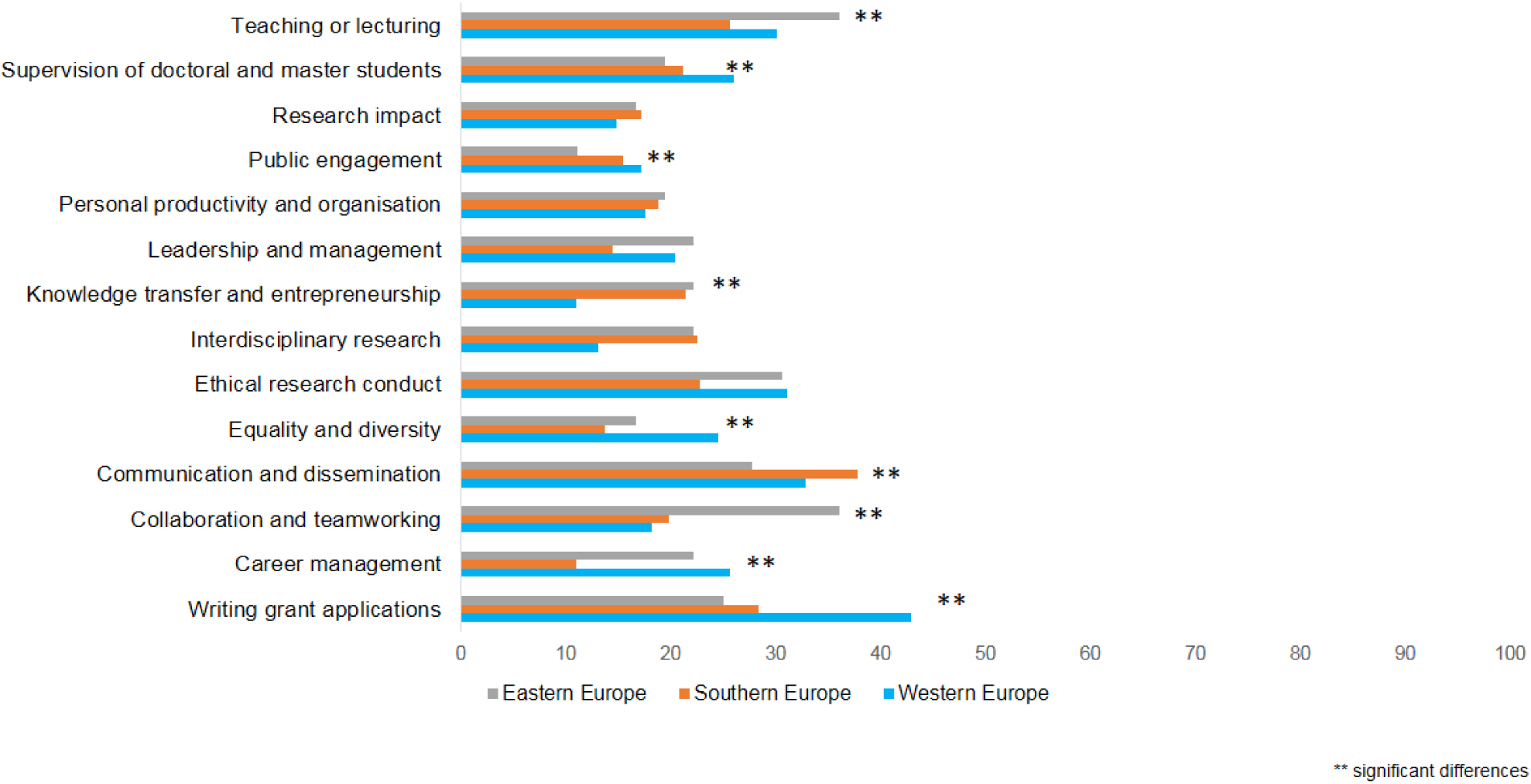
Proportion of postdoctoral researchers that reported having undertaken training in the different topics listed, as a function of European region.

### 3.3 Engagement with and within the postdoctoral community

#### 3.3.1 Institutional support

The well-being and achievement of a postdoctoral researcher can be influenced by multiple factors, including location of the institution, quality of the facilities, support provided to research - related activities, access to funds to support conference attendance or training activities, ethical conduct of the research group, level of scientific output of the host research organization and/or prospects for career progression. In this section, we provide an overview of how these factors are perceived by European postdoctoral researchers and how this changes across Europe, research area, and researcher’s gender.

##### Q43-45 Support provided institutionally on facilities and funding

When asked about the quality of the facilities at the institution, the quality of the support to research activities and the satisfaction with access to funds to support conference attendance or training activities, **at least half of the respondents reported being satisfied:** 74% were very satisfied or satisfied with the facilities, 52% were very satisfied or satisfied with access to supporting funds and 58% were very satisfied or satisfied with the support to research (Figure 40).

**Figure 40.**
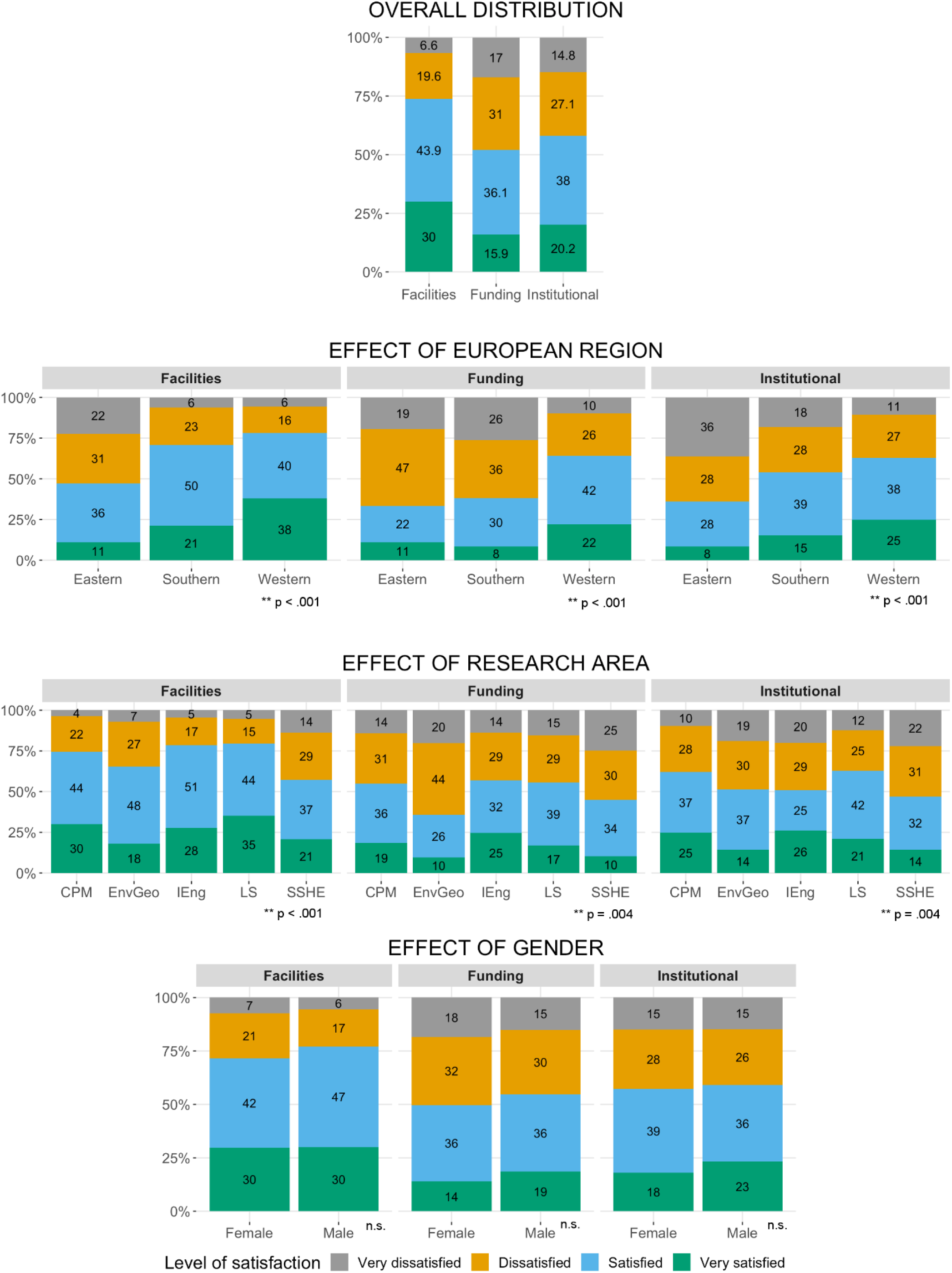
Levels of satisfaction with the quality of facilities, institutional support and funding as a function of European region, research area and gender.

When looking at the rating of the facilities, quality of the support to research activities and access to funding as a function of European region, research area and gender, significant differences were observed as follows:

- There was a significant relationship between European region and the satisfaction with the support provided, across the three categories. Postdoctoral researchers in institutions in Southern and Eastern Europe were in general less satisfied than the ones in Western Europe (*p* <.001);
- There was also a significant relationship between the area of research and satisfaction of postdoctoral researchers with the support provided (*p* <.001; *p =.*0038, *p* =.0036, for facilities, funding and institutional). Researchers in the Env&Geo and SocSci&Hum&Econ areas were less satisfied than researchers from the other fields (Chem&Phys&Maths, IT&Eng and LifeSci), across the three support areas;
- There was no effect of gender (*p* =.07; *p* =.60; *p* =.15, for facilities, funding and institutional support, respectively).

When assessing satisfaction by Institution, responses were aggregated as ‘Dissatisfied’ (‘Very dissatisfied’ and ‘Dissatisfied’) and ‘Satisfied’ (‘Very Satisfied’ and ‘Satisfied’). Of the 13 institutions with at least 10% of the survey responses (N = 145 to 9), the **top two** ranked institutions where postdoctoral researchers rated all these three aspects as satisfactory were the **Max Planck Institute for Molecular Cell Biology in Germany, and the Genetics and Centre for Genomic Regulation in Spain** (Fig. 41). The institution which rated lower for the average of all categories was the **Consejo Superior de Investigaciones Cientificas in Spain**.

**Figure 41.**
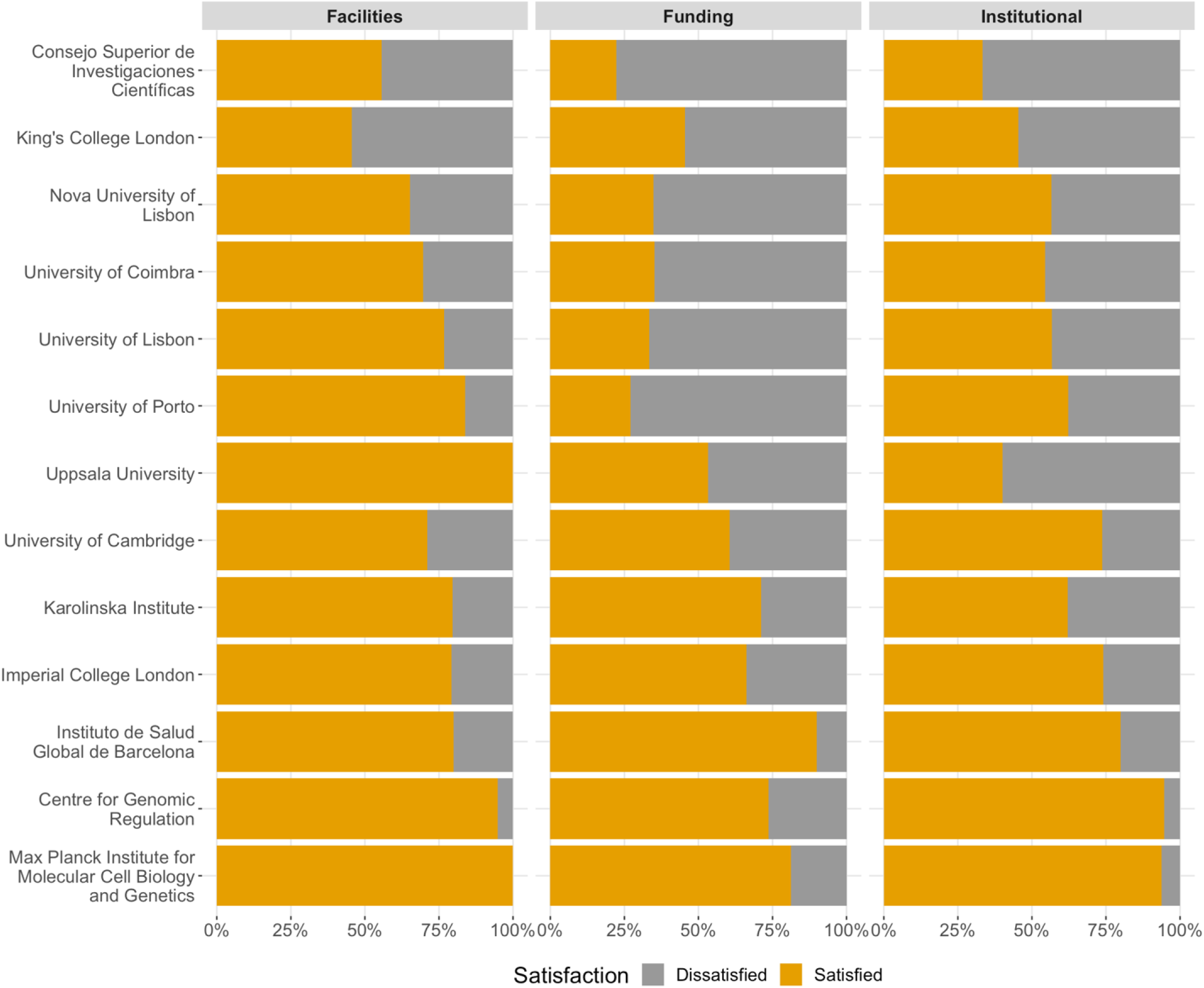
Ranking of institutions for satisfaction with support provided.

##### Q46. Clear duties and rights

In relation to how well defined were the rights and duties of the postdoctoral researcher at the beginning of the contract or fellowship, **69% reported that the rules were clearly or partially defined and 29% indicated that the rules were unclear** (Fig. 42). A small significant difference was found as a function of European region, with only 22 % of postdoctoral researchers in Southern Europe reporting that their duties and rights were clearly established (*p* =.02). No significant differences were found in the clarity of the rights and duties of the postdoctoral researchers as a function of research area (*p* =.14) or gender (*p* =.27). This significance was calculated using aggregated responses: ‘Defined’ for answers ‘Clearly defined rules’ and ‘Partially defined rules’, and ‘Not defined’ for answers ‘I don’t know’ and ‘Undefined’.

**Figure 42.**
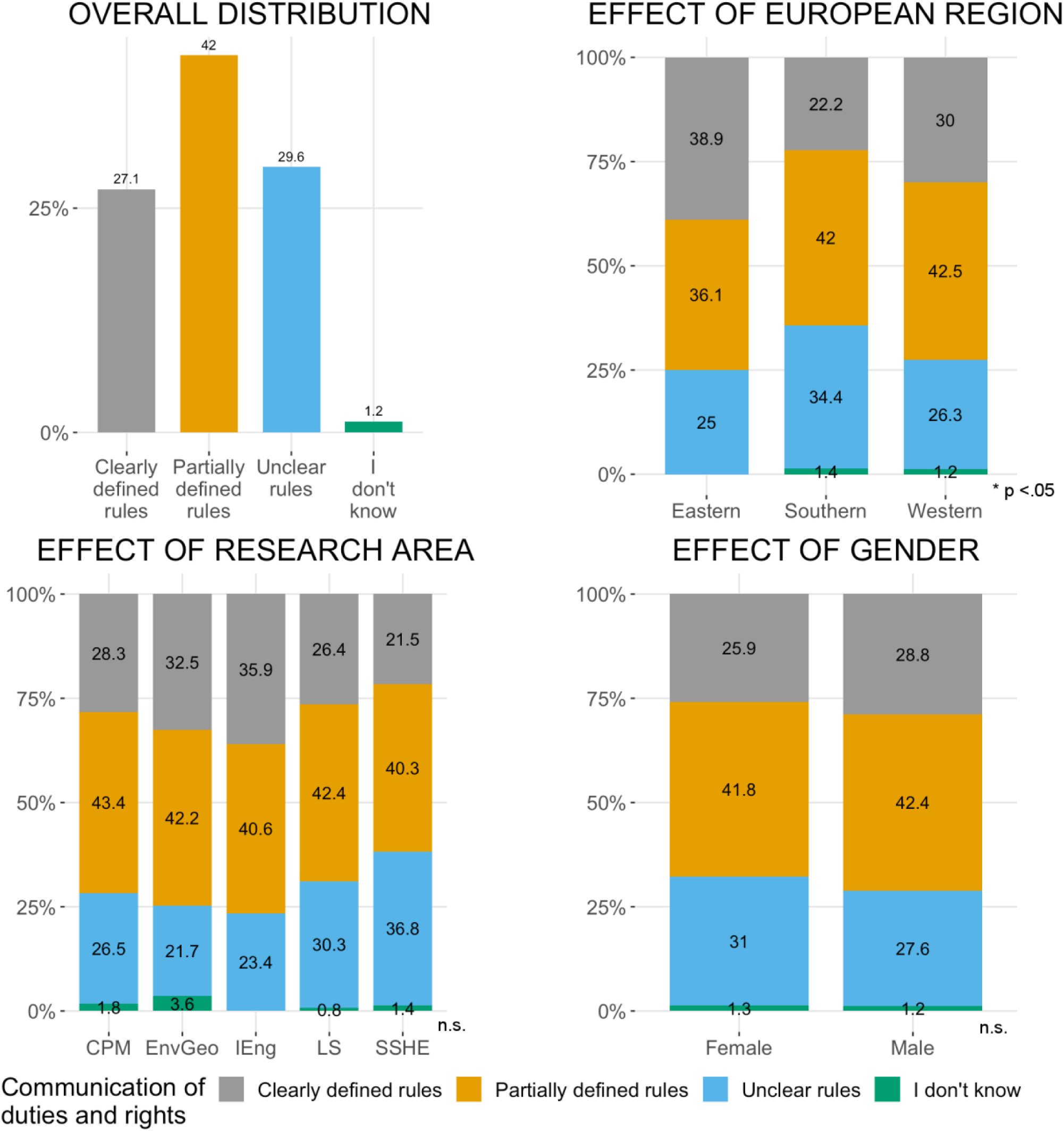
Communication of duties and rights as a function of European region, research area and gender.

##### Q47. Representation of postdoctoral researchers in the institution’s governance

In terms of postdoctoral representation in the management bodies of the institutions, 28% did not know if the representation existed and **33% stated that the postdoctoral researchers were not represented** (Fig. 43).

**Figure 43.**
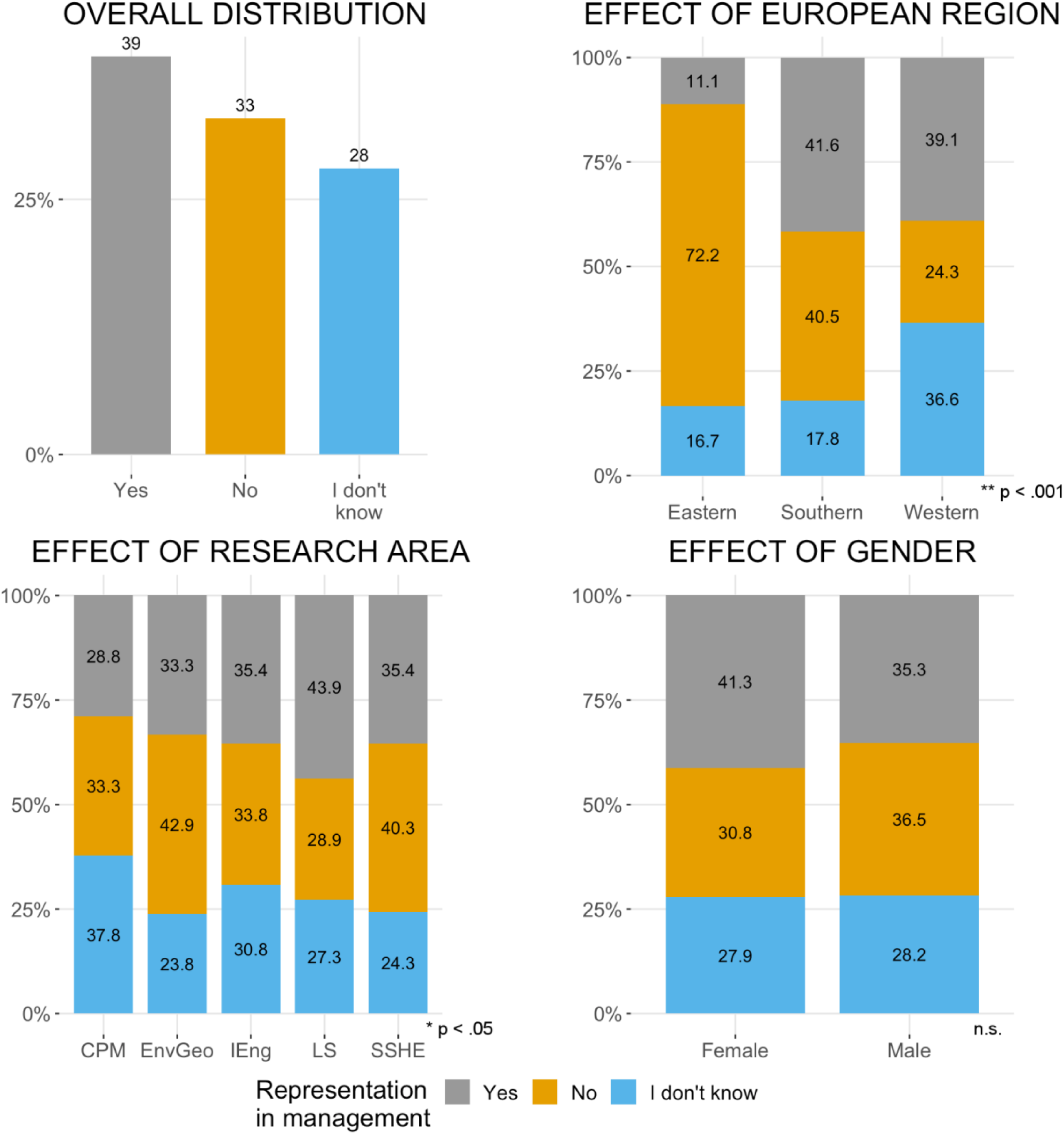
Representation of postdoctoral researchers in management bodies as a function of European region, research area and gender.

Significant differences were observed for European region (*p* <.001) and research area (*p* =.011). Representation was more common in institutions in Western and Southern Europe than in Eastern Europe (39% and 42%, respectively, versus 11%). A higher proportion of postdoctoral researchers in Western Europe did not know if postdoctoral researchers were represented (37% vs 18% in Southern and 17% in Eastern Europe). Postdoctoral researchers in LifeSci were the most likely to be represented in governance bodies (44%). No significant differences were found for gender (*p* =.14).

Interestingly, there was a significant relationship between the lack of representation of postdoctoral researchers in governance bodies and how well their rights and duties were communicated (*p* =.004; Fig. 44). The **lack of clear rules was positively correlated with the lack of representation of postdoctoral researchers in the institutional managing bodies**, i.e. postdoctoral researchers in institutions where there was no representation were 12% more likely to report that their rights and duties were unclear.

**Figure 44.**
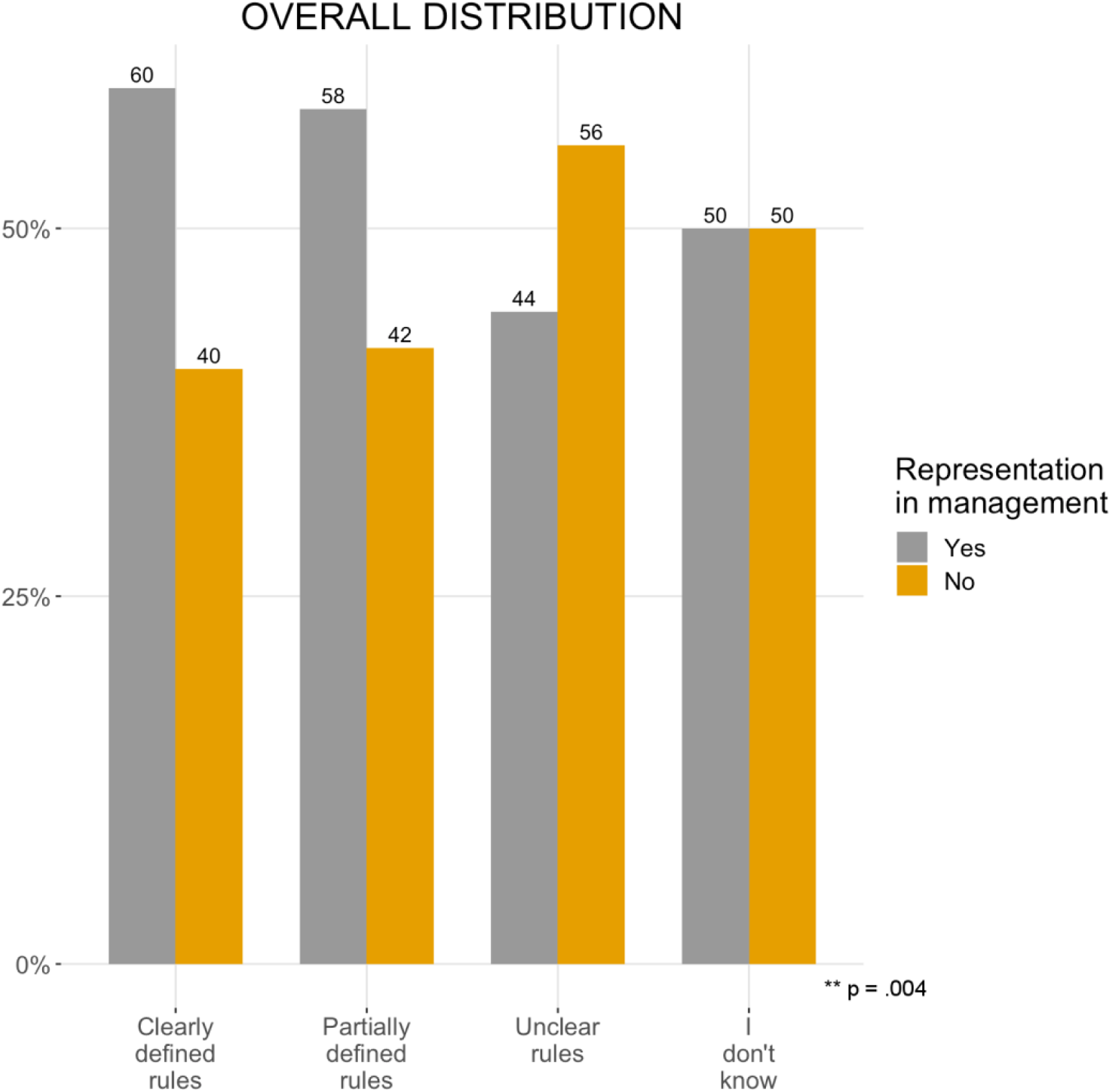
Duties and rights of postdoctoral researchers as a function of being represented in the institution’s governance.

#### 3.3.2 Community engagement

With an ever increasing number of people completing PhDs and staying in research as postdoctoral researchers, the need to define, potentiate and defend their rights and duties has become pivotal in many institutions and countries. Hence, at a global level, the existence of institutional or national postdoctoral researchers associations or unions is often perceived as a positive step and can reflect a mature and healthy academic system.

##### Q55 Union membership

Among the participants of this survey, 75% were not members of a union (Fig. 45). There was no difference in this aspect across European region (*p* =.16) or gender (*p* =.28). There was, however, a significant difference as a function of research area (*p* =.012), with postdoctoral researchers in Chem&Phys&Maths being less likely to be members of a union (only 9.9%).

**Figure 45.**
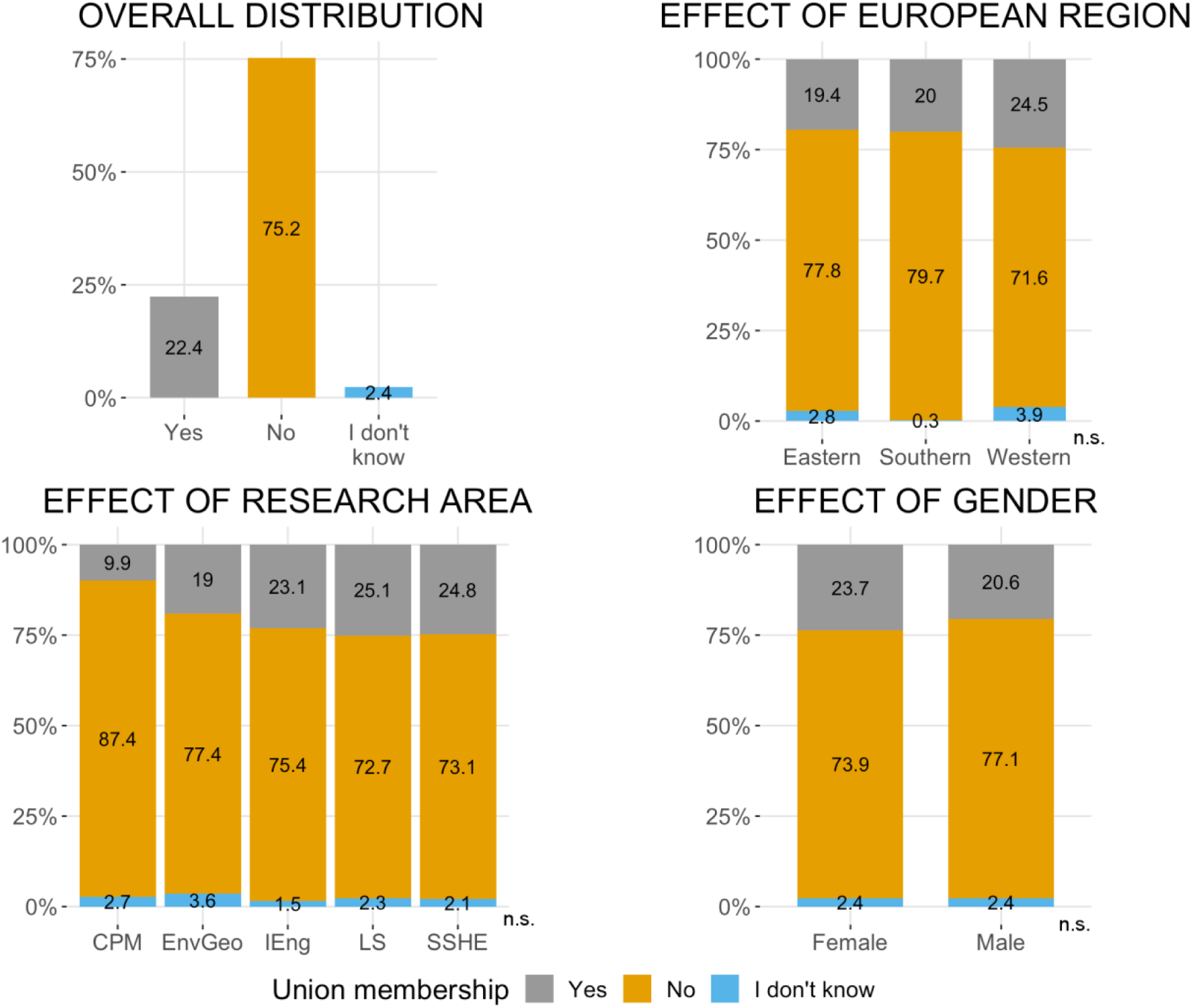
Union membership as a function of European region, research area and gender.

##### Q56 Postdoctoral association at institution

Among the participants of this survey, 53% reported that their institution had a postdoctoral association (Fig. 46). There was a significant effect of European region (*p* <.001) and research area (*p* <.001). It is more common for institutions in Southern and Western Europe to have a postdoctoral association than institutions from Eastern Europe (50% and 59%, respectively, versus 8% for Eastern Europe). Regarding research area, institutions employing postdoctoral researchers working in the LifeSci were more likely to have a postdoctoral association (69%). There was no effect of researcher’s gender (*p* =.38).

**Figure 46.**
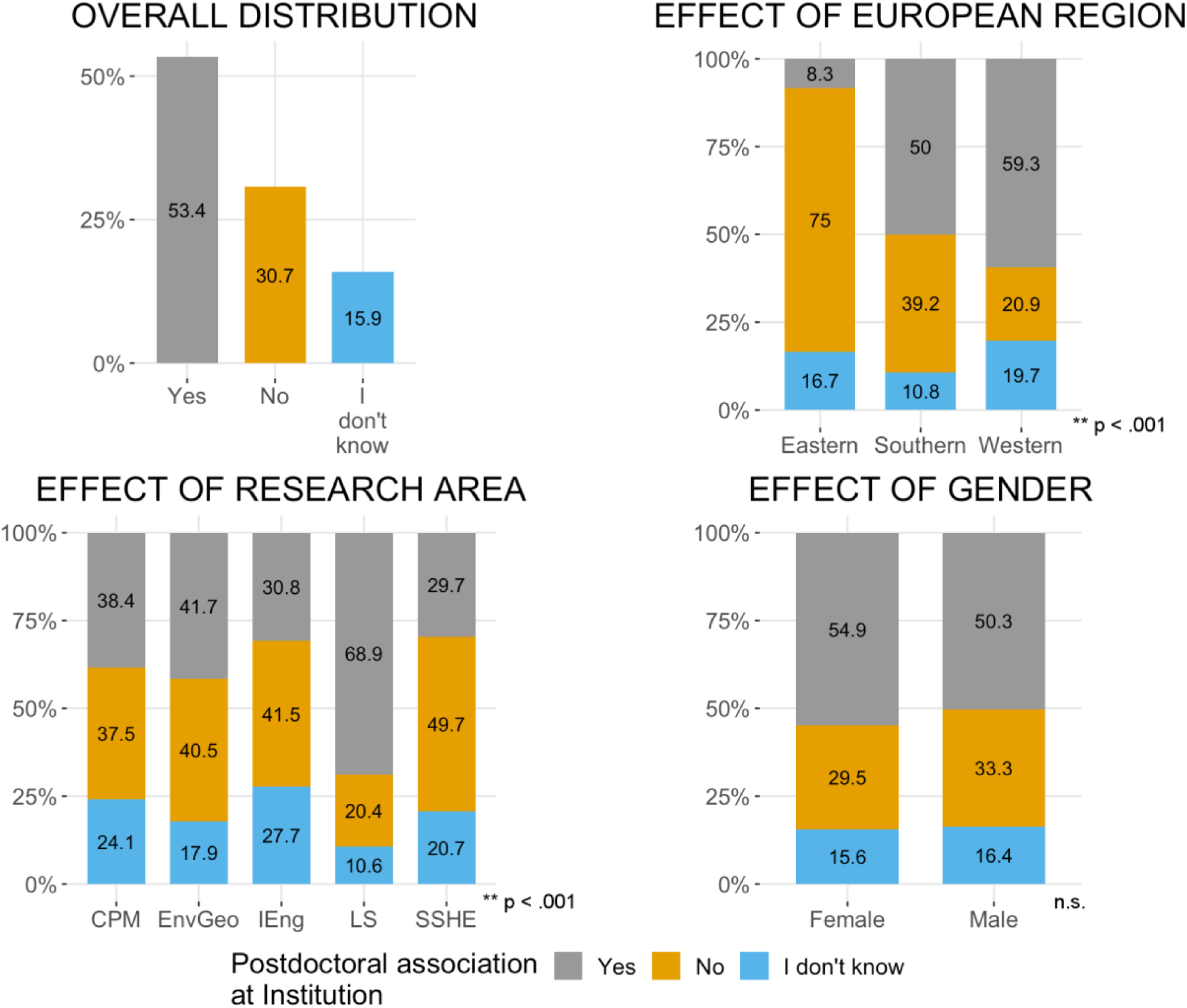
Existence of postdoctoral association at institution, as a function of European region, research area and gender.

##### Q57. Member of a postdoctoral association

Of the postdoctoral researchers that reported that their institution had a postdoctoral association, **the majority (61%) were members of their postdoctoral association** (Fig. 47). However, there was a significant difference depending on which research area they worked in (*p* <.001). Postdoctoral researchers in the Chem&Phys&Maths, and IT&Eng, were less likely to be members of a postdoctoral association compared to postdoctoral researchers in the LifeSci or in the SocSci&Hum&Econ (33% and 32%, versus 65% and 74%, respectively). No significant difference was apparent for European region or gender (*p* =.26 and *p* =.50, respectively).

**Figure 47.**
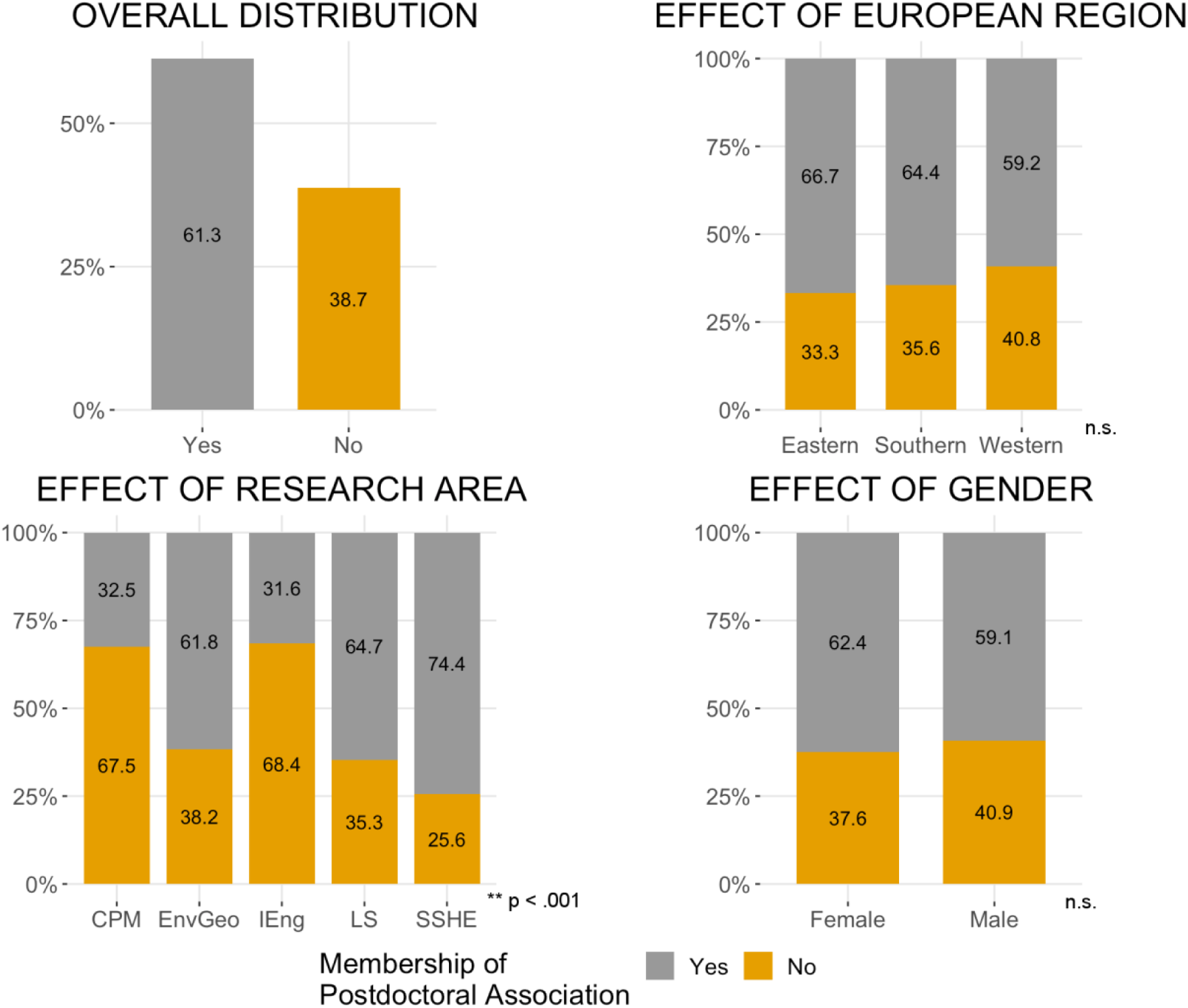
Membership of Postdoctoral Association, as a function of European region, research area and gender.

##### Q58 Being informed of postdoctoral association at institution

Almost **half of all postdoctoral researchers got to know there was a postdoctoral association at their institution through other postdoctoral researchers (49%)** (Fig. 48). Sharing this information via the institution (through induction events or other information) was the second most frequent way of communicating this to postdoctoral researchers (37%). There was a significant difference to how this information was shared depending on the country of work (*p* < .001). Postdoctoral researchers in Eastern Europe did not receive this information via their institution, other than at an induction event. Postdoctoral researchers in Southern Europe were more likely than postdoctoral researchers in Western Europe to receive this information via colleagues than at an induction event (6% vs 16% for induction, 61% vs 41% for colleagues). There were no differences as a function of gender (*p* =.51)

**Figure 48.**
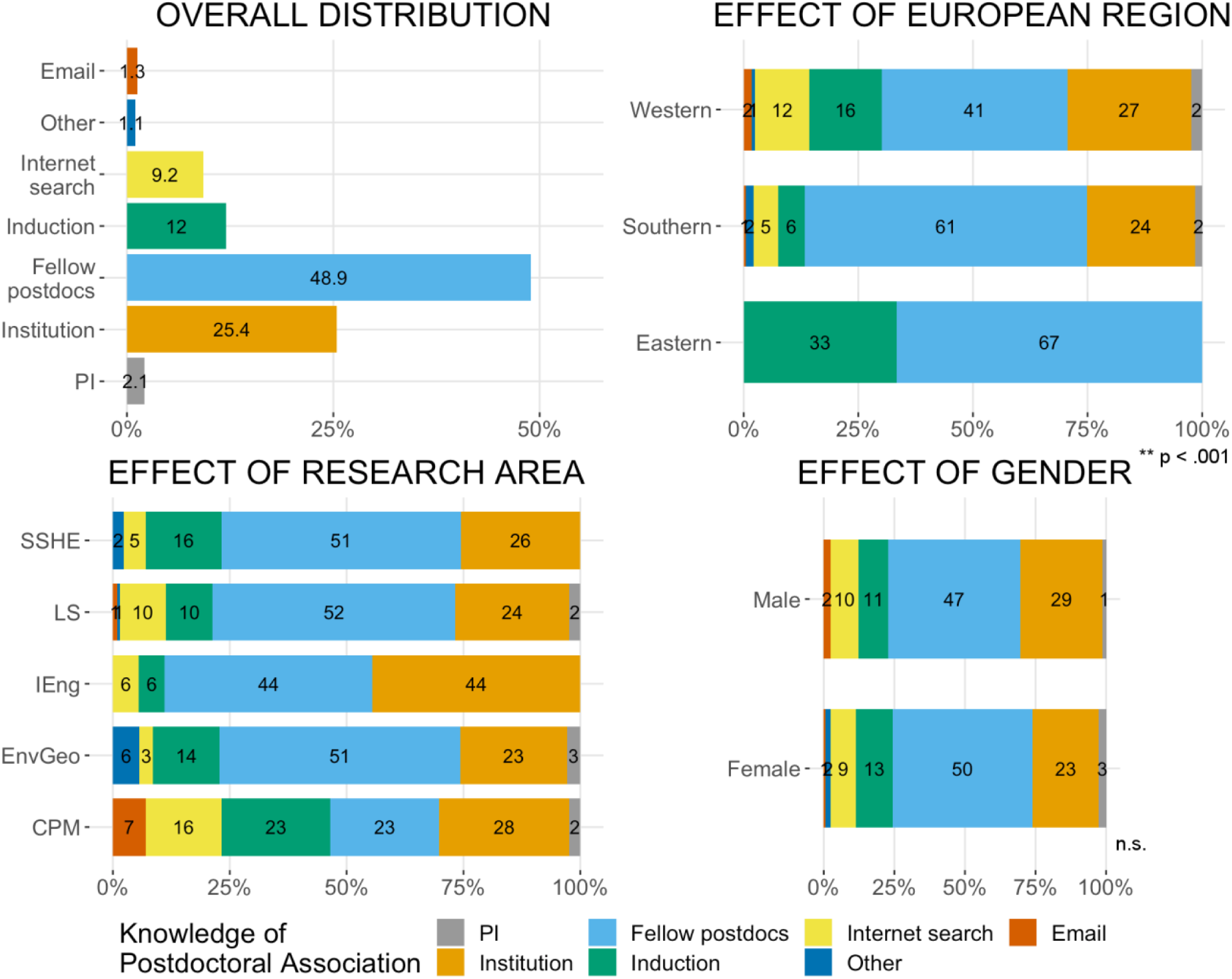
How is the existence of a postdoctoral association communicated to postdoctoral researchers, as a function of European region, research area and gender.

##### Q59. Scope of a postdoctoral association

Regarding what type of events postdoctoral researchers would like postdoctoral associations to organise that are currently lacking, there was a **clear preference for mentoring schemes and career development events (65% and 72%)** (Fig. 49). Social events, networking events, workshops, seminars or discussion panels on career paths were also of interest to postdoctoral researchers, but postdoctoral associations already organise this kind of events.

**Figure 49.**
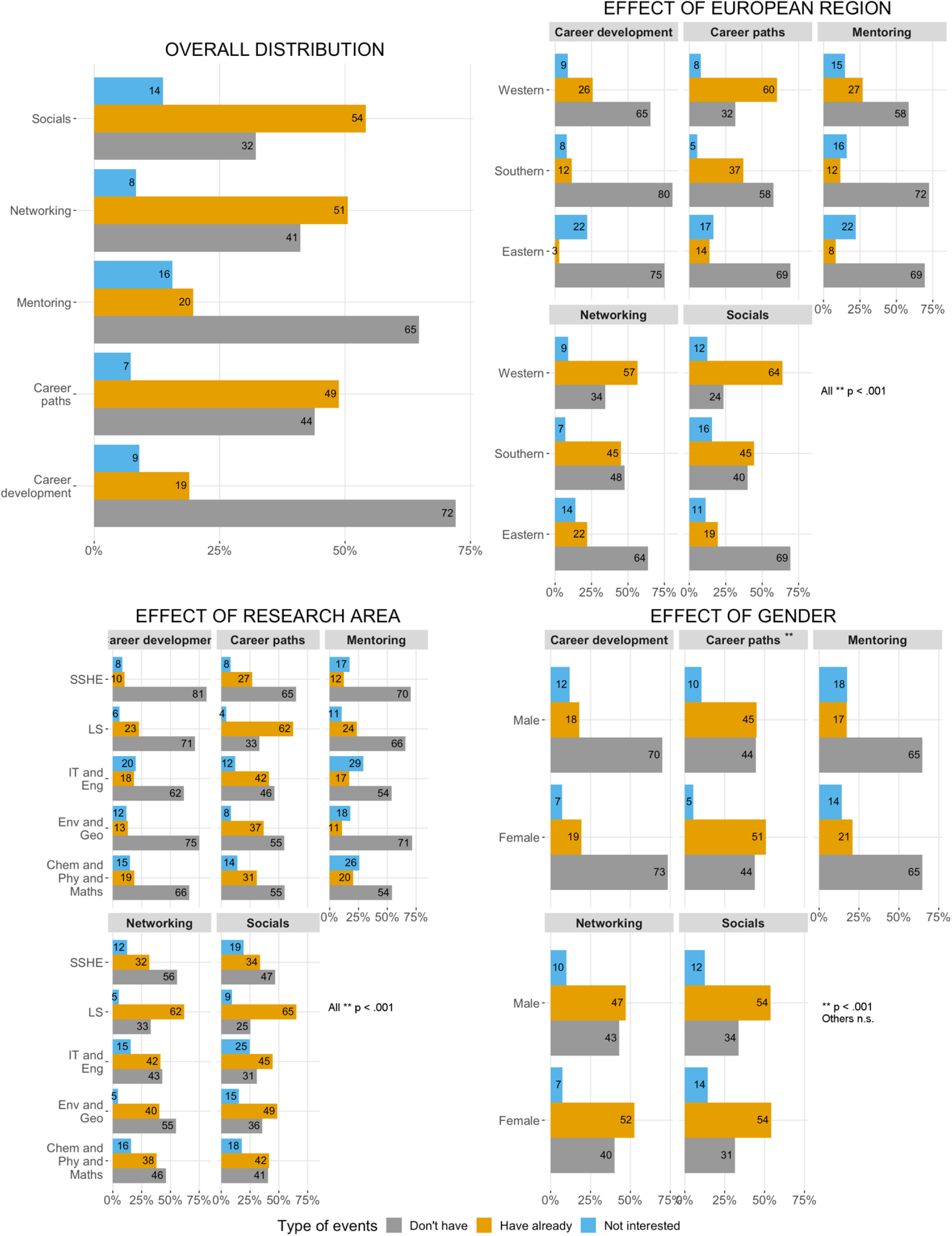
Type of events organised by postdoctoral associations and desired by postdoctoral researchers.

There were significant differences on the type of events postdoctoral associations already organise or are desired to do, depending on the region of work and area of research (*p* <.001). Postdoctoral researchers in Southern Europe would like postdoctoral associations to organise events related to career paths, whereas postdoctoral researchers in Western Europe already have access to these types of events (58% vs. 32%). Postdoctoral associations in Western Europe also organise more networking and social events than associations in Southern or Eastern Europe (57% and 64% vs 45% and 45% for Southern and 22% and 19% for Eastern Europe).

In general, postdoctoral researchers in the LifeSci felt that the postdoctoral associations were already organising events they were interested in, particularly for career paths, networking and social events. Career development and mentoring events seem to be lacking across all research areas, although postdoctoral researchers in IT&Eng and Chem&Phys&Maths showed less interest in the latter.

Finally, there was no effect of gender on the type of events postdoctoral researchers were interested in, except for career path events (*p* <.001). More female postdoctoral researchers reported that their postdoctoral associations already organise events on career paths, compared to male postdoctoral researchers (51% vs 45%, respectively), and male postdoctoral researchers showed less interest in this type of event (10% vs 5%, respectively).

## 4. Conclusions & Recommendations

This survey provided us with a comprehensive view of the demographics of postdoctoral researchers working in European institutions, and what their working environment is like. Based on our results, we are highlighting eight issues which might have a significant impact on the careers of postdoctoral researchers in Europe.

### Longer postdoctoral periods in Southern Europe despite higher publication metrics

Postdoctoral researchers working in Southern Europe reported higher number of publications and higher h-index than researchers in Western or Eastern Europe. This finding remained significant after controlling for the effect of time as a postdoctoral researcher and age. Therefore, the scientific output of postdoctoral researchers working in Southern Europe does not have lower impact than the output produced by postdoctoral researchers in the rest of Europe. Nevertheless, these researchers stay in postdoctoral positions for longer, suggesting difficulties in career progression despite their scientific achievements. Moreover, postdoctoral researchers from Southern European institutions report lower institutional support in career development which may account for such difficulties.

**Recommendation 1: Institutions in Southern Europe should develop clear criteria to support postdoctoral researchers’ career progression.**

### Southern and Eastern Europe pay the lowest salaries and have the lowest number of foreign postdoctoral researchers

Salary differences across European regions were remarkable: Southern regions pay around ⅔ of the Western salary and Eastern institutions pay a little over ⅓ of the Western salaries. Previous reports <u>(eurostat 2018)</u> suggest that even after controlling for cost of living, the differences persist, with some countries paying a lot more than others - Western vs (Southern & Eastern) <u>(European Commission 2017)</u>. Moreover, researchers from Western Europe rarely work as postdoctoral researchers in Southern or Eastern Europe. Differences in salaries might be a barrier for attracting researchers to these regions.

**Recommendation 2: The salary differences across European countries should be addressed as this could be a barrier to mobility and knowledge exchange from higher to lower pay regions.**

### Lack of access to funding is a significant concern of postdoctoral researchers

Around half of postdoctoral researchers were dissatisfied with access to funding, more so than with institutional support and quality of research facilities. Dissatisfaction is higher in Southern and Eastern Europe than in Western Europe, particularly concerning access to funding. This could be a barrier for scientific development and career progression in these regions.

**Recommendation 3: Discrepancies in access to funding should be minimized across Europe.**

### Postdoctoral researchers in Europe work longer hours than required by contract

It is well established that long hours at work are detrimental to our health <u>(Dinh, Strazdins, and Welsh 2017)</u>, yet only 34% of researchers on full-time contracts state that they work 40 hr/week or less. These numbers should raise some concern, given that 39% report that they work 50 hr/week or more and 11% report that they work 60 hr/week or more. Notably, around 80% of researchers say they work more than the hours defined in their work contract (8hr/week more on average, i.e. one more day a week). Around 60% of researchers with part-time contracts report that they are working full-time highlighting the possible abuse of part-time contracts for full time work.

**Recommendation 4: The culture of overwork in the research environment should be addressed in order to protect researchers against the risks associated with long hours at work.**

### The majority of full-time postdoctoral work contracts includes an exclusivity clause

Around 60% of postdoctoral contracts in Europe have exclusivity clauses. This percentage is particularly high in Southern Europe where it reaches 83% of the contracts. The inclusion of exclusivity clauses in these contracts might be counterproductive, as researchers at the end of their contracts will find themselves with no work experience outside academia, and will therefore be less attractive for prospective employers. Allowing researchers to work outside of their research duties, and be paid for it, would enhance their employability and facilitate the collaboration between academia and other job sectors (e.g. industry, clinic, teaching & education). In line with this idea, more than half of the postdoctoral researchers that are not involved in teaching, report they would like to be.

**Recommendation 5: Inclusion of exclusivity clauses in contracts for postdoctoral researchers should be optional in order to allow them to enhance their employability outside academia.**

### Postdoctoral researchers’ career development is poorly supported by their institutions

Most postdoctoral researchers do not have a clear career plan, and this is true regardless of European region or research area. There seems to be a lack of institutional support on career management, as only 38% of postdoctoral researchers reported that their institution has a postdoctoral support office, mainly in Western Europe. When such postdoctoral offices exist, they seem to play an important role, as respondents are mostly satisfied with the support they receive. Furthermore, the majority of researchers report a desire for further training in career management, the implementation of mentoring programmes and career development events.

**Recommendation 6: Postdoctoral researchers’ career prospects and career management should be much more supported by institutions in coordination with postdoctoral associations.**

### Lack of postdoctoral representation in governance is linked to unclear institutional duties and rights

Postdoctoral representation in institutional governance is still not an established practice across Europe, and more so in Eastern Europe. This contrasts with the crucial role that postdoctoral researchers have in the laboratory: doing the bulk of research, helping to supervise students and helping in grant writing. Moreover, only around 30% of postdoctoral researchers report that institutional duties and rights were clearly communicated to them at the start of their contract and this is correlated with postdoctoral representation in governance. Also, more than half of the researchers report the absence of clear rules for authorship and misconduct within their research groups. An example of the difficulties emerging from the lack of clear rules is the lack of formal recognition of a significant percentage of postdoctoral researchers’ contributions towards the supervision of students.

**Recommendation 7: Institutional governance bodies should include postdoctoral researcher representatives. This would ensure that the views of this vital staff group are heard, as well as making postdoctoral researchers feel more engaged with their own institutions. A flexible and proactive communication strategy at the institution and research group level should be developed, taking into account the sometimes transient nature of postdoctoral researchers’ posts.**

### Researchers show higher engagement with their local postdoctoral associations than with workers’ unions

Over half of postdoctoral researchers reported that their institution had a postdoctoral association, with the majority reporting that they were members of such an association. In contrast, only 22% were currently members of a workers union. These observations suggest that postdoctoral associations are currently an important mean to advocate for postdoctoral needs and wants, and that there is scope for more postdoctoral associations to be created and supported by the institutions, to better disseminate what their aims are, and how they can benefit postdoctoral researchers.

**Recommendation 8: Postdoctoral associations are an essential way to advocate for postdoctoral researchers at the governance level. Institutions should engage with, promote and support the work of postdoctoral associations.**

## 5. Methods

### Survey development and dissemination

The survey was developed by representatives of the European Network of Postdoctoral Associations (ENPA) members’ and collaborators, taking inspiration from previous surveys from the Karolinska Institute and Ecole polytechnique fédérale de Lausanne (EPFL) postdoc association surveys and <u>(Vitae 2014)</u>. Research areas descriptors were based on the Marie Sklodowska-Curie Actions classification (2015).

The survey questions can be found in the Supplementary Material and in the ENPA website. Data will be made available from the corresponding author, MJR, upon reasonable request.

The survey was **available online from June 2017 to January 2018** on the ENPA website (https://www.uc.pt/en/iii/postdoc/ENPA/Atividades). Each ENPA national association sent out invitation emails with a link to the survey to their members and to other European institutions. The survey was also spread through social networks to friends and colleagues.

### Number of respondents

The raw data was only accessible to the ENPA members involved in the design and analysis of the survey.

The survey was anonymous and no IP tracking was employed. In order to be able to identify duplicate entries, we requested that respondents insert a code (last three letters of their last name and year of birth). This information was uncoupled from the survey answers after elimination of duplicates. Three duplicate entries were found and removed. We also removed from the analysis the entries from researchers that were currently unemployed (N = 1), researchers working outside Europe (N = 35), researchers working in companies (N = 5), entries with inconsistencies regarding country of work and institution (N = 2), and 1 entry from a PhD student. This resulted in a total of 898 entries.

### Data analysis and visualization

Analyses were conducted collaboratively across different members of ENPA with Matlab R2017b, R 3.4.2 (https://www.r-project.org/), SPSS, and Excel. Data analysed in R was plotted using the ggplot2 package (https://cran.r-project.org/web/packages/ggplot2/index.html), and using the colour palettes from http://www.cookbook-r.com/Graphs/Colors_(ggplot2)/#a-colorblind-friendly-palette.

For most questions on the survey, participants could choose to answer ‘*Prefer not to answer*’. Only a small number of respondents chose not to answer in each question. For the analysis, this option was considered as missing data. This resulted in varying response rates per item.

Descriptive statistics were used for data presentation (absolute and relative frequencies for categorical data; median and interquartile range for ordinal and continuous data). Statistical comparisons were performed to compare across researchers working in different European regions (Western Europe, Southern Europe and Eastern Europe), to compare across gender, to compare across research areas and to compare researchers working in their home country with researchers working abroad. Other comparisons specific to each survey question are described in the results section. To compare across European regions, we grouped the countries where researchers worked as described in Annex 2. To compare across research areas, we grouped them as described in Annex 4.

Statistical analyses of categorical variables were conducted with chi-square tests. Statistical comparisons of numerical variables were conducted with Kruskal-Wallis test (assuming non-normality of the data). Significant was defined as *p* <.05.

All percentage points were rounded to the nearest integer, whenever increased precision was not required. Numerical data were represented graphically as boxplots with Matlab boxplot function. On each box, the central mark is the median, the edges of the box are the 25th and 75th percentiles, and the whiskers extend to the most extreme data points that are not outliers. For ease of visualization, outliers are not plotted in the graphs.

For the questions listed below, we detail specific data treatment and analysis.

Q16: We excluded all the entries where the length of the postdoctoral period reported was longer than the time between survey submission and PhD conclusion. This process excluded 17 entries. Q19: Excluded from analysis all the free text entries from the ‘other’ option. This resulted in an N = 817.

Q20-22: Excluded from analysis respondents that selected full-time contract but said the number of hours contracted was less than 25/week; respondents that selected part-time contract and stated that number of hours contracted was more than 35/week; and all respondents that stated that the number of hours contracted were more than 50/week. These criteria excluded 11 entries. N = 887.

Q23-24: Only respondents that reported to be contracted on full-time contracts were included in the analyses of researchers’ salaries. Moreover, we excluded respondents that selected full-time contract but said the number of hours contracted was less than 25/week and respondents that stated that the number of hours contracted were more than 50/week. We also excluded researchers that reported annual gross salaries below 4000 euros and above 200000 euros, and entries where the net salary was higher than the gross salary. After applying these exclusion criteria we were left with N = 617 entries.

Q37-38: For the analysis of the h-index data, we only included the entries where respondents reported that they calculated their h-index using Scopus, Researcher ID or Google Scholar. We excluded all other entries. Furthermore, this question was not compulsory and there were a number of missing responses. This resulted in an N = 565 for this analyses.

Q43-45: Responses aggregated as ‘Dissatisfied’ (‘Very dissatisfied’ and ‘Dissatisfied’) and ‘Satisfied’ (‘Very Satisfied’ and ‘Satisfied’). Using only employers for which there is more than 1% of the total responses (13 institutions), in each (Facilities, Institutional and Funding). This resulted in an N = 562 for these analyses. Statistical significance was also calculated using the aggregated responses.

Q46: An N *=* 892 was used. Statistical significance was calculated using aggregated responses as ‘Defined’ (‘Clearly defined rules’ and ‘Partially defined rules’) and ‘Not defined’ (‘I don’t know’ and ‘Undefined’).

Q47: An N *=* 892 was used.

Q55: An N *=* 892 was used. Statistical significance was calculated using only ‘Yes’ and ‘No’ answers (N *=* 871).

Q57: An N *=* 465 was used. Statistical significance for European region did not include Eastern Europe, as there were only 3 responses.

Q58: An N = 476 was used. Free text answers to the survey were categorised and aggregated to existing categories when possible. Statistical significance for European region was calculated only for ‘Institution’, ‘Fellow postdoctoral researchers’, ‘Induction’ and ‘Internet Search’ as the other categories had very few responses. Only ‘Southern’ and ‘Western Europe’ were used, as there were very few responses from ‘Eastern Europe’. Statistical significance for research area was not calculated because of the low number of responses in each. On the effect of gender, only for ‘Institution’, ‘Fellow postdoctoral researchers’, ‘Induction’ and ‘Internet Search’ were used, due to low number of responses in the other categories.

Q59: An N = 898 was used. Statistical significance for European region was calculated after removing responses for ‘Eastern Europe’ as there were very few, except for the ‘Mentoring’ category.

## Supporting information

European survey for postdoctoral researchers

## 7. Authors

The postdoctoral associations and researchers involved in the development and analyses of the survey and report writing are listed below.

### Postdocs@UC, University of Coimbra, Portugal

**Maria Ribeiro** Postdoctoral researcher in Cognitive Neuroscience at the University of Coimbra, Portugal. Coordinator of the Projects Committee of the Initiative Postdocs@UC and of the European Network of Postdoctoral Associations (ENPA) aiming to promote discussion regarding the development of solutions improving postdoctoral researchers work conditions and career development.

Contributions: survey design and analyses, including statistical analyses, graphical representation of data, results discussion and report writing **Ana Fonseca** She holds a PhD in Clinical Psychology since 2013 and she is currently a postdoctoral fellow at Center for Research in Neuropsychology and Cognitive-Behavioral Intervention (University of Coimbra).

Contributions: survey design and analyses, including statistical analyses, graphical representation of data, results discussion and report writing.

**Mariana Moura Ramos** She completed her PhD in Psychology in the University of Coimbra in 2011. She works as a clinician at the Coimbra Centre University Hospital and as a researcher at the University of Coimbra. She makes strong efforts to translate the research findings into the clinical practice.

Contributions: survey design and analyses, including statistical analyses, graphical representation of data, results discussion and report writing.

### The Postdocs of Cambridge Society (PdOC), United Kingdom

**Marta Costa** completed her PhD at the University of Cambridge, UK and is currently a Senior Research Associate at the Department of Zoology. She has been a member of The Postdocs of Cambridge Society for the last 3 years, focusing on Researcher Development.

Contributions: survey design and analyses, including report writing, statistical analyses, results discussion and graphical representation of data.

### Karolinska Institutet Postdoc Association (KIPA), Sweden

**Konstantina Kilteni** She studied Electrical and Computer Engineering at the National Technical University of Athens and she holds a PhD in Virtual Reality and Psychology from the University of Barcelona. She is currently a Marie Sklodowska-Curie postdoctoral fellow at the Department of Neuroscience in Karolinska Institutet.

Contributions: survey analyses, including statistical analyses, graphical representation of data, results discussion and report writing.

**Lau Møller Andersen** He completed his PhD in Clinical Medicine in the University of Aarhus 2016. He works as a postdoc at NatMEG at Karolinska Institutet, where he is specialising in new technologies within magnetoencephalography and pushing open science within the field.

Contributions: survey analyses, including statistical analyses, graphical representation of data, results discussion and report writing.

**Lisa Harber-Aschan** is a postdoctoral researcher at the Department of Public Health Sciences at Karolinska Institutet, Sweden. She completed her PhD in Epidemiological Psychiatry at the Institute of Psychiatry, Psychology and Neuroscience, King’s College London, United Kingdom. Contributions: planning statistical analyses and presentation and interpretation of results.

### i3S Postdoctoral Association, University of Porto, Portugal

**Joana Moscoso**

Marie-Curie Postdoctoral Fellow and Director of Native Scientist. She holds a PhD from Imperial College London in Microbiology and she received the 2017 MIT Innovator Under 35 recognition for her humanitarian work. She dreams of having her own restaurant one day.

Contributions: discussion of survey results and report writing.

### Uppsala University Postdoc Association (UUPA), Sweden

**Sonchita Bagchi** Researcher funded by external grants from Swedish agencies and foundations. Originally coming from India, she holds a PhD from Uppsala University in Microbiology. She has founded the postdoctoral association at Uppsala University in 2017 and is actively working to make a difference in lives of postdoctoral researchers in Sweden.

Contributions: discussion of survey results and report writing.

In addition, we would like to acknowledge the contribution of the following researchers and their postdoctoral associations in the design and dissemination of the survey:

Postdocs@UC, University of Coimbra, Portugal

Diogo Proença, Nuno Mendonça, Chiara Carrozza, Antonieta Reis Leite, Marta Passadouro

Karolinska Institutet Postdoc Association (KIPA), Sweden Elisa Floriddia, Alessandro Bosco and João Rosa

The Postdocs of Cambridge Society (PdOC), United Kingdom Adina Feldman

The Postdoc Association (PDA), IST, Austria

Marion Picard, Giacomo Bighin, Laura Rodriguez and Ben Suter

MPI Postdoctoral Community, Max Planck Institute for Terrestrial Microbiology, Germany Anja Paulick, Alvaro Orell and Deepak Anand

Postdoc Initiative at the MPIPZ (PIM), Max Planck Institute for Plant Breeding Research, Germany Clementine Leroux

Postdocs of the MPI-CBG in Dresden, Max Planck Institute of Molecular Cell Biology and Genetics, Germany

Ju Roscito, Karina Pombo-Garcia, Rana Amini, and Lisa Dennison

## 6. Annexes

**Annex 1.** Institutions represented in the survey

| Employer | Responses |
| --- | --- |
| University of Coimbra | 145 |
| Karolinska Institute | 132 |
| Imperial College London | 77 |
| University of Cambridge | 38 |
| University of Porto | 37 |
| University of Lisbon | 30 |
| Nova University of Lisbon | 23 |
| Centre for Genomic Regulation | 19 |
| Max Planck Institute for Molecular Cell Biology and Genetics | 16 |
| Uppsala University | 15 |
| King's College London | 11 |
| Instituto de Salud Global de Barcelona | 10 |
| Consejo Superior de Investigaciones Científicas | 9 |
| University of Córdoba | 8 |
| Czech Academy of Sciences | 6 |
| University College London | 6 |
| University of Copenhagen | 6 |
| University College Dublin | 5 |
| University of Milan | 5 |
| University of Minho | 5 |
| Aarhus University | 4 |
| Max Planck Institute | 4 |
| Pompeu Fabra University | 4 |
| University of Milan Bicocca | 4 |
| University of Turin | 4 |
| Centre National de la Recherche Scientifique | 3 |
| Gheorghe Asachi Technical University of Iasi | 3 |
| Hungarian Academy of Sciences | 3 |
| ISPA | 3 |
| KTH Royal Institute of Technology | 3 |
| Leiden University | 3 |
| Lund University | 3 |
| MRC Laboratory of Molecular Biology | 3 |
| Slovak Academy of Sciences | 3 |
| Umeå University | 3 |
| University of Bergen | 3 |
| University of Oxford | 3 |
| Aalto University | 2 |
| ETH Zurich | 2 |
| European Molecular Biology Laboratory | 2 |
| Ghent University | 2 |
| iBET-Institute of Experimental and Technological Biology | 2 |
| Institut d'Investigació Germans Trias i Pujol | 2 |
| Italian National Research Council | 2 |
| KU Leuven | 2 |
| Ludwig Maximilian University of Munich | 2 |
| Max Delbrück Center for Molecular Medicine | 2 |
| Queen Mary University of London | 2 |
| Radboud University | 2 |
| Stockholm University | 2 |
| Technical University of Catalonia | 2 |
| Technical University of Munich | 2 |
| The Wellcome Trust Sanger Institute | 2 |
| Universidad Autónoma de Madrid | 2 |
| Universitat Autònoma de Barcelona | 2 |
| Universitat de Barcelona | 2 |
| University College Cork | 2 |
| University of Algarve | 2 |
| University of Aveiro | 2 |
| University of Bologna | 2 |
| University of Bristol | 2 |
| University of Geneva | 2 |
| University of Hull | 2 |
| University of Lausanne | 2 |
| University of Liverpool | 2 |
| University of London | 2 |
| University of Oslo | 2 |
| University of Plymouth | 2 |
| University of Rome Tor Vergata | 2 |
| University of Southampton | 2 |
| University of Split | 2 |
| University of Urbino | 2 |
| "Ion Mincu" University of Architecture and Urbanism | 1 |
| Aix-Marseille University | 1 |
| Aristotle University of Thessaloniki | 1 |
| Banat's University of Agricultural Sciences and Veterinary Medicine from Timisoara | 1 |
| Bangor University | 1 |
| Bern University | 1 |
| Botanischer Garten Botanisches Museum Berlin | 1 |
| Brighton and Sussex Medical School | 1 |
| Catholic University of Louvain (Louvain-la-Neuve) | 1 |
| CEA | 1 |
| Center for Regenerative Therapies Dresden | 1 |
| Centre National D'Etudes Spatiales | 1 |
| Charite | 1 |
| Charles University Prague | 1 |
| Christian-Albrechts-Universität zu Kiel | 1 |
| CHUV | 1 |
| Clausthal University of Technology | 1 |
| Czech Technical University in Prague | 1 |
| DKFZ | 1 |
| Durham University | 1 |
| ELI Beamlines | 1 |
| ENS | 1 |
| EPFL | 1 |
| Erasmus Medical Centre | 1 |
| Essen University hospital | 1 |
| European Synchrotron Radiation Facility | 1 |
| Forschungszentrum Jülich | 1 |
| Free University Amsterdam | 1 |
| Friedrich-Alexander University of Erlangen Nuremberg | 1 |
| Friedrich-Schiller-Universität Jena | 1 |
| Gregor Mendel Institute | 1 |
| Gulbenkian Science Institute | 1 |
| Heinrich-Heine University Düsseldorf | 1 |
| IFREMER | 1 |
| INSERM | 1 |
| Institut d'Investigacions Biomèdiques August Pi i Sunyer | 1 |
| Institut du Fer à Moulin | 1 |
| Institut FEMTO-ST | 1 |
| Institute for Bioengineering of Catalonia | 1 |
| Institute for Bioprocessing and Analytical Measurement Techniques | 1 |
| Institute of Archaeology and Ethnology Polish Academy of Science | 1 |
| Institute of Astrophysics and Space Science | 1 |
| Institute of High Energy Physics | 1 |
| Institute of Organic Chemistry, Pan, Warsaw | 1 |
| Instituto do Mar (MARE) | 1 |
| Instituto Nacional de Investigación y Tecnología Agraria y Alimentaria | 1 |
| International Agency for Research on Cancer | 1 |
| International School of Advanced Studies | 1 |
| IRCCS Ospedale San Raffaele | 1 |
| Jagiellonian University in Kraków | 1 |
| John Innes Centre | 1 |
| Lancaster University | 1 |
| London School of Economics and Political Sciences | 1 |
| Loughborough University | 1 |
| Ludwig Institute for Cancer Research | 1 |
| Luleå University of Technology | 1 |
| Maastricht University | 1 |
| Marche Polytechnic University | 1 |
| Martin Luther University of Halle-Wittenberg | 1 |
| Max Planck Institute for Biophysics | 1 |
| Max Planck Institute for Molecular Genetics | 1 |
| Max Planck Institute of Neurobiology | 1 |
| Medical Research Council | 1 |
| Medical University of Graz | 1 |
| Mediterranea University of Reggio Calabria | 1 |
| Middle East Technical University | 1 |
| MRC London Institute of Medical Sciences | 1 |
| Museum für Naturkunde | 1 |
| National Center for Scientific Research Demokritos | 1 |
| National University of Ireland in Galway | 1 |
| Natural History Museum London | 1 |
| New York University | 1 |
| Newcastle University | 1 |
| Nicolaus Copernicus University in Torun | 1 |
| Nordita (KTH & Stockholm University) | 1 |
| Office de la Culture, Canton Jura | 1 |
| Örebro University | 1 |
| Pau Costa Foundation | 1 |
| Polish Academy of Science | 1 |
| Research Institute for Hungarian Communities Abroad | 1 |
| Royal College of Music | 1 |
| Royal College of Surgeons | 1 |
| RWTH Aachen University | 1 |
| Scuola Normale Superiore | 1 |
| SHOM | 1 |
| Technical University of Denmark | 1 |
| Technical University of Dortmund | 1 |
| Technische Universität Dresden | 1 |
| Telecommunications Institute | 1 |
| Telethon Institute of Genetics and Medicine | 1 |
| Tu Delft | 1 |
| Unige | 1 |
| Universidad Complutense Madrid | 1 |
| Universidad de La Laguna | 1 |
| Universidad Pablo de Olavide (CABD) | 1 |
| Universidade de Santiago de Compostela | 1 |
| Universität Bonn | 1 |
| Universität Bremen | 1 |
| Universität Duisburg-Essen | 1 |
| Universität Göttingen | 1 |
| Universität Hamburg | 1 |
| Universität Siegen | 1 |
| Universität Tübingen | 1 |
| Universität Zu Köln | 1 |
| Université Libre de Bruxelles | 1 |
| Universite Paul Sabatier | 1 |
| University Brussels | 1 |
| University Claude Bernard Lyon 1 | 1 |
| University of Alcalá | 1 |
| University of Bergamo | 1 |
| University of Bialystok | 1 |
| University of Bordeaux | 1 |
| University of Bucharest | 1 |
| University of Burgundy | 1 |
| University of Cagliari | 1 |
| University of Calabria | 1 |
| University of Cardiff | 1 |
| University of Clermont Auvergne | 1 |
| University of Cyprus | 1 |
| University of Deusto | 1 |
| University of Dundee | 1 |
| University of East Anglia | 1 |
| University of Eastern Finland | 1 |
| University of Eastern Piedmont | 1 |
| University of Ferrara | 1 |
| University of Franche-Comte | 1 |
| University of Girona | 1 |
| University of Granada | 1 |
| University of Graz | 1 |
| University of Helsinki | 1 |
| University of Jyväskylä | 1 |
| University of Kent | 1 |
| University of Konstanz | 1 |
| University of Leuven | 1 |
| University of Lincoln | 1 |
| University of Louvain la Neuve | 1 |
| University of Luxembourg | 1 |
| University of Madeira | 1 |
| University of Murcia | 1 |
| University of Naples Frederico II | 1 |
| University of Natural Resources and Life Science | 1 |
| University of Navarre | 1 |
| University of Padova | 1 |
| University of Pavia | 1 |
| University of Portsmouth | 1 |
| University of Pula | 1 |
| University of Southern Denmark | 1 |
| University of St Andrews | 1 |
| University of Stirling | 1 |
| University of Strasbourg | 1 |
| University of Surrey | 1 |
| University of Sussex | 1 |
| University of Tartu | 1 |
| University of the Azores | 1 |
| University of the Basque Country | 1 |
| University of Trento | 1 |
| University of Trieste | 1 |
| University of Twente | 1 |
| University of Verona | 1 |
| University of Vigo | 1 |
| University of Warwick | 1 |
| University of Wroclaw | 1 |
| University of York | 1 |
| University of Zadar | 1 |
| University of Zurich | 1 |
| University Paris Descartes | 1 |
| University Tras os Montes e Alto Douro | 1 |
| UPMC | 1 |
| Vilnius University | 1 |
| Total: 241 | 898 |

**Annex 2.**
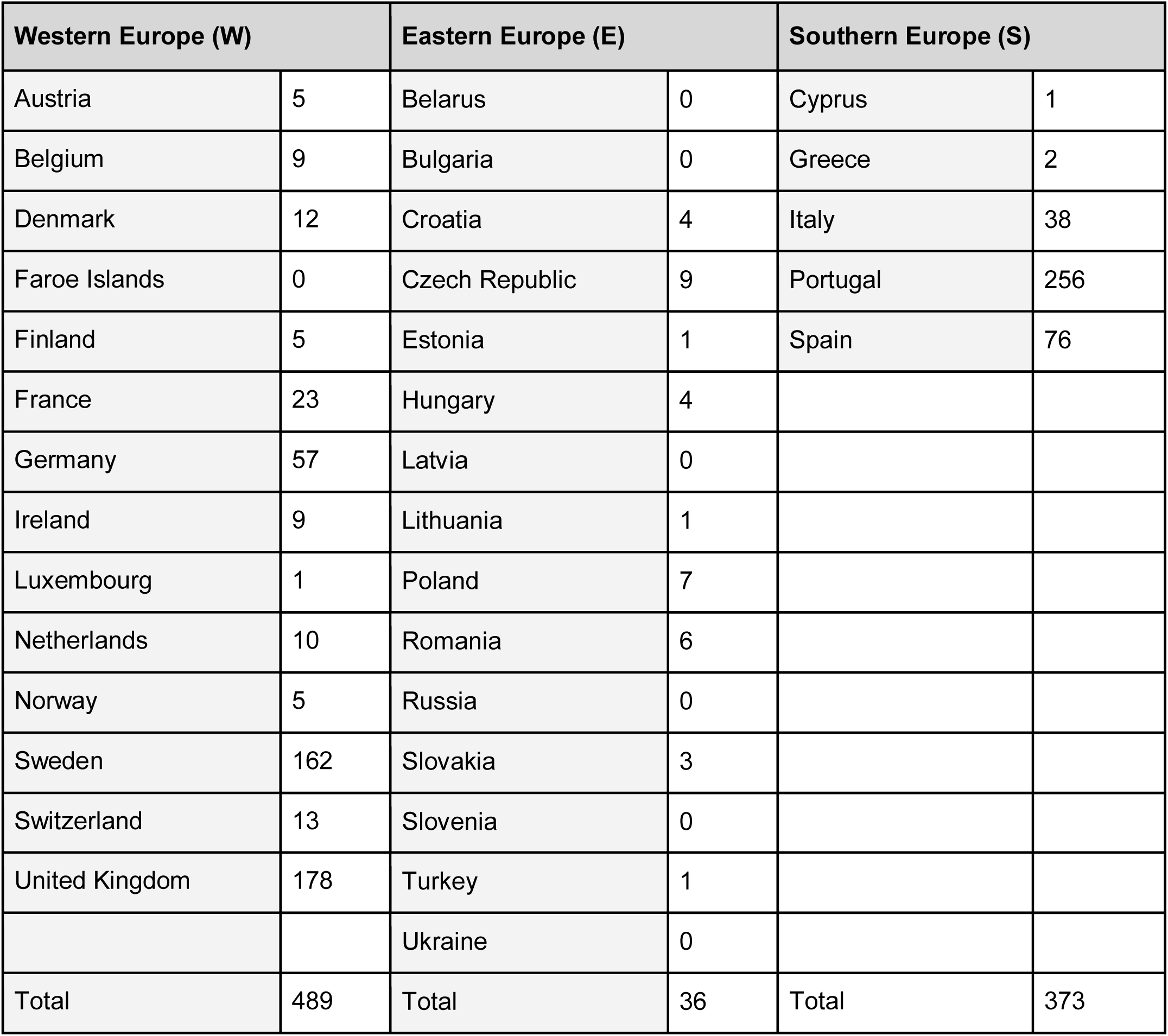
Responses per country of work and groupings

**Annex 3.**
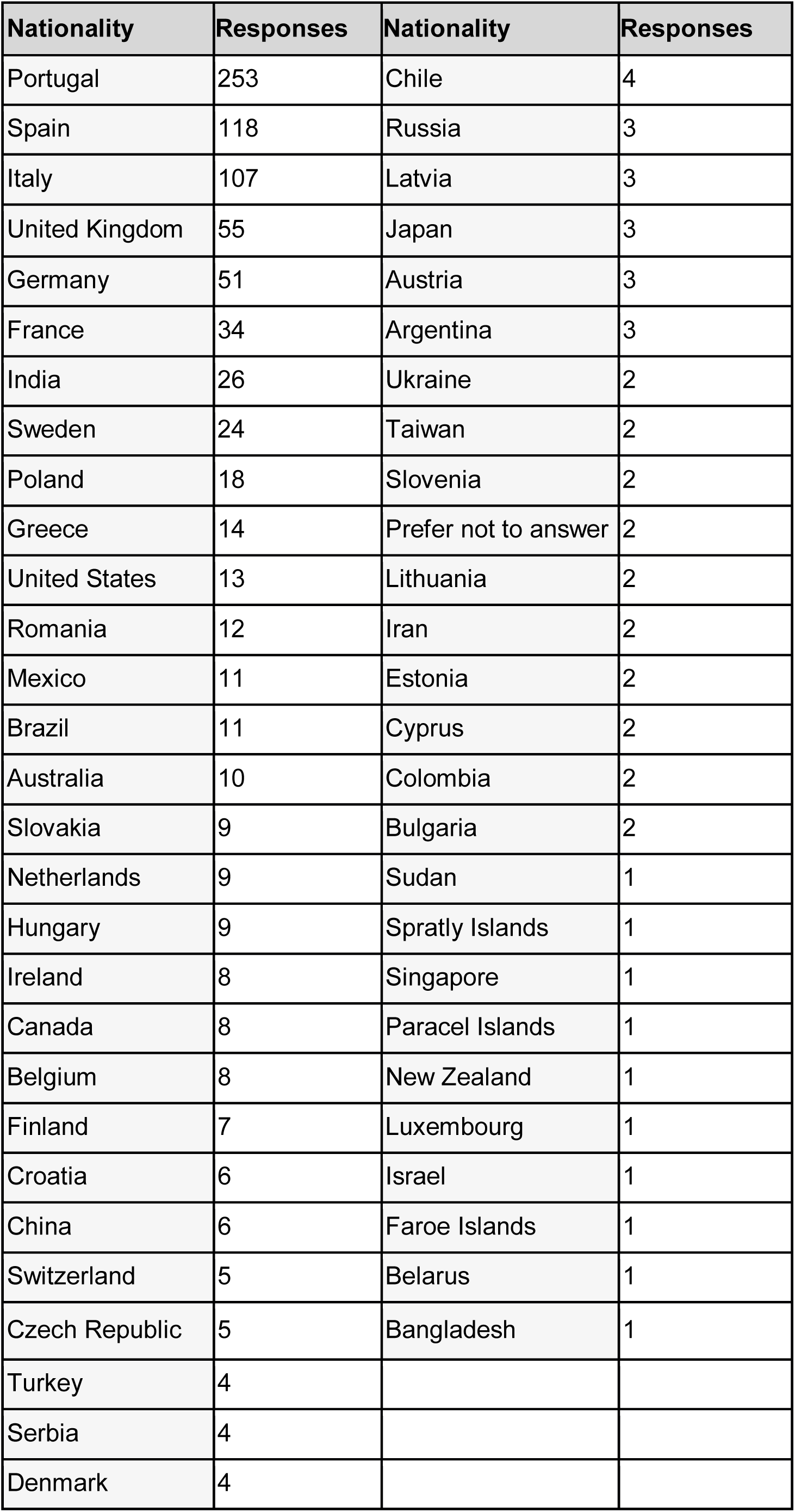
Responses per nationality

**Annex 4.**
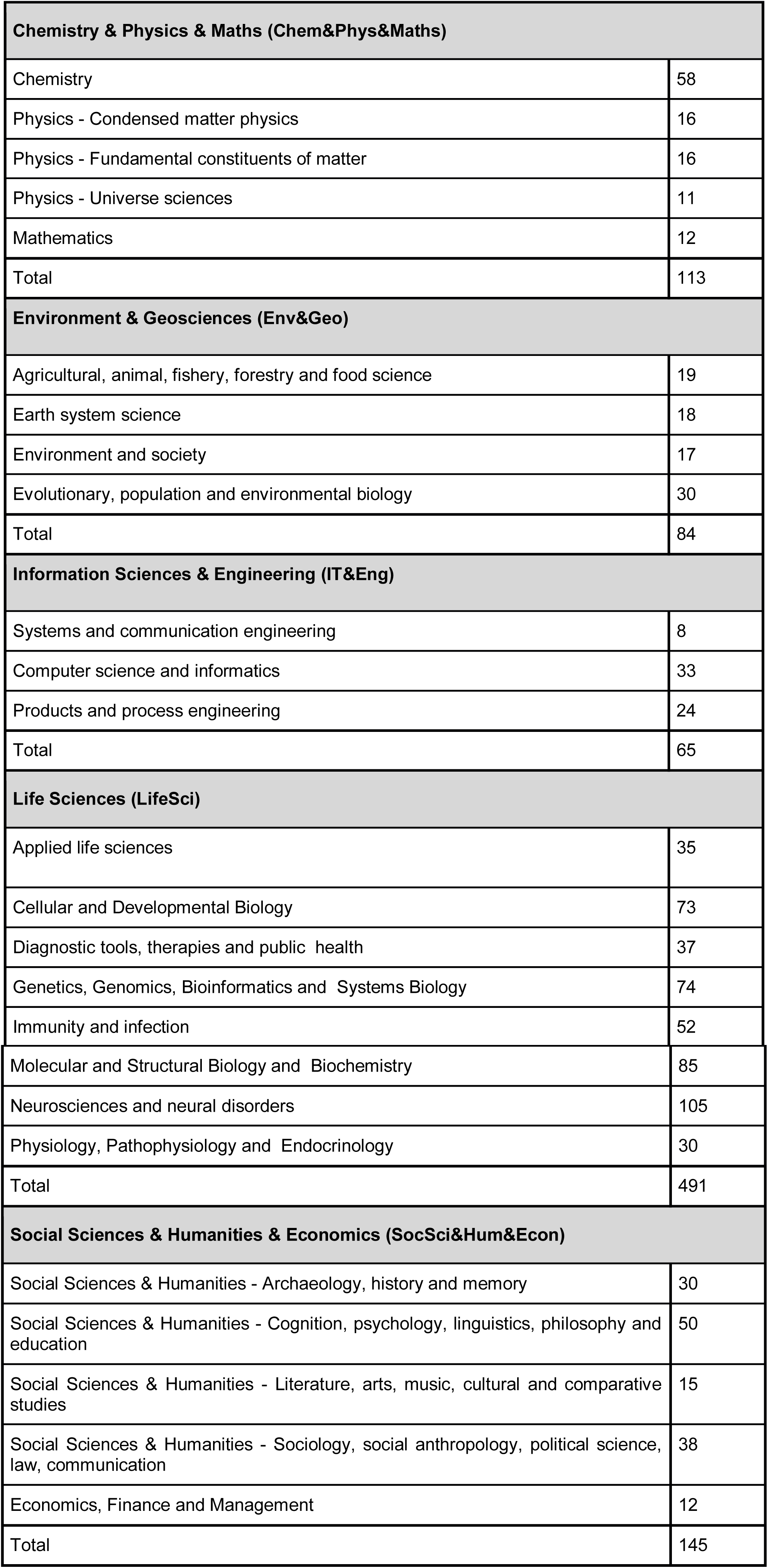
Responses per Primary Research Area and groupings

**Annex 5.** Responses per gender

| Gender | Responses |
| --- | --- |
| Female | 547 |
| Male | 344 |
| Prefer not to answer | 7 |

### Annex 6 Analysis of income differences within Europe

#### Sample

We started with an initial number of records: 898 people. We chose a significance level of α = 0.05.

#### Exclusion criteria

1. Respondents that did not report which country they work in. Removed 0. Rest 898.
2. Respondents that did not report nationality. Removed 0. Rest 898.
3. Respondents that did not identify with the male/female category. Removed 7. Rest 891.
4. Respondents that did not report gross income. Removed 101. Rest 79.
5. Respondents that reported gross income < 1000 euros per year and >100.000 euros per year. Removed 23. Rest 767.
6. Respondents that did not work full time (including NAN and part-time). Removed 64. Rest 703.
7. Respondents that did not report months since PhD. Removed 0. Rest 703.

#### Further processing

- We grouped nationalities and work countries to Western Europe (incl Australia, US, N Zealand), Eastern Europe, Southern Europe, South&Central America, Asia, Africa as specified in Annex 2.
- Asians, Eastern Europeans, South&Central Americans. Removed (1) working on Asia, everyone working in S&C America (n=0), everyone working in Africa (n=0). Rest 702.
- Removed people with Asian nationality (n=34), from S&C America (22), from Africa (0). Rest 646.

#### Predictor variables

- Gender
- Age
- Children
- Years since completing PhD
- Country of origin
- Country of working
- Research area
- Mobility

#### Descriptive tables

We were interested in describing the income distribution for postdoctoral researchers across Europe. We were especially interested in whether there were gender differences in income. From our dataset, we identified *Gender, Age, Children, Years since completing PhD, Country of Origin, Country of Working, Research area* and *Mobility* as the independent variables of interest. For the purposes of initial descriptive analysis, we described mean *gross income in euros* by all independent variables in the total sample, and stratified by gender.

**Table 1:**
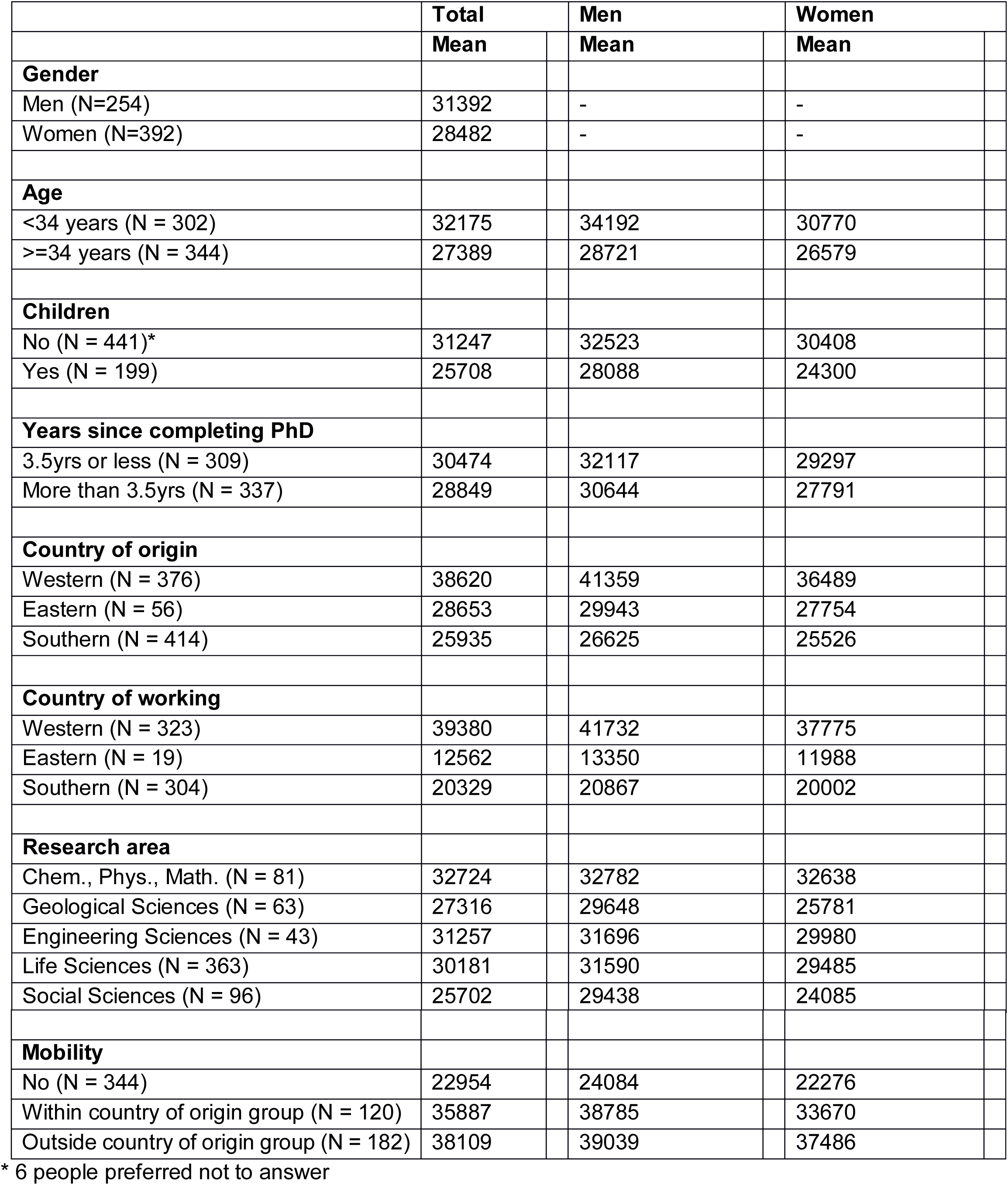
Mean gross income by all independent variables, and by gender (N=646)

|  | Total | Men | Women |
| --- | --- | --- | --- |
|  | Mean | Mean | Mean |
| **Gender** |  |  |  |
| Men (N=254) | 31392 | - | - |
| Women (N=392) | 28482 | - | - |
| **Age** |  |  |  |
| <34 years (N = 302) | 32175 | 34192 | 30770 |
| >=34 years (N = 344) | 27389 | 28721 | 26579 |
| **Children** |  |  |  |
| No (N = 441)* | 31247 | 32523 | 30408 |
| Yes (N = 199) | 25708 | 28088 | 24300 |
| **Years since completing PhD** |  |  |  |
| 3.5yrs or less (N = 309) | 30474 | 32117 | 29297 |
| More than 3.5yrs (N = 337) | 28849 | 30644 | 27791 |
| **Country of origin** |  |  |  |
| Western (N = 376) | 38620 | 41359 | 36489 |
| Eastern (N = 56) | 28653 | 29943 | 27754 |
| Southern (N = 414) | 25935 | 26625 | 25526 |
| **Country of working** |  |  |  |
| Western (N = 323) | 39380 | 41732 | 37775 |
| Eastern (N = 19) | 12562 | 13350 | 11988 |
| Southern (N = 304) | 20329 | 20867 | 20002 |
| **Research area** |  |  |  |
| Chem., Phys., Math. (N = 81) | 32724 | 32782 | 32638 |
| Geological Sciences (N = 63) | 27316 | 29648 | 25781 |
| Engineering Sciences (N = 43) | 31257 | 31696 | 29980 |
| Life Sciences (N = 363) | 30181 | 31590 | 29485 |
| Social Sciences (N = 96) | 25702 | 29438 | 24085 |
| **Mobility** |  |  |  |
| No (N = 344) | 22954 | 24084 | 22276 |
| Within country of origin group (N = 120) | 35887 | 38785 | 33670 |
| Outside country of origin group (N = 182) | 38109 | 39039 | 37486 |

\* 6 people preferred not to answer

Table 1 illustrates that in our sample:

- Men overall earned ∼3000 € more than women
- Young postdocs (< 34 y) earned more than old postdocs (>= 34 y), ∼5000 €
- Postdocs without children earned more than those with children, ∼5500 €
- Postdocs in the West earned more than in the East, ∼27000 €, and in the South 19000 €
- Mobile postdocs earned more than non-mobile postdocs, ∼13000-15000 €
- Social scientists earned less compared to other subject areas

#### Testing adjusted and unadjusted models

Linear regression models then tested these associations. Table 2 presents unadjusted and adjusted estimates with 95% confidence intervals and *p*-values. Unadjusted estimates are based on models including only the variable of interest. The adjusted analysis includes all variables presented in the table. Thus the adjusted estimates control for the effect of the other variables

**Table 2:** Unadjusted and adjusted associations between gender and other independent variables of interest and gross annual income in euros (N=646)

|  | Unadjusted |  |  | Adjusted |  |  |
| --- | --- | --- | --- | --- | --- | --- |
|  | Coef. | (95% CI) | *p* | Coef. | (95% CI) | *p* |
| **Gender** |  |  |  |  |  |  |
| Women | ref. |  |  |  |  |  |
| Men | 2910 | (517, 5303) | 0.017 | 2192 | (318, 4066) | 0.022 |
| **Months since PhD** | -7 | (-37, 23) | 0.645 | 59 | (31, 87) | <0.001 |
| **Children** |  |  |  |  |  |  |
| No | ref. |  |  |  |  |  |
| Yes | -5539 | (-8047, -3031) | <0.001 | -645 | (-2795, 1504) | 0.56 |
| **Age in years** | -645 | (-859, 431) | <0.001 | -394 | (-618, -170) | <0.001 |
| **Country of origin** |  |  |  |  |  |  |
| Western | ref. |  |  |  |  |  |
| Eastern | -9967 | (-14231, -5703) | <0.001 | -7806 | (-12374, -3240) | <0.001 |
| Southern | -12685 | (-15186, -10184) | <0.001 | -4602 | (-7882, -1321) | 0.0060 |
| **Country of working** |  |  |  |  |  |  |
| Western | ref. |  |  |  |  |  |
| Eastern | -26818 | (-32185, -21451) | <0.001 | -20289 | (-26580, -13998) | <0.001 |
| Southern | -19051 | (-20867, -17234) | <0.001 | -14163 | (-17407, -10918) | <0.001 |
| **Research area** |  |  |  |  |  |  |
| Chemistry, Physics, Mathematics | 2543 | (-1098, 6184) | 0.17 | 1187 | (-1613, 3987) | 0.41 |
| Geological Sciences | -2865 | (-6908, 1179) | 0.17 | 900 | (-2169, 3968) | 0.56 |
| Engineering Sciences | 1076 | (-3702, 5855) | 0.66 | 809 | (-2892, 4510) | 0.67 |
| Life Sciences | ref. |  |  |  |  |  |
| Social Sciences | -4478 | (-7878, -1078) | 0.010 | 1537 | (-1143, 4217) | 0.26 |
| **Mobility from birth country** |  |  |  |  |  |  |
| No | ref. |  |  |  |  |  |
| Within country of origin group | 12932 | (10141, 15724) | <0.001 | 470 | (-2453, 3392) | 0.75 |
| Outside country of origin group | 15155 | (12741, 17568) | <0.001 | 5423 | (2195, 8651) | 0.0010 |

We here found that even in the adjusted model, accounting for possible confounders, there was a main effect of gender with men earning 2192 € more annually than women. Other effects were:

- Longer time since completed PhD, was associated with greater earnings
- Older age was associated with less earnings
- Postdoctoral researchers from Southern and Eastern areas of Europe earned less than postdoctoral researchers from Western areas
- Postdoctoral researchers working in the Southern and Eastern regions of Europe earned less that postdoctoral researchers working in Western regions
- Postdoctoral researchers working outside their region of nationality earned more than postdoctoral researchers staying in their region of nationality and postdoctoral researchers who stayed in their home country
- There were no effects observed for having children and subject areas

#### Interactions

We tested four specific gender interaction, *children, work country, research area* and *mobility,* as these are likely to differentially impact on earning opportunities for men and women. This analysis is adjusted for all other covariates.

**Table 3:**
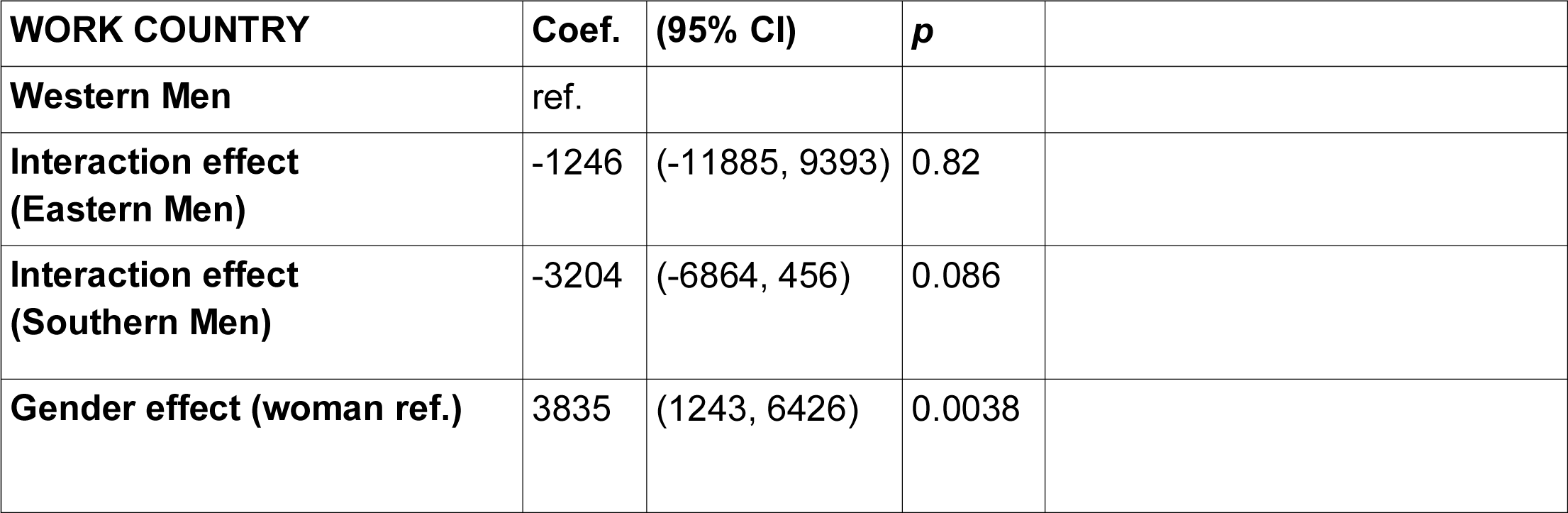
Interaction between Gender and Children (gross income)

No significant interaction effect was found between Gender and Children.

**Table 4:**
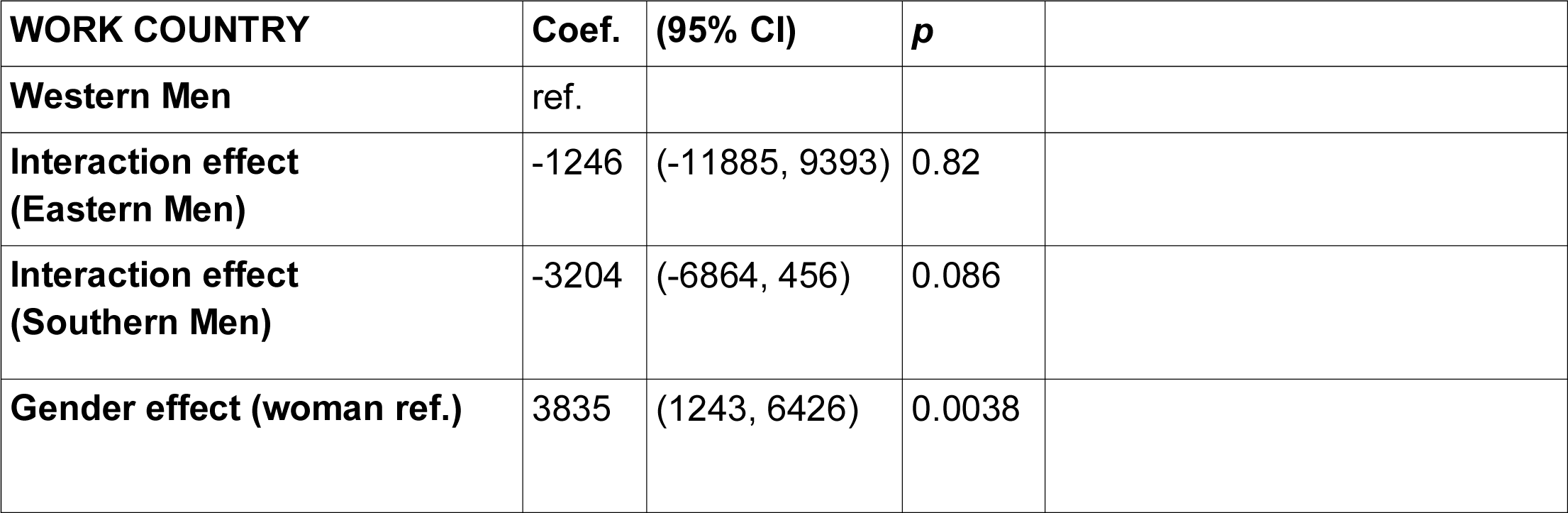
Interaction between Gender and Work Country (gross income)

No significant interaction effect was found between Gender and Work Country, though some indication that gender differences might be smaller in the South than in the East.

**Table 5:**
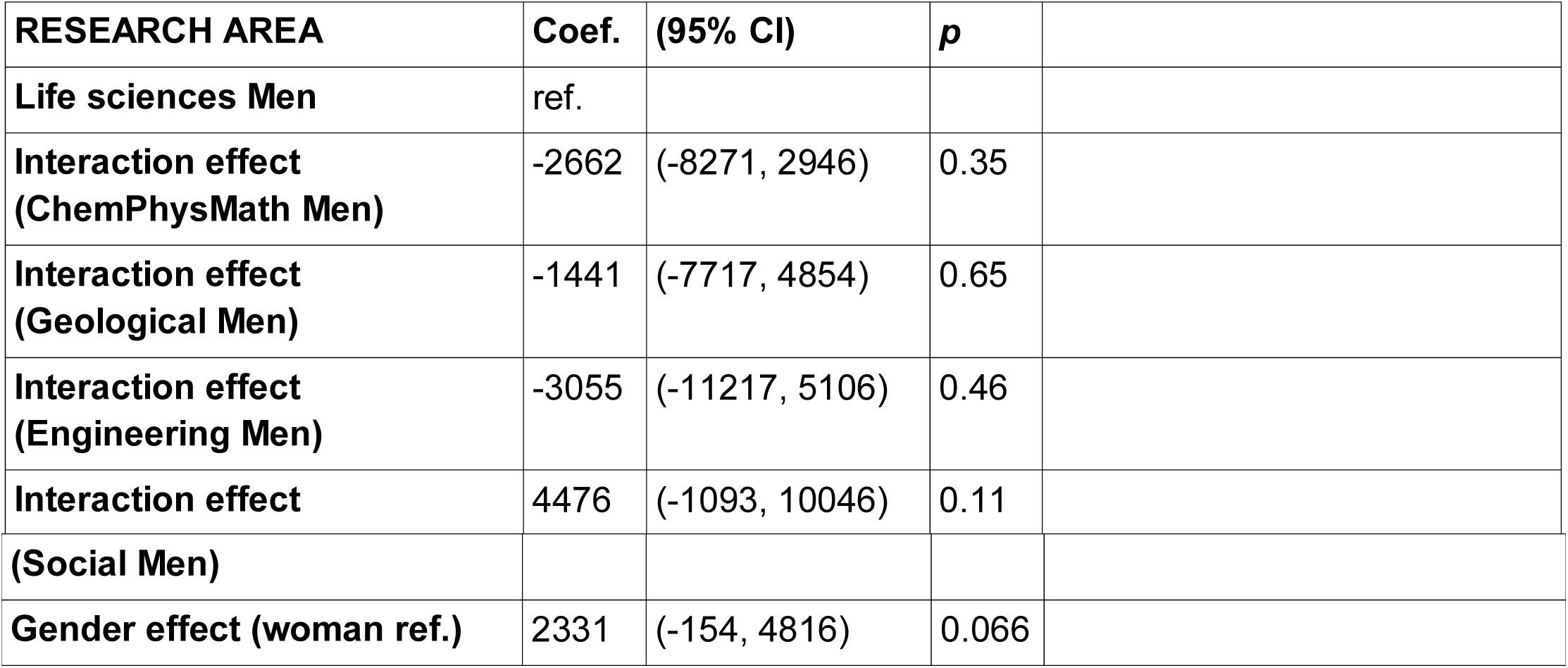
Interaction between Gender and Research Area (gross income)

No significant interaction effect was found between Research Area and gender.

**Table 6:** Interaction between Gender and Mobility (gross income)

| **MOBILITY FROM BIRTH COUNTRY** | **Coef.** | **(95% CI)** | **p** |
| --- | --- | --- | --- |
| **Non-mobile Men** | ref. |  |  |
| **Interaction effect (Mobile Men)** | 4518 | (-269, 9305) | 0.064 |
| **Interaction effect (Very mobile Men)** | 984 | (-3196, 5164) | 0.64 |
| **Gender effect (woman ref.)** | 1165 | (-1339, 3669) | 0.36 |

No significant interaction effect was found between mobility and gender, though with some indication that Mobile men earned more than non-mobile men. Interestingly, with this interaction modelled, the significant main effect of Gender disappeared.

#### Adjusted model for net income

We then run the same adjusted model for *net income*. Here, we found no significant effects of gender. The only variables where differences in net income were observed were Work Country and Country of Origin, where postdoctoral researchers originating from and working in Southern and Western regions earned less compared to their Western counterparts. 32 respondents had not included their net income, thus N = 614 for these analyses.

**Table 7:** Adjusted associations between gender and other independent variables of interest and net annual income in euros (N=612)

|  | **Adjusted** |  |  |
| --- | --- | --- | --- |
|  | **Coef.** | **(95% CI)** | **p** |
| **Gender** |  |  |  |
| Women | ref. |  |  |
| Men | 1254 | (-1867, 4374) | 0.43 |
| **Months since PhD** | 21 | (-26, 68) | 0.37 |
| **Children** |  |  |  |
| No | ref. |  |  |
| Yes | 1033 | (-2530, 4596) | 0.57 |
| **Age in years** | -645 | (-387, 372) | 0.97 |
| **Country of origin** |  |  |  |
| Western | ref. |  |  |
| Eastern | -5688 | (-13259, 1883) | 0.14 |
| Southern | -5524 | (-10901, -147) | 0.044 |
| **Country of working** |  |  |  |
| Western | ref. |  |  |
| Eastern | -19529 | (-30213, -8846) | <0.001 |
| Southern | -9822 | (-15142, -4502) | <0.001 |
| **Research area** |  |  |  |
| Chemistry, Physics, Mathematics | -98 | (-4847, 4650) | 0.97 |
| Geological Sciences | 2757 | (-2294, 7808) | 0.28 |
| Engineering Sciences | -903 | (-7125, 5319) | 0.78 |
| Life Sciences | ref. |  |  |
| Social Sciences | -1575 | (-6015, 2865) | 0.49 |
| **Mobility** |  |  |  |
| No | ref. |  |  |
| Within country of origin group | 661 | (-4236, 5558) | 0.79 |
| Outside country of origin group | 256 | (-5063, 5576) | 0.92 |

This indicates that whatever differences we found for the gross income between the genders must be interpreted carefully. It is possible that these differences may be explained by differences in type of income and whether this is taxed. For example, if women are likely to receive income from tax-free sources such as stipends, this may reflect why differences were observed in gross but not net income. This would suggest that there are no evident income inequalities by gender, however, if women are more likely to make their earnings from stipends while men earn taxable income, it might suggest that women have less social security and poorer employment conditions compared to men.

We tested the same four specific gender interaction, *children, work country, research area* and *mobility,* as these are likely to differentially impact on earning opportunities for men and women. This analysis is adjusted for all other covariates. Again no gender effects were found, even when these interactions were included. An interaction effect indicated though that mobile men earned more than mobile women. It should also be remembered that this interaction effect in the model, the main effect of gender disappeared for the gross income. If there are any gender difference, these apparently appear for mobile researchers, not for the genders as such.

**Table 8:** Interaction between Gender and Children (net income)

| **CHILDREN** | **Coef.** | **(95% CI)** | **p** |
| --- | --- | --- | --- |
| **Men without children** | ref. |  |  |
| **Interaction effect (Men with children)** | -4490 | (-11043, 2063) | 0.18 |
| **Gender effect (women=ref.)** | 2766 | (-1013, 6544) | 0.115 |

No significant interaction effect was found between Gender and Children.

**Table 9:** Interaction between Gender and Work Country (net income)

| **WORK COUNTRY** | **Coef.** | **(95% CI)** | **p** |
| --- | --- | --- | --- |
| **Western Men** | ref. |  |  |
| **Interaction effect (Eastern Men)** | -1769 | (-19416, 15877) | 0.84 |
| **Interaction effect (Southern Men)** | -1760 | (-7841, 4321) | 0.57 |
| **Gender effect (woman ref.)** | 2171 | (-2171, 6514) | 0.33 |

No significant interaction effect was found between Gender and Work Country.

**Table 10:** Interaction between Gender and Research Area (net income)

| **RESEARCH AREA** | **Coef.** | **(95% CI)** | **p** |
| --- | --- | --- | --- |
| **Life sciences Men** | ref. |  |  |
| **Interaction effect (ChemPhysMath Men)** | 243 | (-9319, 9783) | 0.96 |
| **Interaction effect (Geological Men)** | 7666 | (-2732, 18065) | 0.15 |
| **Interaction effect (Engineering Men)** | -2780 | (-16584, 11025) | 0.69 |
| **Interaction effect (Social Men)** | 4376 | (-5723, 12675) | 0.46 |
| **Gender effect (woman ref.)** | 101 | (-4016, 4219) | 0.96 |

No significant interaction effect was found between Research Area and gender.

**Table 11:**
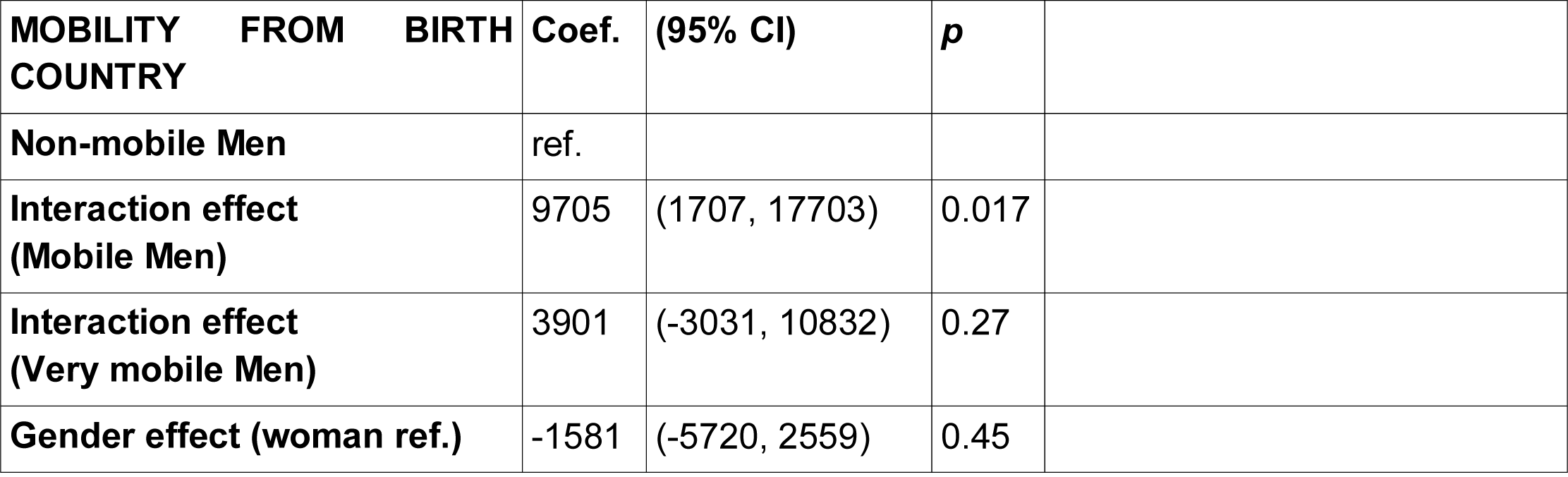
Interaction between Gender and Mobility (net income)

A signification interaction effect was found between Gender and Mobility showing that men that move within their country region report higher pays than women (Fig. 1).

**Figure 1.**
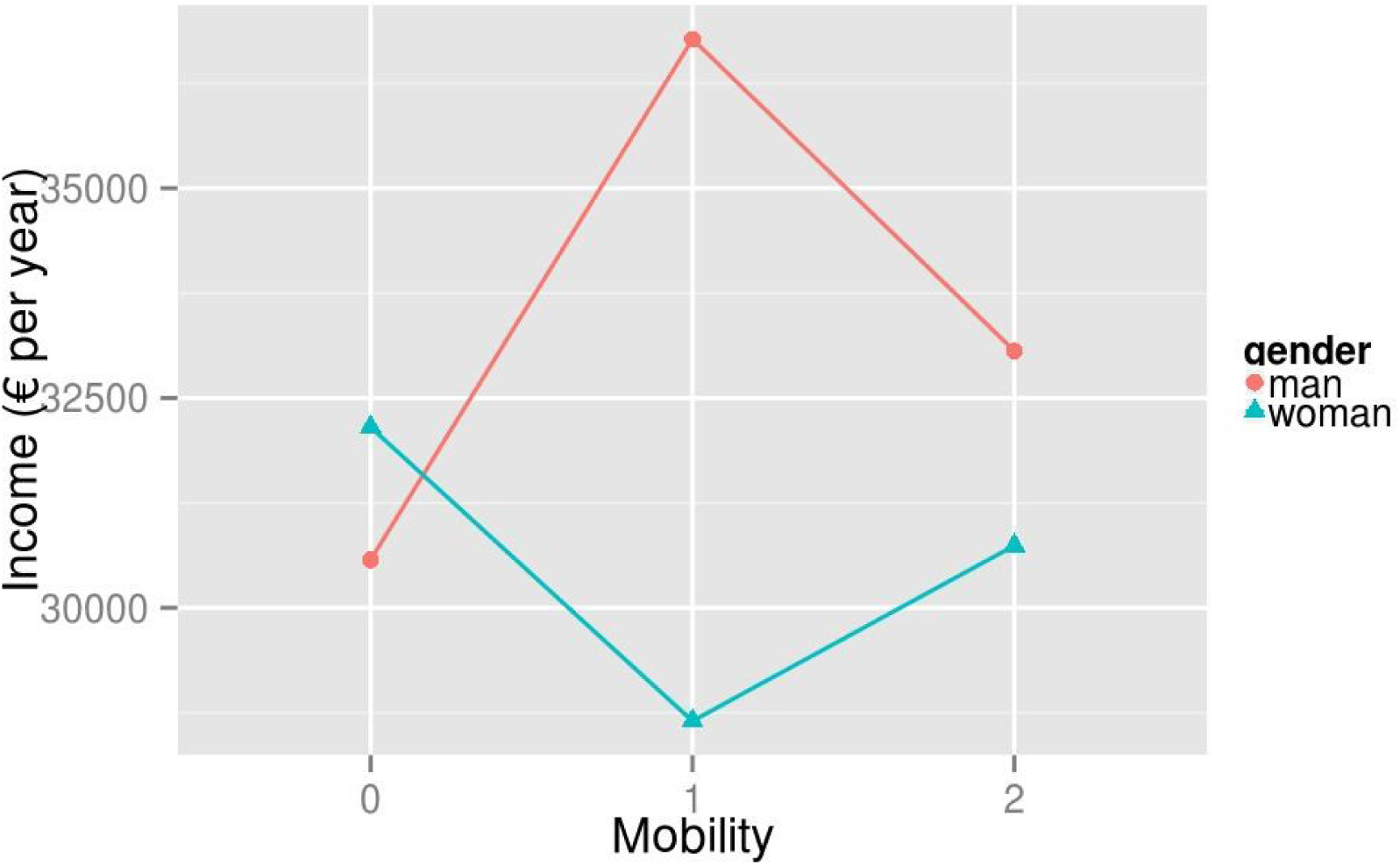
0=non-mobile, 1=mobile within group origin, 2=mobile outside group origin. The interaction is driven by mobile men within group origin earning more than women the same group

#### Conclusions

The conclusions that can be drawn from this survey regarding differences in pay between the genders will have to be tentative. A significant difference in yearly income was found in the gross income, but this effect did not hold up in the net income. A significant interaction between mobility and net income was found however (Table 11) (not found in the gross, even though the *p*-value was close to the significance level; Table 6). In terms of descriptive statistics, non-mobile women earned more than their male counterparts, whereas the pattern was reversed for the mobile researcher. The only significant difference found however was that among researchers who moved within their own nationality group (see definition in Further Processing), men earned more than women.

If any conclusion can be drawn, it is that mobile men may earn more than mobile women. No significant differences were found for the non-mobile researchers. As no significant difference between mobility and gender was found in the gross income, even this purported difference should be interpreted cautiously.

