## Supplementary material for "Postdoc X-ray in Europe 2017 Work conditions, productivity, institutional support and career outlooks": European survey for postdoctoral researchers

Dear postdoctoral fellow
We kindly invite you to contribute to this survey aimed at collecting information about the current work situation and career perspectives of postdoctoral researchers. The information will be used to raise awareness of the problems that concern researchers at this stage of their career.
Your participation in this research study is voluntary. In case you decide to participate, you may withdraw at any time and your responses will be eliminated. The survey will take approximately 15-20 minutes to complete.

Your responses will be confidential and we do not collect identifying information such as your name, email address or IP address. To help protect your confidentiality, the survey results will not contain information that will personally identify you.

The results of this study will only be used for scholarly purposes. They may be presented to the scientific community, included in scientific publications or deposited in relevant scientific database.

The survey questions will focus on your current postdoctoral situation, career opportunities and career development.

If you have any questions about the research study, please contact our team at

The survey was developed by the European Network of Postdoctoral Associations (https://www.uc.pt/en/iii/postdoc/ENPA).

*Obrigatório

#### Sem título

1. **By clicking YES, you state that you are aware of the aims of this study and voluntarily agree to participate in this survey.** *

   *Marcar apenas uma oval.*

   ◯ Yes

2. **ID code** *

   To identify eventual duplicated records, we kindly ask you to insert a code (last three letters of your last name and year of birth). E.g. John Smith, born in 27 March 1971, would insert ITH1971. This information will be uncoupled from the survey answers after elimination of duplicates.

   ______________________________

#### SOCIODEMOGRAPHIC INFORMATION

3. **Gender** *

   *Marcar apenas uma oval.*

   ◯ Male
   ◯ Female
   ◯ Prefer not to answer

4. **Age** *

5. **What is your nationality (country)?** *

*Marcar apenas uma oval.*

- Afghanistan
- Akrotiri
- Albania
- Algeria
- American Samoa
- Andorra
- Angola
- Anguilla
- Antarctica
- Antigua and Barbuda
- Argentina
- Armenia
- Aruba
- Ashmore and Cartier Islands
- Australia
- Austria
- Azerbaijan
- Bahamas, The
- Bahrain
- Bangladesh
- Barbados
- Bassas da India
- Belarus
- Belgium
- Belize
- Benin
- Bermuda
- Bhutan
- Bolivia
- Bosnia and Herzegovina
- Botswana
- Bouvet Island
- Brazil
- British Indian Ocean Territory
- British Virgin Islands
- Brunei
- Bulgaria
- Burkina Faso
- Burma
- Burundi
- Cambodia

- Cameroon
- Canada
- Cape Verde
- Cayman Islands
- Central African Republic
- Chad
- Chile
- China
- Christmas Island
- Clipperton Island
- Cocos (Keeling) Islands
- Colombia
- Comoros
- Congo, Democratic Republic of the
- Congo, Republic of the
- Cook Islands
- Coral Sea Islands
- Costa Rica
- Cote d'Ivoire
- Croatia
- Cuba
- Cyprus
- Czech Republic
- Denmark
- Dhekelia
- Djibouti
- Dominica
- Dominican Republic
- Ecuador
- Egypt
- El Salvador
- Equatorial Guinea
- Eritrea
- Estonia
- Ethiopia
- Europa Island
- Falkland Islands (Islas Malvinas)
- Faroe Islands
- Fiji
- Finland
- France
- French Guiana

- French Polynesia
- French Southern and Antarctic Lands
- Gabon
- Gambia, The
- Gaza Strip
- Georgia
- Germany
- Ghana
- Gibraltar
- Glorioso Islands
- Greece
- Greenland
- Grenada
- Guadeloupe
- Guam
- Guatemala
- Guernsey
- Guinea
- Guinea-Bissau
- Guyana
- Haiti
- Heard Island and McDonald Islands
- Holy See (Vatican City)
- Honduras
- Hong Kong
- Hungary
- Iceland
- India
- Indonesia
- Iran
- Iraq
- Ireland
- Isle of Man
- Israel
- Italy
- Jamaica
- Jan Mayen
- Japan
- Jersey
- Jordan
- Juan de Nova Island
- Kazakhstan

- Kenya
- Kiribati
- Korea, North
- Korea, South
- Kuwait
- Kyrgyzstan
- Laos
- Latvia
- Lebanon
- Lesotho
- Liberia
- Libya
- Liechtenstein
- Lithuania
- Luxembourg
- Macau
- Macedonia
- Madagascar
- Malawi
- Malaysia
- Maldives
- Mali
- Malta
- Marshall Islands
- Martinique
- Mauritania
- Mauritius
- Mayotte
- Mexico
- Micronesia, Federated States of
- Moldova
- Monaco
- Mongolia
- Montenegro
- Montserrat
- Morocco
- Mozambique
- Namibia
- Nauru
- Navassa Island
- Nepal
- Netherlands

- Netherlands Antilles
- New Caledonia
- New Zealand
- Nicaragua
- Niger
- Nigeria
- Niue
- Norfolk Island
- Northern Mariana Islands
- Norway
- Oman
- Pakistan
- Palau
- Panama
- Papua New Guinea
- Paracel Islands
- Paraguay
- Peru
- Philippines
- Pitcairn Islands
- Poland
- Portugal
- Puerto Rico
- Qatar
- Reunion
- Romania
- Russia
- Rwanda
- Saint Helena
- Saint Kitts and Nevis
- Saint Lucia
- Saint Pierre and Miquelon
- Saint Vincent and the Grenadines
- Samoa
- San Marino
- Sao Tome and Principe
- Saudi Arabia
- Senegal
- Serbia
- Seychelles
- Sierra Leone
- Singapore

- Slovakia
- Slovenia
- Solomon Islands
- Somalia
- South Africa
- South Georgia and the South Sandwich Islands
- Spain
- Spratly Islands
- Sri Lanka
- Sudan
- Suriname
- Svalbard
- Swaziland
- Sweden
- Switzerland
- Syria
- Taiwan
- Tajikistan
- Tanzania
- Thailand
- Timor-Leste
- Togo
- Tokelau
- Tonga
- Trinidad and Tobago
- Tromelin Island
- Tunisia
- Turkey
- Turkmenistan
- Turks and Caicos Islands
- Tuvalu
- Uganda
- Ukraine
- United Arab Emirates
- United Kingdom
- United States
- Uruguay
- Uzbekistan
- Vanuatu
- Venezuela
- Vietnam
- Virgin Islands

- Wake Island
- Wallis and Futuna
- West Bank
- Western Sahara
- Yemen
- Zambia
- Zimbabwe
- Prefer not to answer

6. **Do you speak the language of the country you work in?** *
*Marcar apenas uma oval.*

- Yes, it is my mothertongue
- Yes, fluently
- Yes, but with difficulty
- No
- Prefer not to answer

7. **Which is your marital status?** *
*Marcar apenas uma oval.*

- Single
- Married or cohabiting
- In a relationship but not cohabiting
- Divorced/Widowed
- Prefer not to answer

8. **If in a relationship, do you live in the same country as your partner?** *
*Marcar apenas uma oval.*

- Yes
- No
- I am not in a relationship
- Prefer not to answer

9. **Do you have children?** *
*Marcar apenas uma oval.*

- Yes
- No
- Prefer not to answer

#### CURRENT WORK INFORMATION

This section refers to your current work situation.

10. **In which country do you currently work?** *

*Marcar apenas uma oval.*

- Afghanistan
- Akrotiri
- Albania
- Algeria
- American Samoa
- Andorra
- Angola
- Anguilla
- Antarctica
- Antigua and Barbuda
- Argentina
- Armenia
- Aruba
- Ashmore and Cartier Islands
- Australia
- Austria
- Azerbaijan
- Bahamas, The
- Bahrain
- Bangladesh
- Barbados
- Bassas da India
- Belarus
- Belgium
- Belize
- Benin
- Bermuda
- Bhutan
- Bolivia
- Bosnia and Herzegovina
- Botswana
- Bouvet Island
- Brazil
- British Indian Ocean Territory
- British Virgin Islands
- Brunei
- Bulgaria
- Burkina Faso
- Burma
- Burundi
- Cambodia

- Cameroon
- Canada
- Cape Verde
- Cayman Islands
- Central African Republic
- Chad
- Chile
- China
- Christmas Island
- Clipperton Island
- Cocos (Keeling) Islands
- Colombia
- Comoros
- Congo, Democratic Republic of the
- Congo, Republic of the
- Cook Islands
- Coral Sea Islands
- Costa Rica
- Cote d'Ivoire
- Croatia
- Cuba
- Cyprus
- Czech Republic
- Denmark
- Dhekelia
- Djibouti
- Dominica
- Dominican Republic
- Ecuador
- Egypt
- El Salvador
- Equatorial Guinea
- Eritrea
- Estonia
- Ethiopia
- Europa Island
- Falkland Islands (Islas Malvinas)
- Faroe Islands
- Fiji
- Finland
- France
- French Guiana

- French Polynesia
- French Southern and Antarctic Lands
- Gabon
- Gambia, The
- Gaza Strip
- Georgia
- Germany
- Ghana
- Gibraltar
- Glorioso Islands
- Greece
- Greenland
- Grenada
- Guadeloupe
- Guam
- Guatemala
- Guernsey
- Guinea
- Guinea-Bissau
- Guyana
- Haiti
- Heard Island and McDonald Islands
- Holy See (Vatican City)
- Honduras
- Hong Kong
- Hungary
- Iceland
- India
- Indonesia
- Iran
- Iraq
- Ireland
- Isle of Man
- Israel
- Italy
- Jamaica
- Jan Mayen
- Japan
- Jersey
- Jordan
- Juan de Nova Island
- Kazakhstan

- Kenya
- Kiribati
- Korea, North
- Korea, South
- Kuwait
- Kyrgyzstan
- Laos
- Latvia
- Lebanon
- Lesotho
- Liberia
- Libya
- Liechtenstein
- Lithuania
- Luxembourg
- Macau
- Macedonia
- Madagascar
- Malawi
- Malaysia
- Maldives
- Mali
- Malta
- Marshall Islands
- Martinique
- Mauritania
- Mauritius
- Mayotte
- Mexico
- Micronesia, Federated States of
- Moldova
- Monaco
- Mongolia
- Montenegro
- Montserrat
- Morocco
- Mozambique
- Namibia
- Nauru
- Navassa Island
- Nepal
- Netherlands

- Netherlands Antilles
- New Caledonia
- New Zealand
- Nicaragua
- Niger
- Nigeria
- Niue
- Norfolk Island
- Northern Mariana Islands
- Norway
- Oman
- Pakistan
- Palau
- Panama
- Papua New Guinea
- Paracel Islands
- Paraguay
- Peru
- Philippines
- Pitcairn Islands
- Poland
- Portugal
- Puerto Rico
- Qatar
- Reunion
- Romania
- Russia
- Rwanda
- Saint Helena
- Saint Kitts and Nevis
- Saint Lucia
- Saint Pierre and Miquelon
- Saint Vincent and the Grenadines
- Samoa
- San Marino
- Sao Tome and Principe
- Saudi Arabia
- Senegal
- Serbia
- Seychelles
- Sierra Leone
- Singapore

- Slovakia
- Slovenia
- Solomon Islands
- Somalia
- South Africa
- South Georgia and the South Sandwich Islands
- Spain
- Spratly Islands
- Sri Lanka
- Sudan
- Suriname
- Svalbard
- Swaziland
- Sweden
- Switzerland
- Syria
- Taiwan
- Tajikistan
- Tanzania
- Thailand
- Timor-Leste
- Togo
- Tokelau
- Tonga
- Trinidad and Tobago
- Tromelin Island
- Tunisia
- Turkey
- Turkmenistan
- Turks and Caicos Islands
- Tuvalu
- Uganda
- Ukraine
- United Arab Emirates
- United Kingdom
- United States
- Uruguay
- Uzbekistan
- Vanuatu
- Venezuela
- Vietnam
- Virgin Islands

- Wake Island
- Wallis and Futuna
- West Bank
- Western Sahara
- Yemen
- Zambia
- Zimbabwe
- Prefer not to answer

11. **In which University or Institution do you currently work?** *

Please provide the name of the University/Institution in English (e.g. University of Uppsala). Do not use abbreviations. Please do not include the department.

12. **What is your primary reseach area** *

*Marcar apenas uma oval.*

- CHEMISTRY
- ECONOMICS, FINANCE AND MANAGEMENT
- INFORM SCIENCE AND ENGINEERING | Computer science and informatics
- INFORM SCIENCE AND ENGINEERING | | Systems and communication engineering
- INFORM SCIENCE AND ENGINEERING | Products and process engineering
- ENVIRON AND GEOSCIENCES | Environment and society
- ENVIRON AND GEOSCIENCES | Earth system science
- ENVIRON AND GEOSCIENCES | Evolutionary, population and environmental biology
- ENVIRON AND GEOSCIENCES | Agricultural, animal, fishery, forestry and food science
- LIFE SCIENCES | Molecular and Structural Biology and Biochemistry
- LIFE SCIENCES | Genetics, Genomics, Bioinformatics and Systems Biology
- LIFE SCIENCES | Cellular and Developmental Biology
- LIFE SCIENCES | Physiology, Pathophysiology and Endocrinology
- LIFE SCIENCES | Neurosciences and neural disorders
- LIFE SCIENCES | Immunity and infection
- LIFE SCIENCES | Diagnostic tools, therapies and public health
- LIFE SCIENCES | Applied life sciences
- MATHEMATICS
- PHYSICS | Fundamental constituents of matter
- PHYSICS | Condensed matter physics
- PHYSICS | Universe sciences
- SOC SCIENCES AND HUMANITIES | Sociology, social anthropology, political science, law, communication
- SOC SCIENCES AND HUMANITIES | Cognition, psychology, linguistics, philosophy and education
- SOC SCIENCES AND HUMANITIES | Literature, arts, music, cultural and comparative studies
- SOC SCIENCES AND HUMANITIES | Archaeology, history and memory

13. **What is your secondary research area?**
Below you can select another area of research. Please leave blank if you do not have another area of research that you would like to mention.
*Marcar apenas uma oval.*

- CHEMISTRY
- ECONOMICS, FINANCE AND MANAGEMENT
- INFORM SCIENCE AND ENGINEERING | Computer science and informatics
- INFORM SCIENCE AND ENGINEERING | | Systems and communication engineering
- INFORM SCIENCE AND ENGINEERING | Products and process engineering
- ENVIRON AND GEOSCIENCES | Environment and society
- ENVIRON AND GEOSCIENCES | Earth system science
- ENVIRON AND GEOSCIENCES | Evolutionary, population and environmental biology
- ENVIRON AND GEOSCIENCES | Agricultural, animal, fishery, forestry and food science
- LIFE SCIENCES | Molecular and Structural Biology and Biochemistry
- LIFE SCIENCES | Genetics, Genomics, Bioinformatics and Systems Biology
- LIFE SCIENCES | Cellular and Developmental Biology
- LIFE SCIENCES | Physiology, Pathophysiology and Endocrinology
- LIFE SCIENCES | Neurosciences and neural disorders
- LIFE SCIENCES | Immunity and infection
- LIFE SCIENCES | Diagnostic tools, therapies and public health
- LIFE SCIENCES | Applied life sciences
- MATHEMATICS
- PHYSICS | Fundamental constituents of matter
- PHYSICS | Condensed matter physics
- PHYSICS | Universe sciences
- SOC SCIENCES AND HUMANITIES | Sociology, social anthropology, political science, law, communication
- SOC SCIENCES AND HUMANITIES | Cognition, psychology, linguistics, philosophy and education
- SOC SCIENCES AND HUMANITIES | Literature, arts, music, cultural and comparative studies
- SOC SCIENCES AND HUMANITIES | Archaeology, history and memory

#### RESEARCH SITUATION AND CAREER

14. **Year of PhD conclusion** *
Please include just the year in which you have been awarded your PhD (Year of the PhD diploma)

15. **Did you obtain your PhD at your current institution** *
*Marcar apenas uma oval.*

- Yes
- No, at another European institution
- No, at an non-European institution
- Prefer not to answer

16. **Since your PhD, how long (in months) have you been working as a researcher?** *
Please provide your reply in months.

_______________________

17. **Did you have another postdoc position before you started your current contract/fellowship?** *
*Marcar apenas uma oval.*

- No
- Yes, in another country
- Yes, in the same country but in a different department/institution
- Yes, in the same department/institution
- Prefer not to answer

18. **What is the nature of your current fellowship /contract?** *
*Marcar apenas uma oval.*

- Fixed term
- Open ended (can be known as permanent)
- Casual/hourly paid
- Not sure
- Prefer not to answer

19. **Your postdoctoral position is funded by** *
*Marcar apenas uma oval.*

- Your own funding (e.g. research fellowship like Marie Curie Grant)
- A research grant (awarded to a project)
- Your institution (core funding)
- Prefer not to answer
- Outra: _______________________

20. **Is your contract/fellowship** *
*Marcar apenas uma oval.*

- Full-time with exclusivity clause (not allowed to have another income)
- Full-time without exclusivity clause (allowed to have another income)
- Part-time (by your own choice)
- Part-time (it was the only available option)
- I don't know
- Prefer not to answer

21. **For how many hours per week were you contracted?** *
Insert 0 (zero) if you don't have this specification

22. **On average, how many hours do you work per week?** *

23. **Average annual income (in Euros) [Gross income]**
Please, if your income is not in euros, use a currency converter (e.g. http://www.xe.com/currencyconverter/)

24. **Average annual income (in Euros) [Net income]**
Please, if your income is not in euros, use a currency converter (e.g. http://www.xe.com/currencyconverter/)

25. **Does your contract/fellowship include provision for:** *
*Marcar apenas uma oval por linha.*

| | Yes | No | I don't know | Not applicable |
|---|---|---|---|---|
| Parental leave | | | | |
| Access to healthcare | | | | |
| Unemployment benefit | | | | |
| Sick leave | | | | |

### TEACHING AND RESEARCH ACTIVITIES

26. **Do you contribute to any teaching (e.g. classroom or small group teaching) ?** *
*Marcar apenas uma oval.*

- No *Passe para a pergunta 28.*
- Yes, undergraduate *Passe para a pergunta 27.*
- Yes, graduate *Passe para a pergunta 27.*
- Yes, both *Passe para a pergunta 27.*
- Prefer not to answer

27. **Are you obliged to teach?** *
*Marcar apenas uma oval.*

- Yes
- No
- Prefer not to answer

*Passe para a pergunta 29.*

28. **Would you like to teach but are not given the opportunity?** *

*Marcar apenas uma oval.*

- Yes
- No
- Prefer not to answer

29. **Regarding master students' supervision** *

*Marcar apenas uma oval.*

- I am allowed to supervise and co-supervise
- I am allowed to co-supervise but not to be the main supervisor
- I supervise, but my supervision is not formally acknowledged
- I am not allowed to do any type of supervision
- I do not want to supervise
- I don't know
- I prefer not to say

30. **Regarding doctoral students' supervision** *

*Marcar apenas uma oval.*

- I am allowed to supervise and co-supervise
- I am allowed to co-supervise but not to be the main supervisor
- I supervise, but my supervision is not formally acknowledged
- I am not allowed to do any type of supervision
- I don't know
- I prefer not to say

31. **Are you allowed to apply for funding as a lead investigator of a research project?** *

*Marcar apenas uma oval.*

- Yes
- No
- I don't know
- Prefer not to answer

32. **Do you collaborate with your supervisor in grant writing?** *

*Marcar apenas uma oval.*

- Yes
- No
- Prefer not to answer

#### RESEARCHER PRODUCTIVITY

Please refer to publications where you are author or co-author in any position.

33. **How many papers in peer reviewed journals have you published (already published or in press)?** *
Please refer to publications where you are author or co-author in any position.

34. **How many books have you published?** *
Please refer to publications where you are author or co-author in any position.

35. **How many book chapters have you published?** *
Please refer to publications where you are author or co-author in any position.

36. **How many preprints have you published in preprint archives?** *

37. **Which is your h-index?**
The h index is an author-level metric. You can obtain your h-index at scopus, researcher ID or google scholar. This question is not compulsory.

38. **Which method did you use to calculate the h-index?**
*Marcar apenas uma oval.*

- Scopus
- Researcher ID
- Google Scholar
- Outra: ____

39. **Are you acknowledged as first author in the publications for which you have done the most significant contributions?** *
*Marcar apenas uma oval.*

- Yes, always
- Yes, but just in some situations
- No
- Prefer not to answer

40. **Does your group have clear rules on authorship issues, or an authorship policy?** *
*Marcar apenas uma oval.*

- Yes
- No
- I do not know
- Prefer not to answer

41. **Does your group have clear rules on scientific misconduct?** *
*Marcar apenas uma oval.*

- Yes
- No
- I do not know
- Prefer not to answer

42. **Have there been instances in which you haven´t agreed with the authorship order in a paper to which you contributed?**
*Marcar apenas uma oval.*

- Yes
- No
- Prefer not to answer

### INSTITUTIONAL SUPPORT

43. **How do you rate the quality of the facilities at your institution [e.g. work space, equipment]** *
*Marcar apenas uma oval.*

| | 1 | 2 | 3 | 4 | |
|---|---|---|---|---|---|
| Not at all satisfied | | | | | Very much satisfied |

44. **How do you rate the quality of the institutional support to your research [e.g. support in project management, technical support, grant writing]?** *
*Marcar apenas uma oval.*

| | 1 | 2 | 3 | 4 | |
|---|---|---|---|---|---|
| Not at all satisfied | | | | | Very much satisfied |

45. **How do you rate your satisfaction with access to funding for research activities [other than the scholarship; e.g. funding for presenting your work at conferences, training activities]?** *
*Marcar apenas uma oval.*

| | 1 | 2 | 3 | 4 | |
|---|---|---|---|---|---|
| Not all satisfied | | | | | Very much satisfied |

46. **How well established were your duties and rights at the beginning of your contract/fellowship [e.g. working hours, leave/holidays, external collaborations]?** *
*Marcar apenas uma oval.*

- Clearly defined rules
- Partially defined rules
- Unclear rules
- I don't know
- Prefer not to answer

47. **Are postdoctoral researchers represented in the management bodies of your University/Institution?** *
*Marcar apenas uma oval.*

- Yes
- No
- I don't know
- Prefer not to answer

#### CAREER DEVELOPMENT

48. **When thinking of your future career, in which role would you like to work in the long run?** *
*Marcar apenas uma oval.*

- Outside academia, in a researcher role
- Outside academia, in a non-researcher role
- In academia, as a professor/researcher
- In academia, as a researcher only
- In academia, teaching only
- In academia, in science communication or research management
- In scientific publishing
- Not yet decided
- Prefer not to answer
- Outra: ______________________

49. **How clear is your career plan?**
*Marcar apenas uma oval.*

| | 1 | 2 | 3 | 4 | 5 | |
|---|---|---|---|---|---|---|
| Not at all clear | ◯ | ◯ | ◯ | ◯ | ◯ | Very clear |

50. **Are you currently searching for jobs?** *
*Marcar apenas uma oval.*

- Yes, in academia
- Yes, outside academia
- Yes, both
- No
- Prefer not to answer

#### CAREER DEVELOPMENT SUPPORT

51. **Does your institution have an office to support postdoctoral researchers?**
*Marcar apenas uma oval.*

- Yes *Passe para a pergunta 52.*
- No *Passe para a pergunta 53.*
- I don't know *Passe para a pergunta 53.*

52. **How satisfied are you with the support provided by your postdoctoral researchers' office?**
*Marcar apenas uma oval.*

| | 1 | 2 | 3 | 4 | 5 | |
|---|---|---|---|---|---|---|
| Not satisfied at all | | | | | | Very much satisfied |

53. **To what extent do you agree that the institution where you work:** *
*Marcar apenas uma oval por linha.*

| | Not appplicable | I don´t know | Strongly disagree | Disagree | Agree | Strongly agree |
|---|---|---|---|---|---|---|
| provides assistance and advice about career opportunities | | | | | | |
| offers necessary transferable skills training | | | | | | |
| encourages you to engage in personal and career development | | | | | | |
| provides assistance with work related conflict resolution | | | | | | |
| provides a mentorship program | | | | | | |

54. **In which areas have you undertaken training?** *
*Marcar apenas uma oval por linha.*

| | Undertaken | Not undertaken, but I would like to | This is of no interest to me currently |
|---|---|---|---|
| Writing grant applications | | | |
| Career management | | | |
| Collaboration and teamworking | | | |
| Communication and dissemination | | | |
| Equality and diversity | | | |
| Ethical research conduct | | | |
| Interdisciplinary research | | | |
| Knowledge transfer and entrepreneurship | | | |
| Leadership and management | | | |
| Personal productivity and organisation | | | |
| Public engagement | | | |
| Research impact | | | |
| Supervision of doctoral and master students | | | |
| Teaching or lecturing | | | |

### POSTDOCTORAL COMMUNITY ENGAGEMENT

55. **Are you a member of a union?** *
A union is an organization that represents the collective interests of workers, and that usually negotiates with employers and governments over wages, hours, benefits and working conditions
*Marcar apenas uma oval.*

- No
- Yes
- I don't know
- Prefer not to answer

56. **Does your institution have a postdoctoral association?** *
*Marcar apenas uma oval.*

- Yes *Passe para a pergunta 57.*
- No *Passe para a pergunta 59.*
- I don't know *Passe para a pergunta 59.*
- Prefer not to answer

57. **Are you a member of a postdoctoral association?** *
*Marcar apenas uma oval.*

- No
- Yes
- Prefer not to answer

58. **How did you get to know of the postdoctoral association at your institution?** *
*Marcar apenas uma oval.*

- Through your PI
- Institution
- Fellow postdoctoral researchers
- Induction events
- Internet search
- Prefer not to answer
- Outra: ______________

59. **In your opinion, which of these events should a postdoctoral association organise?** *
*Marcar apenas uma oval por linha.*

| | We already have these events | We don't have these events but would like to | I am not interested in this |
|---|---|---|---|
| Mentoring sheme | | | |
| Someone other than your PI that helps with your career development | | | |
| Scientific networking events | | | |
| Social networking events | | | |
| Workshops/seminars/discussion panels on career paths | | | |

60. **Do you have any other suggestions concerning the postdoctoral associations?**
Please include these in the box below.

#### Other comments and suggestions

61. **Comments and suggestions**
Please leave any comments or suggestions in the box below regarding any topic related to this survey.

Com tecnologia
Google Forms
